# Structural Basis of βKNL2 Centromeric Targeting Mechanism and Its Role in Plant-Specific Kinetochore Assembly

**DOI:** 10.1101/2024.07.30.605747

**Authors:** Ramakrishna Yadala, Amanda S Camara, Surya P Yalagapati, Pascal Jaroschinsky, Tobias Meitzel, Mariko Ariyoshi, Tatsuo Fukagawa, Twan Rutten, Thu-Giang T Bui, Inna Lermontova

## Abstract

The kinetochore is an essential protein complex that ensures proper chromosome segregation during cell division. Kinetochore assembly is initiated by the incorporation of CENP-A/CENH3. This process depends on KNL2/M18BP1 and CENP-C proteins. In plants, two variants of KNL2, αKNL2 and βKNL2, are present. Both possess the conserved SANTA domain, while αKNL2 additionally has the centromere-targeting CENPC-k motif. Despite lacking the CENPC-like motif, the plant-specific βKNL2 localizes to centromeres and aids in CENP-A/CENH3 loading. We found that efficient centromeric targeting of βKNL2 requires the SANTA domain and the C-terminal part, while nuclear targeting depends on a conserved C-terminal motif-III. Structural predictions and experimental validations reveal that βKNL2 forms homodimers and interacts with centromeric DNA and αKNL2. We confirm that centromeric targeting of βKNL2 depends on αKNL2 in a tissue-dependent manner. Our findings provide crucial insights into the unique mechanisms of plant-specific kinetochore assembly, highlighting βKNL2’s essential role in this process.

**Graphical abstract:** 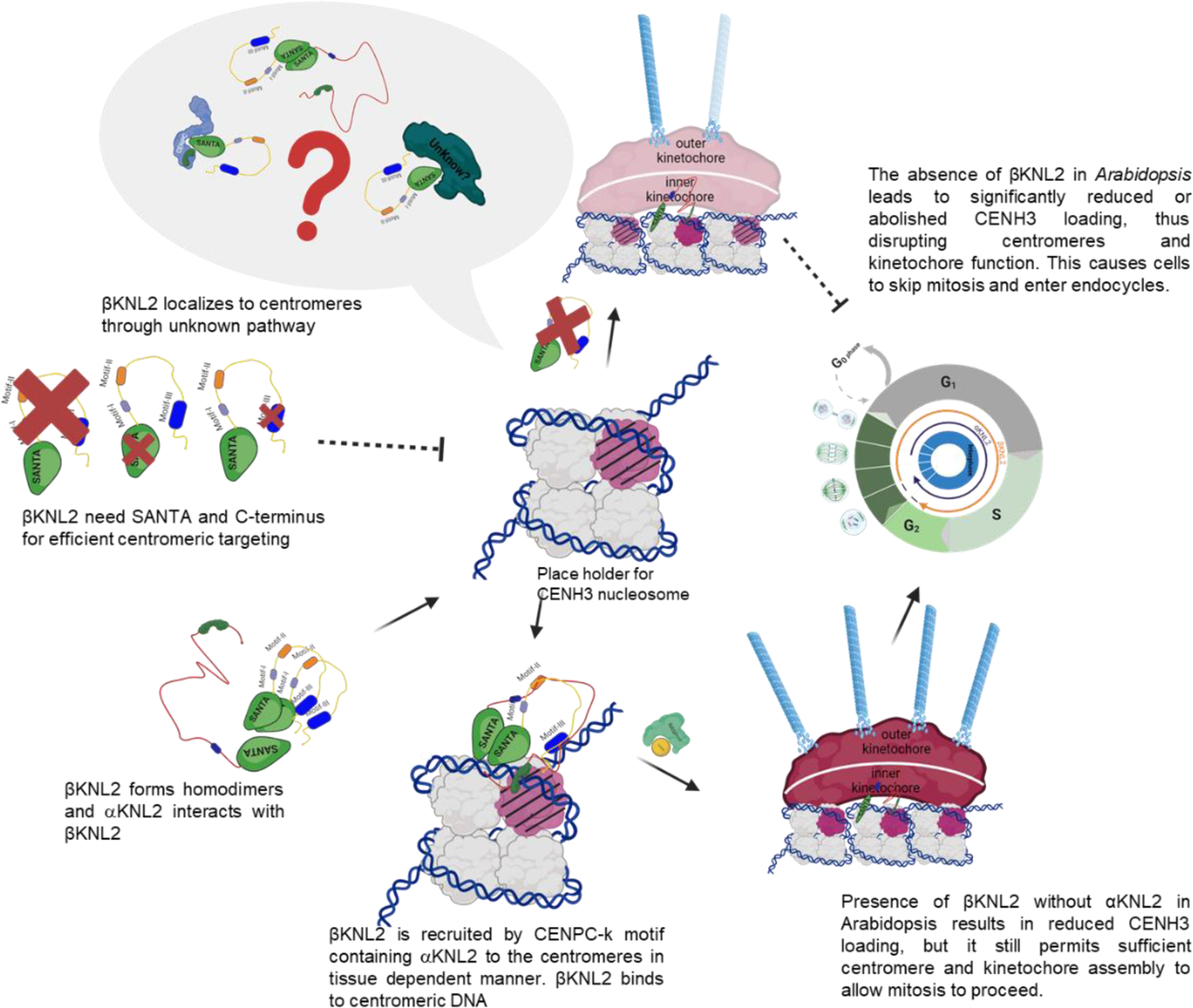

## Introduction

Kinetochores are large protein complexes that assemble on the centromere of each chromosome and connect the chromatids to mitotic spindles. The kinetochore consists of an inner kinetochore module directly attached to centromeric chromatin and an outer module connects to microtubules. (Cheeseman and Desai 2008; Cheeseman 2014; Yamagishi, et al. 2014; Musacchio and Desai 2017; Hara and Fukagawa 2018). Loss of kinetochore function causes chromosome mis-segregation, aneuploidy and cell death (Fachinetti, et al. 2013; McKinley and Cheeseman 2014; Barra and Fachinetti 2018). Centromere specific Histone3 (CENP-A/CENH3) nucleosomes trigger the assembly of a functional kinetochore (Talbert, et al. 2002). CENP-A/CENH3 loading to centromeres involves initiation, deposition, and maintenance stages (De Rop, et al. 2012).

KINETOCHORE NULL2 (KNL2, also termed M18BP1) plays a key role in new CENP-A/CENH3 deposition (Maddox, et al. 2007; Moree, et al. 2011; Lermontova, et al. 2013). In vertebrates, M18BP1/KNL2 is part of the Mis18 complex, which is essential for assembling and maintaining centromeres and kinetochore during cell division (Hayashi, et al. 2004; Fujita, et al. 2007; Maddox, et al. 2007). The Mis18 is an oligomeric structure with multiple copies of its subunits (Mis18α, Mis18β, and M18BP1/KNL2) (Barnhart-Dailey and Foltz 2014; Smith and Maddox 2017). CDK1 regulates Mis18 complex recruitment to centromeres by controlling M18BP1/KNL2 oligomerization (Pan, et al. 2017). In plants Mis18α and Mis18β have not yet been identified. In chicken and *Xenopus* the M18BP1/KNL2 protein is present at centromeres throughout the cell cycle (French, et al. 2017; Hori, et al. 2017) and in plants as well except from metaphase to mid-anaphase (Lermontova, et al. 2013).

All identified M18BP1/KNL2 proteins typically feature the SANT-associated domain (SANTA), which is about 90 amino acids (AA) long (Zhang, et al. 2006; Stellfox, et al. 2013). This domain is often accompanied by a SANT domain and/or a CENPC-like motif (Figure 1A). The functional significance of the SANTA remains under debate. Deletion of the SANTA containing part in *Arabidopsis* αKNL2 did not impair its centromere targeting (Lermontova, et al. 2013) nor disrupted its interaction with DNA (Sandmann, et al. 2017). The SANTA of *Xenopus* M18BP1/KNL2 mediates its interaction with CENP-C during the metaphase, thus regulating its centromere targeting (French and Straight 2019). However, M18BP1/KNL2 interaction with CENP-A/CENH3 nucleosomes in *Xenopus* and chicken does not require the SANTA (French, et al. 2017; Jiang, et al. 2022). Whereas modification of the CENPC-like motif in *Arabidopsis*, chicken and *Xenopus* led to mis-localization of KNL2 (French, et al. 2017; Hori, et al. 2017; Sandmann, et al. 2017). In contrast to this, plant-specific βKNL2 localizes to centromeres and functions as a CENP-A/CENH3 assembly factor despite lacking a CENPC-like motif (Zuo, et al. 2022). The underlying mechanisms of βKNL2 centromere targeting remain unclear and hinders our understanding of centromere and kinetochore assembly.

**Figure. 1.**
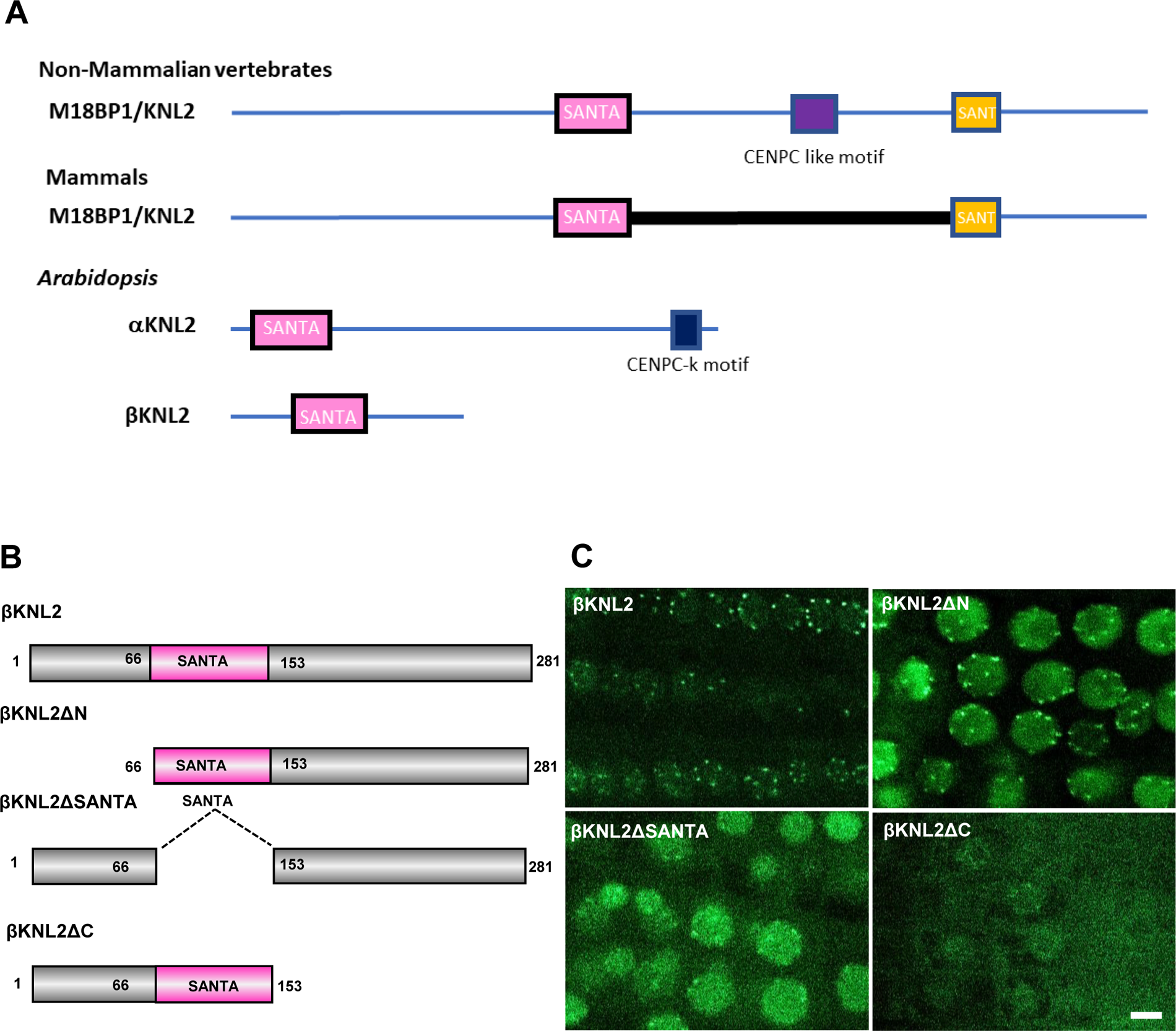
SANTA domain and C-terminus of βKNL2 regulate its centromeric targeting. (A) Domain organization of M18BP1/KNL2 in different species. M18BP1/KNL2 in non-mammalian vertebrates includes SANTA (magenta) and SANT (yellow) domains and a CENPC-like motif (purple), while in mammals it includes SANTA and SANT domains separated by the CENP-C binding region (thick black line). *Arabidopsis* αKNL2 contains the SANTA domain and CENPC-k motif (blue), while βKNL2 is characterized solely by the SANTA domain. (B) Schemata of βKNL2 truncated variants created by site-directed mutagenesis: βKNL2ΔN lacks the N-terminus; βKNL2ΔSANTA lacks the SANTA domain; βKNL2ΔC lacks the C-terminus. (C) Localization patterns of EYFP-tagged βKNL2 truncated variants (referred in 1B) in root tip nuclei of 7-day-old *A. thaliana* seedlings. The deletion of the SANTA domain and C-terminus affect the centromeric localization, underscoring their critical roles in targeting βKNL2 to the centromere. Scale bar 5µm

Unlike *αknl2* mutants *βknl2* exhibits embryo lethal phenotype in *Arabidopsis* (Lermontova, et al. 2013; Zuo, et al. 2022). We hypothesize that βKNL2 has a crucial role in centromere and kinetochore assembly. To fill the knowledge gap in plant kinetochore assembly, the present study focuses on four open questions: i) how is βKNL2 targeted to centromeres despite lacking CENPC-like motif: which domains/motifs regulate its centromeric targeting; ii) what is the position of plant specific KNL2 in the kinetochore (inner or outer kinetochore); iii) how do these key components bind to centromeric nucleosomes and iv) does centromeric targeting of βKNL2 depend on other proteins? To answer these questions, we used Alphafold2 to build centromeric nucleosome in complex with KNL2 proteins and dissected βKNL2 to identify functional motifs and domains necessary for centromere targeting and interactions with kinetochore proteins.

## Materials and methods

### Plasmid construction, plant transformation, and plant growth conditions

For the localization studies, various constructs (βKNL2ΔSANTA, βKNL2ΔN, βKNL2ΔC, βKNL2(C), βKNL2Δmotif-I, βKNL2Δmotif-II, βKNL2Δmotif-III, βKNL2Δmotif-III new) were generated through site-directed mutagenesis using the βKNL2 cDNA cloned in pDONR221, as previously described inZuo, et al. (2022). The mutagenesis was carried out following the manufacturer’s protocol with the Phusion site-directed mutagenesis kit from Thermo Fisher Scientific. Entry plasmids were then subcloned into Gateway-compatible expression pGWB641via LR reaction (Invitrogen). The endogenous βKNL2 was amplified from Col0 genomic DNA and cloned into pDONR221 via BP reaction, followed by subcloning into pGWB640 through LR reaction.

For PPI studies, constructs of βKNL2, βKNL2ΔSANTA, αKNL2(C), and CENP-A/CENH3 were subcloned into 3’Ven-N-pBAR GW and 3’Ven-C-pBAR GW vectors by LR reaction. This GATEWAY-compatible BIFC vectors generated based on the highly efficient and small binary vector pPZP200BAR (Jozefkowicz, et al. 2016). To this end, the cauliflower mosaic virus 35S promoter (35S) was assembled with the terminator sequence of the octopine synthase gene (tocs) from *Agrobacterium tumefaciens* via overlap extension PCR. The overlap primer were designed to create a 5’-*EcoR*I-*Spe*I-*Xba*I-3’ multiple cloning site (MCS) between the promoter and terminator sequences, and the resulting 35S:MCS:tocs fusion product was additionally flanked with restriction sites for *BamH*I and *Pst*I, After restriction enzyme digestion using *BamH*I/*Pst*I the 35S:MCS:tocs cassette was ligated into the accordingly linearized pPZP200BAR binary vector. To facilitate PPI analyses with the Venus fluorescent protein in all possible orientations (Waadt, et al. 2008), the coding sequences of both the VN (residues 1-173) and VC (residues 156-239) of Venus were amplified by PCR using primers suitable to create *EcoR*I and *Spe*I restriction sites for N-terminal fusions or *Spe*I and *Xba*I restriction sites for C-terminal fusions.

Additionally, a c-myc epitope tag was attached to both Ven_N_ variants and a hemagglutinin (HA) epitope tag to all Ven_C_ variants. The PCR fragments of the truncated Venus variants were cloned into the previously assembled binary 35S:MCS:ocs-pPZP200BAR vector using the aforementioned restriction sites. To modify the resulting BiFC binary vectors into GATEWAY-compatible destination vectors, a *Spe*I flanked PCR-fragment containing a GATEWAY *ccdB* gene cassette for LR was ligated into the linearized plasmids through digestion with *Spe*I. The final set of GATEWAY-compatible BIFC destination vectors, including 3’Ven-N-pBAR GW and 3’Ven-C-pBAR GW, was verified by sequencing. a schematic map of the GATEWAY-compatible BIFC vectors is given in Supplementary Figure S1

Genomic βKNL2 fragments were transformed into *αknl2* and *βknl2* mutants. In the case of wheat germ expression, βKNL2 was amplified with primers including a FLAG tag from the βKNL2 cDNA cloned in pDONR221 (Zuo, et al. 2022). The amplified product was then cloned into the pF3A WG (Promega) vector through restriction cloning using PmeI and SgfI restriction enzymes. All the clones were verified at each step through colony PCR using attB primers (Supplementary Table.1) and further confirmed by Sanger sequencing.

*Agrobacterium*-mediated transient expressions were conducted in tobacco plants following the protocol outlined by Yadala, et al. (2022). Stable transformants of *Arabidopsis* accession Columbia-0 were generated using the floral dip method (Clough and Bent 1998). Transformants were selected on Murashige and Skoog (MS) medium containing 20 mg/L of phosphinotricine under long-day conditions (16 h light/8 h dark). *Arabidopsis* plants were initially germinated under short-day conditions (8 h light/16 h dark) and, two weeks after transplanting, grown under long-day conditions (16 h light/8 h dark).

### Electrophoretic mobility shift assay

FLAG-βKNL2 was expressed using the TnT SP6 High-Yield Wheat Germ Protein Expression System following the manufacturer’s instructions (https://www.licor.com/bio/reagents/odyssey-emsa-kit). The expression reaction was incubated at 25°C for 2 hours. The expressed protein was confirmed by Western blot analysis using anti-FLAG-tag antibodies from Sigma. The centromeric repeat *pAL1* was amplified from *Arabidopsis* Col-0 genomic DNA using IRD700 labeled and unlabeled oligos, serving as the probe and competitor, respectively. Amplicons were purified using the oligonucleotide purification kit from BioRad.

The binding reaction was set up using the Odyssey EMSA kit (LICOR) with a protocol adapted fromEysholdt-Derzsó, et al. (2023). Motif-III extended peptide from brassica species (Lifetain) was included in the reaction for peptide DNA binding ability testing. The expressed FLAG-βKNL2 and motif-III extended peptides with the *pAL1* labeled probe was incubated for 30 minutes at room temperature. The reaction was then loaded onto a 5% native polyacrylamide gel and run at 4°C with a voltage of 70 V until the dye reached the bottom. Gel images were captured using a LICOR Odyssey scanner.

### Bimolecular Fluorescence Complementation (BIFC)

The BIFC assay was conducted following Yadala, et al. (2022). Overnight cultures of *Agrobacterium tumefaciens* strain GV3101 carrying the desired constructs were harvested and reconstituted in AIM medium. The cultures were adjusted to 0.8 OD, and cultures of the testing partners were combined in equal proportions, followed by incubation for 1 hour at room temperature. The abaxial side of leaves from 2-3-week-old *N. benthamiana* plants was infiltrated with the mixture using a 1ml syringe. Fluorescence was observed 48 hours post-infiltration under a Zeiss observer fluorescence microscope (Carl Zeiss Jena GmbH, Germany), utilizing a 514 nm laser line for excitation and a 505–550 nm band pass for detection.

### Co-Immuno Precipitation

βKNL2 was fused to a MYC tag and other binding partners to HA tag. Leaves from 2-3-week-old *N. benthamiana* plants were co-infiltrated with *Agrobacterium* carrying the interaction partner plasmid on their abaxial side, following the BIFC assay infiltration protocol. After 48 hours, 2-4 grams of infiltrated leaves were harvested, and total protein extraction was performed using Phosphate buffer (50mM NaH2PO4, 1mM β-Mercaptoethanol, 100mM NaCl, 0.5% Nonidet-P40, pH 7-7.5). Co-immunoprecipitation (Co-IP) was conducted using the Anti-MYC tag magnetic agarose trap kit from Proteintech, following the manufacturer’s protocol (https://www.ptglab.com/products/Myc-Trap-Magnetic-Agarose-ytma.htm).

The leaf material was ground using a mortar and pestle in liquid nitrogen, then extracted in Phosphate buffer. After a 30-minute incubation on ice, centrifugation at 4°C and 13000 rpm for 10 min, the supernatant was diluted at a ratio of 1:3 (grams:mL) and incubated with 50 μL MYC-Trap agarose. Agarose beads were incubated for 2 hours at room temperature or overnight at 4°C, and protein was eluted by boiling at 95°C for 5 mins with 60 μL 2× PLB loading buffer. A 10 μL aliquot of each sample was electrophoretically separated and transferred to a membrane, which was then probed with anti-HA antibodies (Anti HA tag McAb, Proteintech). An extract of not infiltrated *N. benthamiana* leaves served as a negative control and detected using a LICOR Odyssey scanner.

### Microscopy Analysis of Fluorescent Signals

Confocal fluorescence imaging was performed on a Zeiss LSM780 confocal laser scanning microscope (Carl Zeiss Jena GmbH, Germany). EYFP was visualized using a 488 nm laser line for excitation in combination with a 490–550 nm band pass for detection. Images were analyzed with the Zeiss LSM software ZEN Black edition. Fluorescence loss in photobleaching (FLIP) and fluorescence recovery after photobleaching (FRAP) were recorded using a 20x water objective (N.A. 0.8) with zoom set at 4.0 and image size 512 x 80 pixels. Recordings were made with a pixel dwell time of 1 µs and pinhole setting equaling 2 airy units. For bleaching experiments the laser intensity was set at 100%.

FRAP experiments started with three to five pre-scans after which an area of interest was bleached with the number of iterations adjusted to ensure 50-70% reduction in original fluorescence. Recovery was followed until an area of stability was reached. For FLIP experiments a small area of interest was bleached repeatedly using 1 or 5 iterations until stability was reached.

Stable transformants of *Arabidopsis* expressing fusion constructs were germinated on agar medium and roots of 7-10 days old seedlings were analysed for EYFP expression using a 40x - water objective (NA 1.2).

### AlphaFold2 protein predictions and molecular dynamics analysis

We used a local installation of AlphaFold2 in all structural predictions of the current work. We predicted the structures of βKNL2, αKNL2 and CENP-A/CENH3 and their combinations in different complexes. A complete list of the complexes we tried is in Supplementary Figure S2 with their respective confidence and sequence coverage by the multiple sequence alignment.

For the model of βKNL2 and αKNL2 bound to the nucleosome, we first predicted the centromeric histone octamer with CENP-A/CENH3, H2AW6, H2BHTB9, H4 using AlphaFold2. Then we superposed this structure with the recently resolved *Arabidopsis* euchromatic nucleosome (PDB code 7ux9 (Lee, et al. 2023)), and we found that AlphaFold2 predicts a nearly identical structure, with RMSD of 1.2 Å for the 391 aligned residues (Supplementary Figure S3). We then tried to predict the histone octamer in complex with βKNL2 and αKNL2. Some previously predicted disordered regions of the histones and of αKNL2 were neglected in the prediction of the complex to decrease computational costs (Supplementary Figure S4). We superposed this prediction with the chicken CENP-A nucleosome in complex with KNL2, recently resolved by cryo-EM (pdb code 7y71) (Jiang, et al. 2022) . AlphaFold2 correctly placed the CENPC-k motif of αKNL2 in a similar position of the experimentally resolved CENPC-like motif of chicken KNL2 (Supplementary Figure S3). βKNL2 and αKNL2 were consistently predicted to be bound by their SANTA domains, but interacting with the octameric histones in a place that would collide with the DNA and not interacting with CENP-A/CENH3. So, we sought to build a model of the nucleosome with βKNL2, αKNL2 and the CENPC-k motif of αKNL2 combining different models predicted by AlphaFold2 that presented consistent characteristics and relate well with experimental observations.

We created the proposed model in 5 steps.

**1.** We started with a centromeric nucleosome made of the predicted octameric histones (CENP-A/CENH3, H2AW6, H2BHTB9 and H4) with the DNA from the superposed *Arabidopsis* euchromatic nucleosome. We also kept the CENPC-k motif of αKNL2 in the same position predicted to bind to the histones core.
**2.** Next, we superposed the heterotrimer model of βKNL2, αKNL2 and CENP-A/CENH3 (Supplementary Figure S2K) with the CENP-A/CENH3 from the octameric nucleosomes opposed to the CENPC-k motif of αKNL2.
**3.** We superposed the heterodimer of βKNL2 and αKNL2 with the βKNL2 of the heterotrimer and rotated it by 25° around the second α-helix of CENP-A/CENH3, next to Loop1. This way we avoided the clashes of βKNL2 and αKNL2 with the histones and kept what we believe is an important interaction of βKNL2 with CENP-A/CENH3 close to Loop1.
**4.** We included the extended α-helix of motif-III (residues 213-281), that appeared in the predicted models of different dimer and trimers, by superposing residues 208-218 of βKNL2, which folded into an α-helix in both models (the βKNL2 and αKNL2 heterodimer and the homotetramer).
**5.** We performed a molecular dynamics of 150 ns in equilibrium, starting from the thus assembled chimeric model, to remove small clashes and check the stability of the proposed model. The model presented is the last conformation of the molecular dynamics.

The molecular dynamics was performed with GROMACS software (Abraham, et al. 2015). We used CHARMM36 all-atom force field (Huang, et al. 2017) with an updated port for GROMACS (http://mackerell.umaryland.edu/charmm_ff.shtml#gromacs). We used periodic boundary conditions and explicit solvation with SPC water model and neutralizing salt ions to fill a cubic box circumscribing the model with 10 Å from its edge. We ran minimization, 100 ps pressure (NVT) and 100 ps volume (NPT) equilibrations prior to the 100 ns dynamics, all steps with Particle Mesh Ewald long-range electrostatics, velocity rescale thermostat, isotropic Parrinello-Rahman pressure coupling (for NPT and dynamics) and Verlet cutoff-scheme for neighbor searching with steps of 2 fs.

We analyzed the models by its secondary structure with DSSP algorithm (Kabsch and Sander 1983) and by its interchain interactions as the distance of a query residue to the closest residue of another chain, measured by their alpha carbons and with a cutoff of 20 Å. Pictures of the structures were made with PyMOL (The PyMOL Molecular Graphics System, Version 3.0 Schrödinger, Inc.). Charge distribution surfaces were calculated with APBS Electrostatics method inside PyMOL and PDB2PQR for calculation of Poisson-Boltzmann electrostatics (Dolinsky, et al. 2004).

## Results

### SANTA and C-terminus are required for efficient centromeric targeting of βKNL2

In plants, contrasting to other organisms, two KNL2 protein variants (α and βKNL2) containing conserved SANTA domain have been identified (Figure 1A) (Zuo, et al. 2022). Though the SANTA-containing N-terminus of αKNL2 is not required for centromeric targeting (Lermontova, et al. 2013; Sandmann, et al. 2017), the role of the SANTA domain in βKNL2 remains elusive. Online prediction tools like PredictProtein, LambdaPP, DNABIND predicted that the SANTA and C-terminus of βKNL2 are likely involved in both protein-protein (PPI) and protein-nucleic acid interactions (Supplementary Figure S5).

Thus, to determine the role of SANTA and other regions of βKNL2 in centromere targeting, three different truncation variants were generated: i) fragment lacking the N-terminal part (βKNL2ΔN), ii) fragment lacking the SANTA (βKNL2ΔSANTA) and iii) fragment lacking the C-terminal part (βKNL2ΔC) (Figure 1B). All fragments were fused with EYFP and transiently expressed in *N. benthamiana* leaves. βKNL2ΔN and βKNL2ΔSANTA showed localization patterns similar to full-length βKNL2, while βKNL2ΔC localized to the nucleoplasm and cytoplasm (Supplementary Figure S6).

We generated stable *Arabidopsis* lines expressing each construct and analyzed the localization of the corresponding fusion proteins in T_2_ seedling root tips. βKNL2ΔN, fused with EYFP, exhibited centromeric localization similar to that of full-length βKNL2 (Figure 1C). However, βKNL2ΔSANTA transformants showed less efficient centromeric localization, while βKNL2ΔC transformants had mis-localization to the cytoplasm and nucleoplasm similar to what was observed in transient expression (Figure 1C, Supplementary Figure S6). This confirmed that efficient centromeric localization of βKNL2 requires both the SANTA domain and C-terminus.

### Motif-III in the C-terminus of βKNL2 is required for the nuclear localization of βKNL2

Previously, we identified four βKNL2 specific conserved motifs (Zuo, et al. 2022). Extended sequence analysis to approximately 100 KNL2 variants without CENPC-k motif from monocots and dicots (Supplemental File.1) confirmed that the conserved motifs are specific to βKNL2. Each motif spanning approximately 20 AA (Figure 2A, Supplemental File.2). The first motif (residues 24-44) is located at the N-terminus, upstream of SANTA, while the other three - motif-I (residues 164-185), motif-II (residues 186-205), and motif-III (residues 229-249)—are positioned in the C-terminus, directly downstream of the SANTA (Figure 2A).

**Figure. 2.**
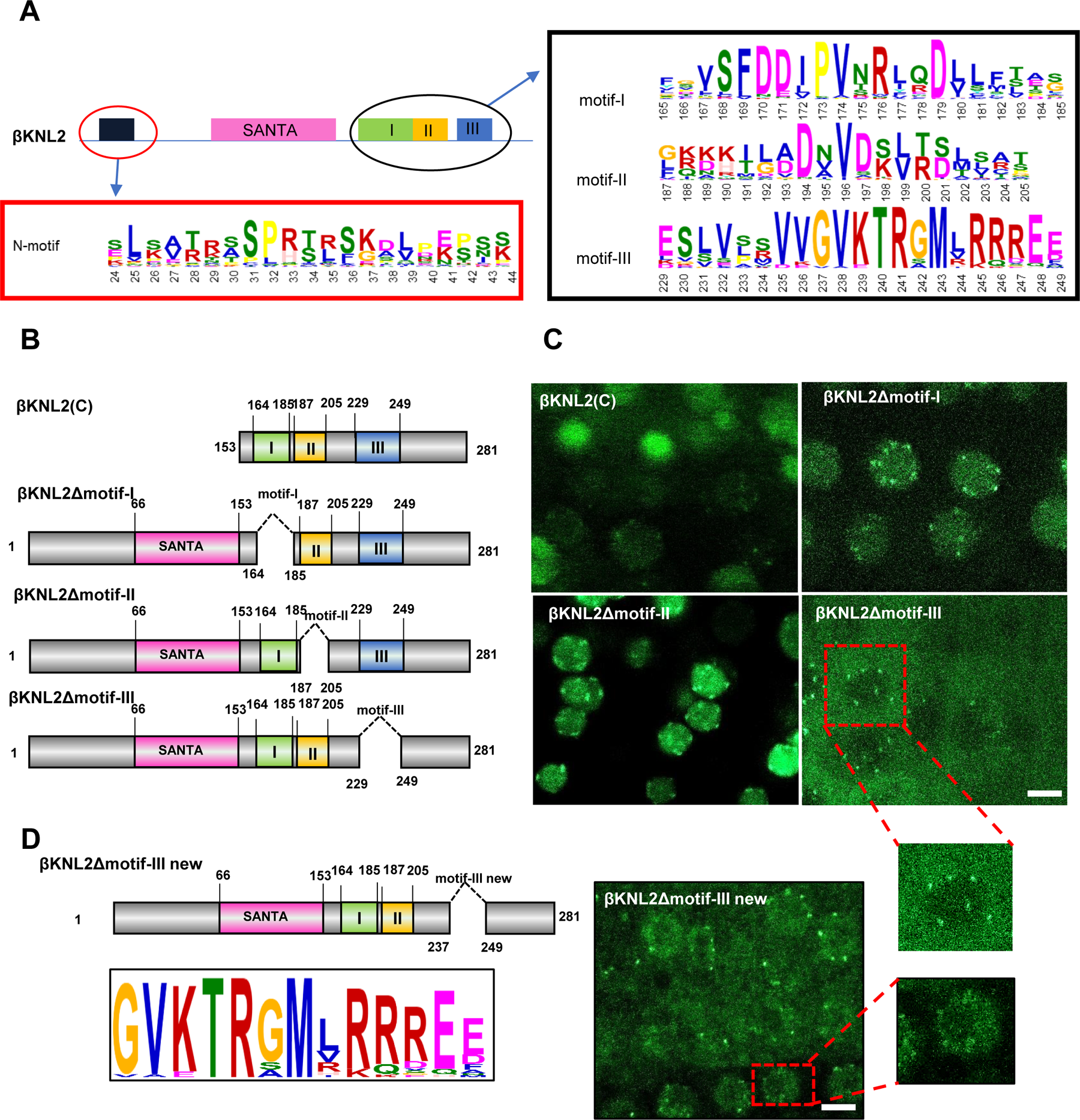
βKNL2 nuclear localization is regulated by conserved C-terminal motif-III. (A) Schematic representation of the βKNL2 protein structure. Newly identified conserved motifs at the N- and C-termini of βKNL2 are encircled and accompanied by WebLogo alignments showing conserved AA across dicots (B) Truncated constructs of βKNL2 to elucidate the functional role of the entire C-terminus and its individual motifs. The βKNL2(C) variant retains only the C-terminal region, while the βKNL2Δmotif-I, βKNL2Δmotif-II and βKNL2Δmotif-III variants omit one of the three corresponding motifs. (C) Localization patterns of EYFP-Tagged βKNL2 truncated protein variants (referred in 2B) in *Arabidopsis* root tip nuclei. βKNL2(C) showed reduced centromeric localization. βKNL2Δmotif-I and βKNL2Δmotif-II fragments displayed normal nuclear and centromeric signals, whereas βKNL2Δmotif-III exhibited mislocalization to the cytoplasm, indicating the critical role of motif-III in nuclear targeting. (D) βKNL2Δmotif-III new, truncated variant excluding highly conserved AA of motif-III, but including the predicted SUMOylation site accompanied by WebLogo alignment. In contrast to the βKNL2Δmotif-III fragment, this variant showed partially recovered nuclear localization (right panel). Scale bar 5µm

To explore whether the C-terminus alone is capable to localize to the centromere, we generated a truncated construct containing solely the C-terminal part (βKNL2(C)) fused to EYFP (Figure 2B). This variant showed a low centromere targeting efficiency in root tip nuclei of stably transformed *Arabidopsis* (Figure 2C, Supplementary Figure S7). This confirms our assumption that efficient centromeric targeting of βKNL2 requires the C-terminal part and the SANTA.

Furthermore, the influence of C-terminal motifs on βKNL2 localization was tested by generating constructs with individual deletions of each motif, which were then fused to EYFP (Figure 2B). These constructs were then employed for transient expression in *N. benthamiana* and for stable expression in *A. thaliana*. The T_2_ lines of *A. thaliana* expressing βKNL2Δmotif-I and βKNL2Δmotif-II constructs showed fluorescence at centromeres and in nucleoplasm. In contrast, βKNL2Δmotif-III expressing lines exhibited mis-localization to the cytoplasm (Figure 2C, Supplementary Figure S7). Similar results were obtained when all constructs were transiently expressed in *N. benthamiana* (Supplementary Figure S7).

BLASTP analysis of “motif-III” identified numerous matches across eukaryotes and prokaryotes, particularly with transcription factors and nucleic acid binding proteins. Further analysis of motif-III with ELM and GPS-biocuckoo revealed a potential SUMOylation site spanning residues 221-231 which overlaps with motif-III (residues 229 to 231) (Supplementary Figure S8, Supplemental File.3). We designed a new truncated construct (βKNL2Δmotif-III new) that includes the predicted SUMOylation region but excludes the conserved region of motif-III (Figure 2D). The lines expressing βKNL2Δmotif-III new showed partial recovery of nucleoplasm localization compared to βKNL2Δmotif-III (Figure 2D). These observations suggest that motif-III could be SUMOylated, which might be crucial for proper localization of βKNL2. However, the remaining conserved part of the motif-III is also required for efficient targeting of βKNL2 to centromeres.

### βKNL2 binds to the centromeric DNA *in vitro*

Different inner kinetochore proteins associate with the centromeric DNA (Cheeseman 2014; Yamagishi, et al. 2014; Musacchio and Desai 2017; Jiang, et al. 2022) . For example, αKNL2 associates with DNA in a sequence-independent manner *in vitro* and preferentially with *pAL1 in vivo* (Sandmann, et al. 2017; Yalagapati, et al. 2024). Different predictions show that βKNL2 has ability to interact with nucleic acids (Supplementary Figure S5) at the regions shown in Figure 3A. To validate this, we performed Electrophoretic Mobility Shift Assay (EMSA) using IRD700 labelled *pAL1* DNA and *in-vitro* expressed FLAG-βKNL2. IRD700-*pAL1* amplified from *Arabidopsis* genomic-DNA, so it contained monomeric and multimeric repeats of *pAL1* (Supplementary Figure S9). FLAG-αKNL2(C) served as a positive control and expression of both proteins was verified by Western blot (Supplementary Figure S9). We observed upward shift in the lane of IRD700-*pAL1* when incubated with FLAG-βKNL2, similar to FLAG-αKNL2(C) (Figure 3B). However, when unlabeled *pAL1* used as a competitor along with labelled probes, a reduced shift was observed, indicating successful competition.

**Figure. 3.**
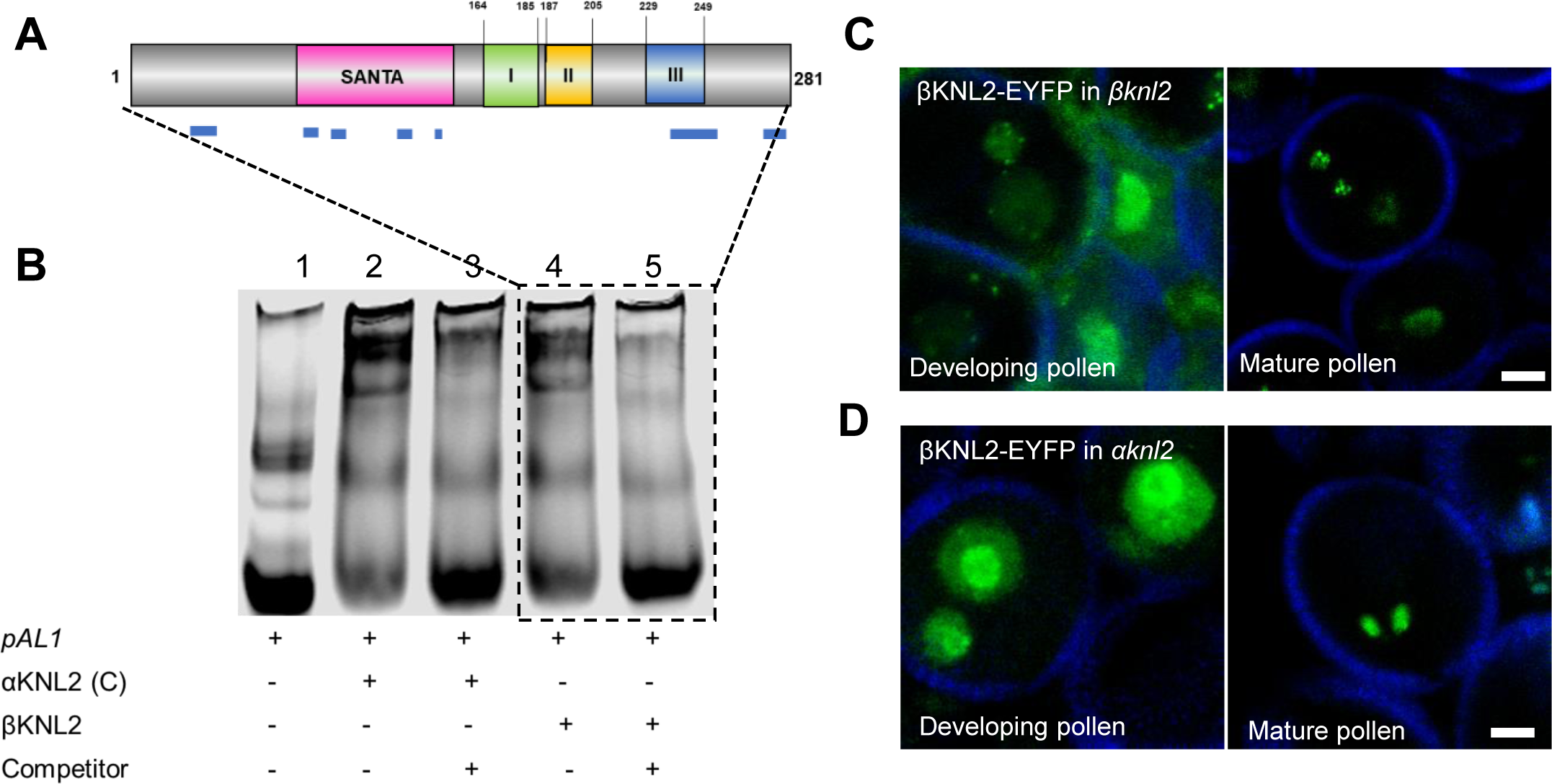
βKNL2 binds to centromeric DNA and its centromeric recruitment depends on αKNL2 in pollen nuclei. (A) Schematic representation of the βKNL2 protein with predicted DNA-binding regions marked by blue bars below. (B) EMSA on native gel showing FLAG-βKNL2 binding to centromeric *pAL1* DNA. IRD700-labeled *pAL1* monomer and multimeric repeats probes were amplified from *Arabidopsis* genomic DNA (lane 1). IRD700-*pAL1* probes shifted upwards indicating binding by in vitro expressed FLAG-αKNL2(C) as a positive control (lane 2) and FLAG-βKNL2 (lane 4). Competition with an excess of unlabeled *pAL1* DNA reduced the shifts confirming specific binding of FLAG-αKNL2 (lane 3) and FLAG-βKNL2 (lane 5) to *pAL1* DNA. (C-D) Disturbed centromeric localization of βKNL2-EYFP in *αknl2* compared to *βknl2* mutant pollen nuclei, indicating centromeric recruitment of βKNL2 is dependent on αKNL2. (C) In *βknl2* mutant transformants, βKNL2-EYFP localizes to centromeres and nucleoplasm in the nuclei of developing and mature pollen. (D) In *αknl2* mutant transformants, βKNL2-EYFP fluorescence is localized in the nucleoplasm and nucleoli. Pollen wall autofluorescence in blue. Scale bar: 5µm

As mentioned earlier, BLAST analysis of motif-III resulted in several hits with various nucleic-acid binding proteins. To verify if motif-III is capable to bind to the DNA, we synthesized a peptide containing the motif-III (V**K/E**TRGMLRRREE**Y/G**EASIG**K/E**R). Since this peptide was designed to perform EMSA experiments and to generate antibodies against different Brassica species, it contains varied residues than motif-III. Increased concentrations of this peptide (10-30µg) in the EMSA experiment with *pAL1* revealed an increased shift of *pAL1* (Supplementary Figure S10). With this, we conclude that βKNL2 can bind to centromeric DNA possibly via extended motif-III.

### βKNL2 depends on CENPC-like motif containing αKNL2 for centromeric localization

Previous studies have shown that the M18BP1/KNL2 requires the CENPC-like motif for centromeric localization (Hori, et al. 2017; Sandmann, et al. 2017; Smith and Maddox 2017; Jiang, et al. 2022). Interestingly, βKNL2 exhibits centromeric localization even without this motif. In our previous study, we hypothesized that βKNL2 might rely on CENPC-like motif containing proteins (αKNL2 or CENP-C) for its centromere targeting (Zuo, et al. 2022).

To test this hypothesis, we introduced βKNL2::βKNL2-EYFP construct into *αknl2* and *βknl2* mutants. We expected reduced centromeric localization of βKNL2-EYFP in *αknl2* mutants, reflecting βKNL2 dependency on αKNL2. We assessed βKNL2-EYFP in floral tissues of both mutants, given that both KNL2 genes have high expression in meristematic cells.

We initially focused on pollen (Fig 3 C-D, Supplementary Figure S11). In *βknl2* mutants, βKNL2-EYFP was present in the centromeres and nucleoplasm of both generative and vegetative nuclei in bi-nucleate pollen and to the sperm nuclei in tri-nucleate pollen (Figure 3C). Interestingly, vegetative nuclei of mature pollen, showed exclusive nucleoplasm localization of the fluorescence. In contrast, *αknl2* mutant transformants exhibit fluorescence in both the nucleoplasm and nucleolus, but lack distinct centromeric signals in the bi-nucleate and tri-nucleate pollen nuclei (Figure 3D, Supplementary Figure S11).

Next, we examined developing seeds with embryos, and both *αknl2* and *βknl2* transformants showed similar centromeric localization patterns in embryo (Supplementary Figure S11). Notably, other tissues in the developing seeds displayed disrupted centromeric localization similar to that observed in pollen nuclei. Thus, we conclude that centromeric localization of βKNL2 depends on αKNL2 in a tissue-specific manner.

### βKNL2 form homodimer and heterodimer with αKNL2

Across various organisms, kinetochore components tend to form homodimers and oligomers (Stellfox, et al. 2016; Subramanian, et al. 2016; Pan, et al. 2017). The Mis18 complex is formed by a M18BP1/KNL2 homodimer and a hetero-oligomer of Mis18α&β. Notably, plants possess two KNL2s (α&βKNL2) and lack the Mis18α&β. Thus, we addressed the question of whether βKNL2 can dimerize similar to Mis18 components. Using AlphaFold2, we predicted βKNL2 monomer, dimer and oligomers with αKNL2 in various combinations (Supplementary Figure S2). Monomer features structured regions in SANTA (with β-roll linked to small α-helices and loops) and C-terminus (three α-helical motifs), while the rest appears disordered (Figure 4A). The proline-rich N-terminus likely disrupts helix and sheet formations, aligning with its intrinsic disorder. Exposed positively charged and hydrophobic surfaces on βKNL2 suggest roles in DNA binding and PPI (Supplementary Figure S12).

**Figure 4.**
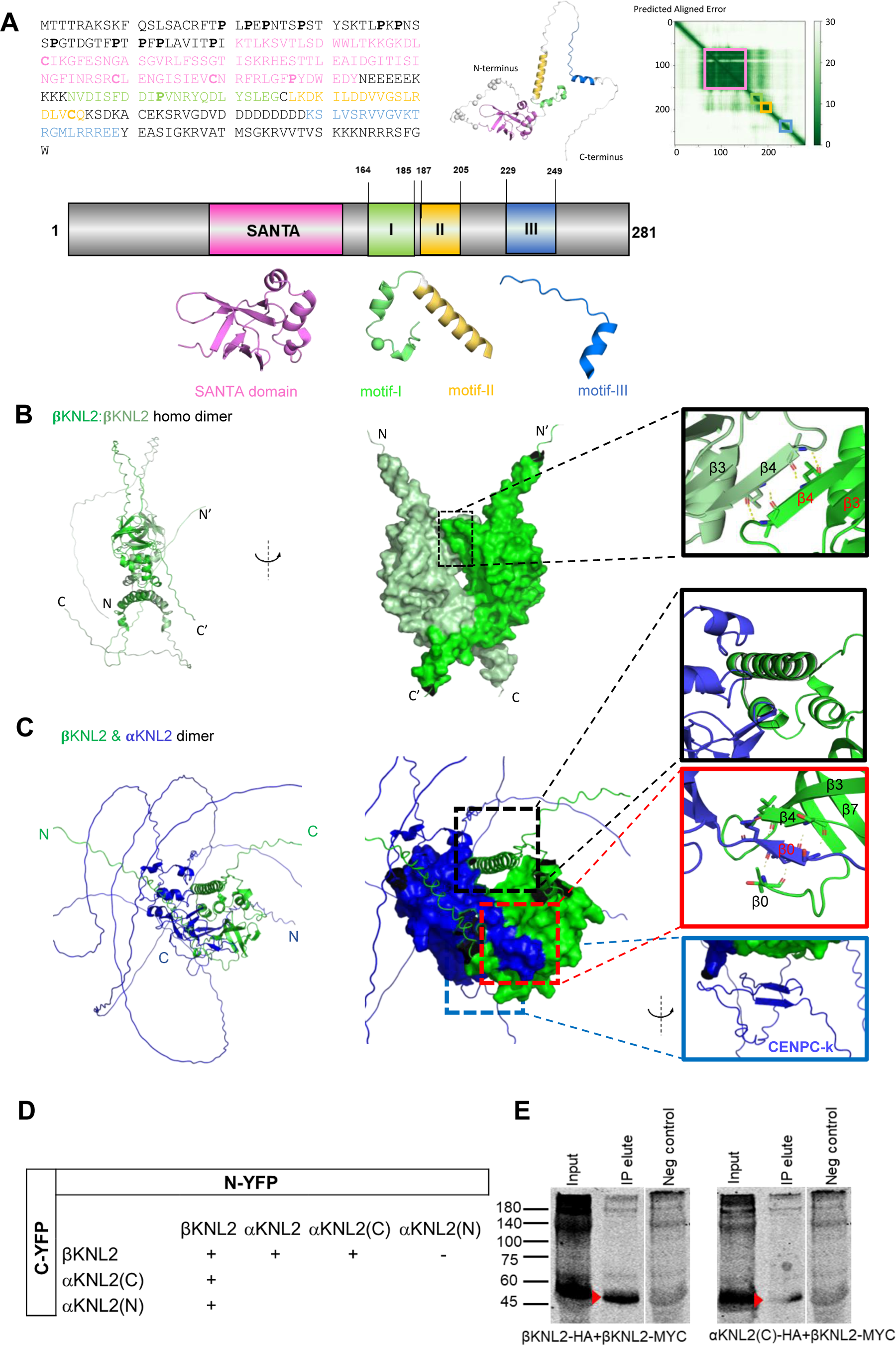
βKNL2 form homo- and heterodimers. (A) Structural prediction of βKNL2 by AlphaFold2. The SANTA domain (pink), motifs I (green), II (yellow), and III (blue) of βKNL2 are highlighted. Annotations detail these elements alongside a Predicted Aligned Error (PAE) plot, indicating a confident structure for the SANTA domain and the relative mobility of the three motifs. (B) AlphaFold2-predicted βKNL2 homodimer structure. The left view depicts secondary structures and the right view presents a surface representation, each monomer distinctly colored. A zoomed view illustrates the interaction site predicted by AlphaFold2 between the β4 strands of the SANTA domain, with additional predicted contacts in the C-terminal motifs. (C) AlphaFold2-predicted heterodimer of βKNL2 (green) and αKNL2 (blue) shows the interaction mediated through the SANTA and C-terminal conserved motifs. Zoomed insets top highlights motifs I and II interaction with αKNL2; middle: an intermonomeric β-sheet formation between βKNL2 (β0 and β4 strands) and αKNL2 (β0 strands); and bottom: CENPC-k motif of αKNL2 being close to the SANTA domains. (D) BIFC assays for confirmation of βKNL2 homo and hetero dimer formations: Positive interactions were observed between βKNL2:βKNL2, βKNL2:αKNL2 and βKNL2:αKNL2(C). (E) Western blots with anti-HA tag antibodies display monomeric size bands (red arrow heads) from co-immunoprecipitation (co-IP) performed with MYC agarose trap. Left shows co-IP experiment of βKNL2-MYC and βKNL2-HA and the right shows co-IP experiment of βKNL2-MYC and αKNL2(C)-HA.

AlphaFold2 predicts that βKNL2 can form homodimers and heterodimers with αKNL2 through the SANTA and C-terminal motifs (Figure 4B-C). Notably, the N-terminal tails (pre-SANTA) of both αKNL2 and βKNL2 fold into β-strands, complementing each other to form an intermonomeric β-sheet (Figure 4C). We observed structural changes in the C-terminal motifs of βKNL2 in dimeric or trimeric forms (Supplementary Figure S13).

To validate homo- and heterodimer formation, Bimolecular Fluorescence Complementation (BiFC) assays were performed. βKNL2 and αKNL2 were fused to the N (VN)- and the C (VC)-terminus of Venus in both directions. To surpass overexpression difficulty of αKNL2 due to its proteasomal regulation (Lermontova, et al. 2013), we employed MG115 proteasomal inhibitor. We observed fluorescence when βKNL2-VC is co-infiltrated with βKNL2-VN and αKNL2-VN, thus validating βKNL2 homo and heterodimer formations (Figure 4D). Furthermore, we performed BIFC assay using truncated αKNL2 constructs, αKNL2(N) (residues 1-362, containing the SANTA) and αKNL2(C) (residues 363-598, with CENPC-k motif). αKNL2(C) showed interaction with βKNL2, whereas αKNL2(N) failed to show interaction in both directions (Figure 4D).

We performed co-immunoprecipitation (co-IP) with anti-MYC trap by transiently expressing βKNL2-MYC-VN, βKNL2-HA-VC and αKNL2(C)-HA-VC in combinations. Monomer-sized bands were detected (between 45kDa-60kDa) higher than the predicted 43.42kDa for βKNL2-HA-VC (Figure 4E, left) and 38.21 kDa for αKNL2(C)-HA-VC (Figure 4E, right) from precipitated elutes. This could be due to post-translational modifications. A similar migration pattern was observed for βKNL2 and αKNL2(C) when expressed *in vitro* using a wheat germ system (Supplementary Figure S9A). These results substantiate the structural predictions and support the formation of βKNL2 homo and heterodimers.

### βKNL2 is highly dynamic and loosely bound to centromeric nucleosome through its SANTA and C-terminal conserved motifs

We performed FLIP (Fluorescence Loss in Photobleaching) and FRAP (Fluorescence Recovery After Photobleaching) experiments to track the dynamics of βKNL2-EYFP within *Arabidopsis* elongated nuclei (Supplementary Figure S14). These nuclei are structurally stable and less active and they do not move out of focus. FLIP showed that βKNL2-EYFP has high mobility, as indicated by rapid fluorescence reduction followed by swift movement. This observation was supported by FRAP, which revealed that rapid recovery of fluorescence with a short half-time (t1/2) of approximately 0.7 seconds. Similar turnover rates and mobility were also observed for EYFP-αKNL2 as well (Lermontova, et al. 2013), indicating that both KNL2 proteins bind flexibly to nucleosomes.

We used AlphaFold2 to model how KNL2 proteins bind to centromeric nucleosomes, but these models couldn’t accurately position βKNL2 and αKNL2 (Supplementary Figure S15). Thus, we created a chimeric model combining AlphaFold2 predictions, which was independently validated twice with molecular dynamics for the stability (methods). This model aligns with the experimentally resolved structures, like the αKNL2 CENPC-k motif binds to histones similar to ggKNL2 (Figure 5A) (Jiang, et al. 2022). Both SANTA domains interact with each other and contact all histones and DNA. Residues from motif-III (230 to 237) were positioned to be exposed and accessible for SUMOylation (Figure 5B). Furthermore, starting from motif-III, a long stretch of positively charged interface forms extended α-helix, which embraces the DNA (Figure 5C). In total this placement KNL2s complement the charge distribution to the other regions of the nucleosome. The results including the dynamics of βKNL2-EYFP and other findings from our study support our proposed model.

**Figure 5.**
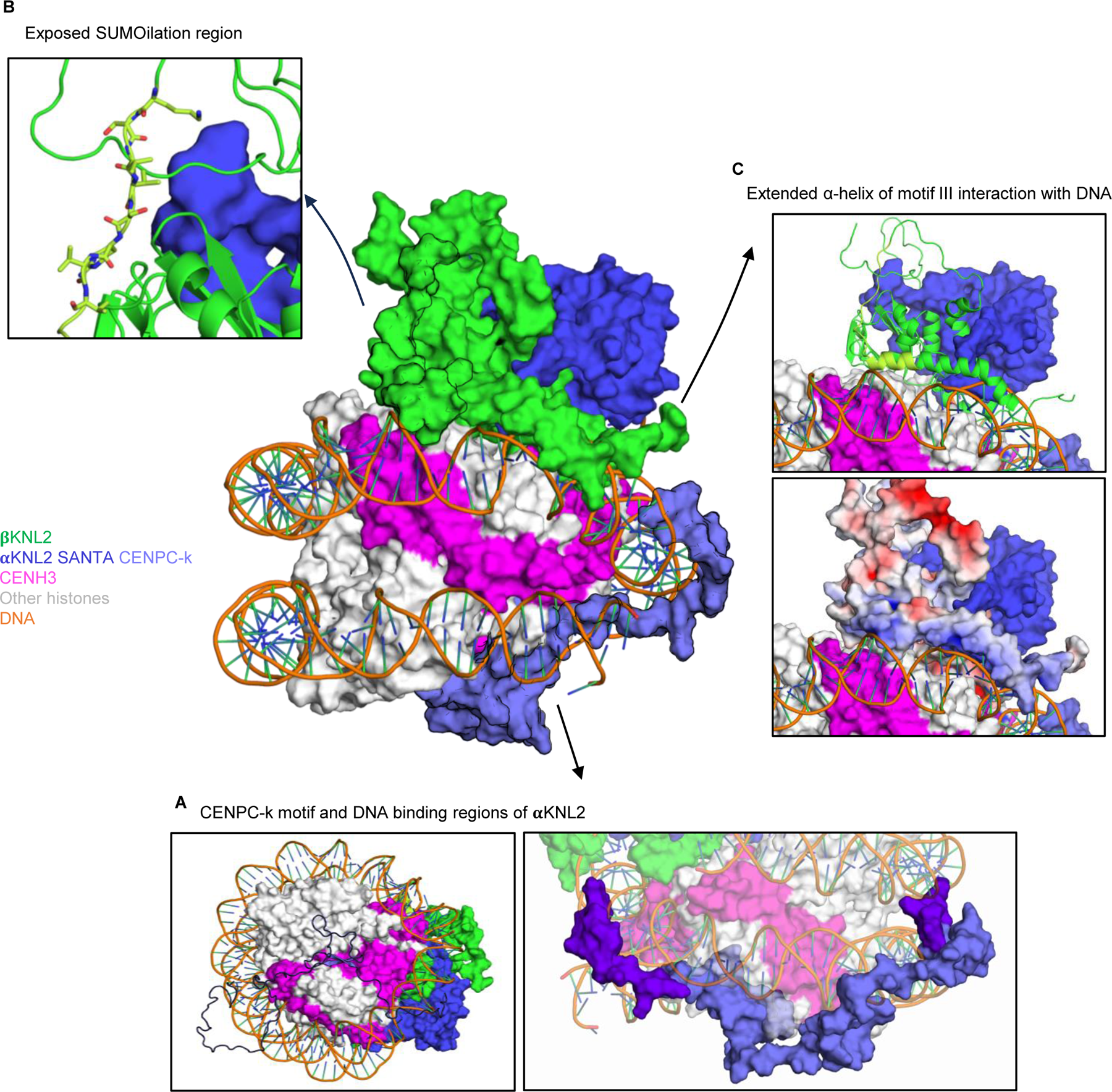
Proposed model of plant-specific inner kinetochore assembly involving βKNL2, αKNL2, and the centromeric nucleosome. The primary visualization highlights the protein surfaces of βKNL2 (green), αKNL2 (blue), and CENP-A/CENH3 (pink) as they interact with the double helix of DNA. These color codes are consistent throughout all detailed views. (A) Left: The CENPC-k motif (blue) is positioned on the underside of the nucleosome, close to the lower monomer of CENP-A/CENH3. This was similar to the ggKNL2-nucleosome interaction detailed by Jiang et al., 2022. Right: A purple overlay marks DNA-binding regions on αKNL2, as identified by Yalagapati et al., 2024. (B) The surface exposure of conserved residues at a potential SUMOylation site, as described in the results of this study and illustrated in Figure 2D, highlights their accessibility to solvent. (C) motif-III of βKNL2, with its positively charged surface, wraps around the DNA. Emphasizing its interaction with phosphate groups, which facilitates secure attachment to the nucleosome. These interactions were verified in this study and are presented in Figure 3B.

## Discussion

In metazoans, a single M18BP1/KNL2 variant is maintained, while plants have evolved different versions: α-and βKNL2 in dicots, and γ-and δKNL2 in monocots. Notably, the centromeric localization of αKNL2 in dicots depends on the presence of the CENPC-k motif (Sandmann, et al. 2017). In contrast, βKNL2 localizes to centromeres even without this motif Zuo, et al. (2022).

Analysis of the βKNL2 domain structure revealed four specific conserved motifs: one in the N-terminus and three in the C-terminus relative to the SANTA domain.

Our experimental data show that while deletion of the N-terminus of βKNL2 does not affect its centromeric targeting, removal of the SANTA domain or the C-terminal region significantly impairs this localization (Figure 2, Supplementary Figure S6). This indicates that centromeric targeting of βKNL2 primarily relies on the SANTA domain and C-terminal motifs, independent of the N-terminal region. Further specificity was explored using constructs with separate deletions of motifs I, II, and III. Results showed that motif-III (229-249 aa) in the C-terminus is essential for correct localization of βKNL2, as its deletion causes the protein to mislocalize to the cytoplasm, similar to the complete deletion of the C-terminus. In contrast, deletions of motif-I and motif-II do not affect centromeric localization(Figure 3, Supplementary Figure S7). . Alphafold2 predicted structural changes in these motifs when βKNL2 in dimer or oligomer forms () suggesting these motifs may have functional role in protein structure

Further investigation revealed that motif-III overlaps with a SUMO site (221-231 aa) and can undergo SUMOylation. However, deleting the highly conserved region of motif-III (237-249 aa) while retaining the predicted SUMOylation region led to partial restoration of nucleoplasm localization. This demonstrates the crucial role of the entire motif-III in targeting the βKNL2 protein to the nucleus and centromeres. The role of SUMOylation in βKNL2 localization remains to be fully elucidated. However, it is known that SUMOylation plays an important role in the correct localization and function of kinetochore proteins. For instance, loss of the SUMO protease SENP6 in human cells results in the mislocalization of essential kinetochore proteins CENP-C, CENP-I (Mitra, et al. 2020), M18BP1/KNL2, and other kinetochore proteins (Liebelt, et al. 2019).

Our study identified specific regions essential for the centromeric targeting of βKNL2, prompting further investigation into the mechanisms of this targeting process and the role of βKNL2 in kinetochore assembly. French and Straight (2019) showed that human M18BP1/KNL2 localizes to centromeres during metaphase by binding to the CENPC-motif containing CENP-C protein using a conserved SANTA domain. Notably, in plants, both αKNL2 and CENP-C contain the CENPC-motif. In reproductive tissues like pollen and developing seeds, but not in embryos, centromeric localization of βKNL2 relies on αKNL2, indicating a tissue-specific recruitment mechanism (Supplementary Figure S11).

In humans, M18BP1/KNL2 forms homodimers that interact with a Mis18α&β 4:2 hexamer (Pan, et al. 2017). In plants, no Mis18α or Mis18β have been identified, but instead, two KNL2 variants are present. Thus, different compositions of KNL2 complexes can be expected in plants. Structural predictions and experimental evidence show that βKNL2 can interact with αKNL2 and form homodimers. Predicted structures indicate that dimerization of βKNL2 is mediated by the pre-SANTA and SANTA domain, similar to *Xenopus* and human KNL2 (Pan, et al. 2017; French and Straight 2019). No direct interactions were detected between βKNL2 and the SANTA-containing N-terminus of αKNL2, suggesting these interactions may require full-length αKNL2.

Since deletion of the proline-rich N-terminus did not affect the centromeric localization of βKNL2, it is conceivable that it might enhance oligomerization and complex stability. The N-terminal prolines are predicted to be phosphorylated, indicating possible regulatory control through post-translational modifications. In vertebrates, Cyclin-dependent kinase (CDK) regulates M18BP1/KNL2 centromere localization through dimerization and oligomerization (Pan, et al. 2017; French and Straight 2019). Interestingly, we identified a conserved region just before the SANTA (TPV/IK), recognized as a CDK motif in the N-terminus (Supplemental File.3). Furthermore, AlphaFold2 and molecular dynamics predict that βKNL2 and αKNL2 bind to the nucleosome. This prediction also shows that the SANTA domains and pre-SANTA conserved amino acids (S_14_F_15_Q_16_ in αKNL2 & T_58_P_59_V/I_60_K_61_ in βKNL2) form an intermonomeric β-sheet. All these findings underscore the role of β-SANTA in centromere targeting and dimerization with αKNL2.

The inner kinetochore consists of structural and DNA-binding proteins that typically bind to centromeres (Cheeseman 2014; Ramakrishnan Chandra, et al. 2024). αKNL2 can bind centromeric DNA *in vitro* and *in vivo* (Sandmann, et al. 2017) and its centromeric localization requires the presence of DNA-binding regions in addition to the CENPC-k motif (Yalagapati, et al. 2024). Here we show that βKNL2 can bind to centromeric DNA independent of αKNL2, supporting βKNL2’s involvement in inner kinetochore dynamics and chromatin interactions. In different organisms, M18BP1/KNL2 and CENP-C assume common roles in different cell cycle stages, such as CENP-A loading and kinetochore assembly. The CENP-C cupin domain, a feature absent in plants, oligomerizes for kinetochore assembly (Hara, et al. 2023). Highlighting these unique features of plants with two KNL2s and their ability to dimerize raises questions about their adaptation to a different kinetochore assembly platform. We propose that βKNL2, together with αKNL2 and/or CENP-C, facilitates or supports the kinetochore assembly platform (Figure 6). Evidence such as the consistent presence of βKNL2 at centromeres throughout the cell cycle, reduced or abolished CENP-A/CENH3 loading onto centromeres when βKNL2 expression is disrupted, and its interaction with αKNL2 and CENP-A/CENH3 all support its role in the kinetochore assembly platform (Zuo, et al. 2022).

**Figure 6.**
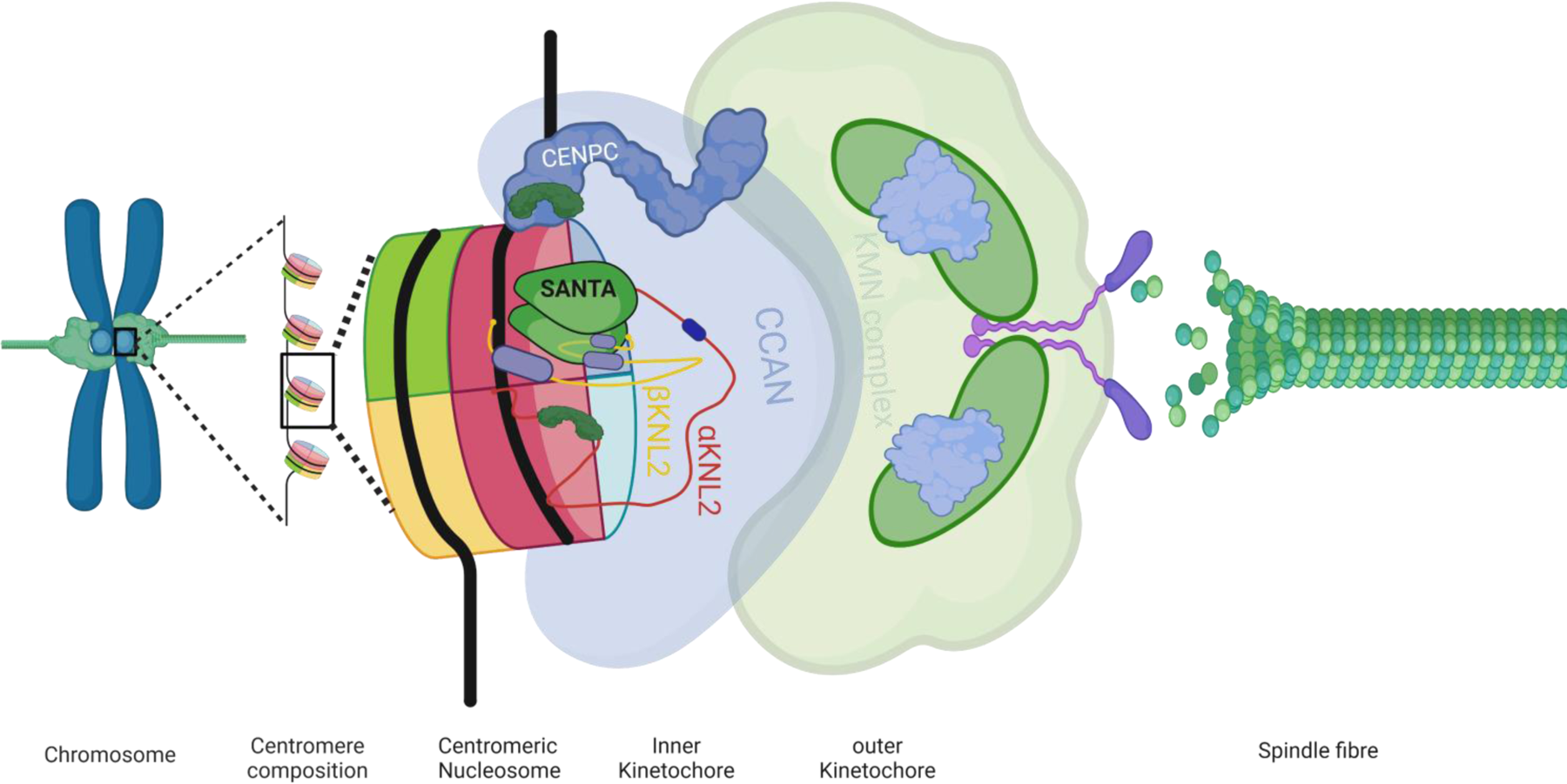
Graphical model of βKNL2 dimerization supporting centromere assembly and kinetochore platform. This graphical representation illustrates how βKNL2 dimerizes to facilitate the assembly and function of the centromere and kinetochore structures. βKNL2 forms dimers with αKNL2 and interacts with the centromeric nucleosome. These interactions are mediated through the SANTA domain and conserved motifs in the C-terminus, which are essential for stabilizing the kinetochore platform and promoting efficient centromere assembly.

## Acknowledgements

The authors thank Dr. Jos Schippers group for help with EMSA establishment. Dr. Dmitri Demidov for valuable discussions and support in establishing protein biochemistry methods in the group and Heike Kuhlmann and Annette Heber for technical assistance. R.Y. was supported by WIPANO Wissens und Technologietransfer durch Patente und Normen project grant (03THWST001) and breeding company Enza Zaden, A.S.C by Ministry of Science, Energy, Climate Protection and Environment of the State of Saxony-Anhalt. The publication of this article was funded by the German Research Foundation (DFG)— HE 9114/1-1.

## Author information

### Authors and Affiliations

Leibniz Institute of Plant Genetics and Crop Plant Research (IPK) Gatersleben, Corrensstrasse 3, D-06466 Seeland, Germany.

Ramakrishna Yadala, Amanda S Camara, Surya P Yalagapati, Pascal Jaroschinsky, Tobias Meitzel Twan Rutten, Thu-Giang T Bui and Inna Lermontova

UMR 1332 Biologie du Fruit et Pathologie, Univ. Bordeaux, INRAE, 33883 Villenave d’Ornon, France Tobias Meitzel

Graduate School of Frontier Biosciences, Osaka University, Suita, Japan Mariko Ariyoshi and Tatsuo Fukagawa

### Contributions

R.Y and I.L, conceived study, R.Y, A.S.C, T.F and I.L designed experiments. R.Y, A.S.C, M.A performed protein structure predictions. R.Y and P.J performed mutagenesis and localization studies. S.P.Y, R.Y and T. G. T.B responsible for EMSA. R.Y and T.M plasmids construction and cloning. R.Y and T.R microscopy analysis. R.Y, A.S.C and I.L wrote the manuscript. R.Y, A.S.C, M.A, T.F and I.L revised and corrected the manuscript. I.L supervised and received funding for the study.

### Correspondence

Correspondence to Ramakrishna Yadala or Inna Lermontova

---

**Supplementary Figure S1.**
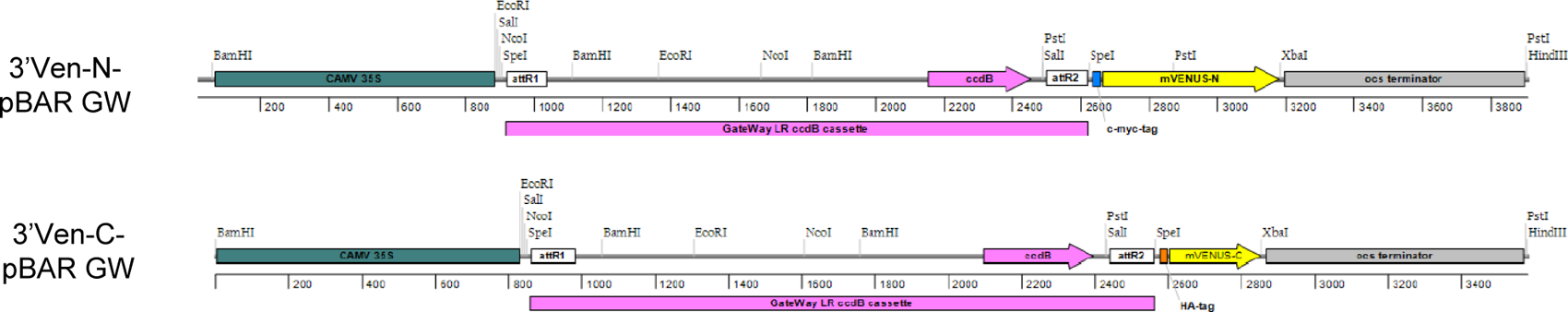
Schematic Diagram of BiFC Expression Vector Construction. The construction of BiFC expression vectors using the pPZP200BAR binary vector. The CAMV35S promoter and OCS terminator were integrated via overlap extension PCR. The GateWay LR ccdB cassette along with mVENUS-N or mVENUS-C, each tagged with cMYC or HA respectively, were amplified by PCR and subsequently cloned into the pPZP200BAR vector between the CAMV35S promoter and OCS terminator.

**Supplementary Figure S2.**
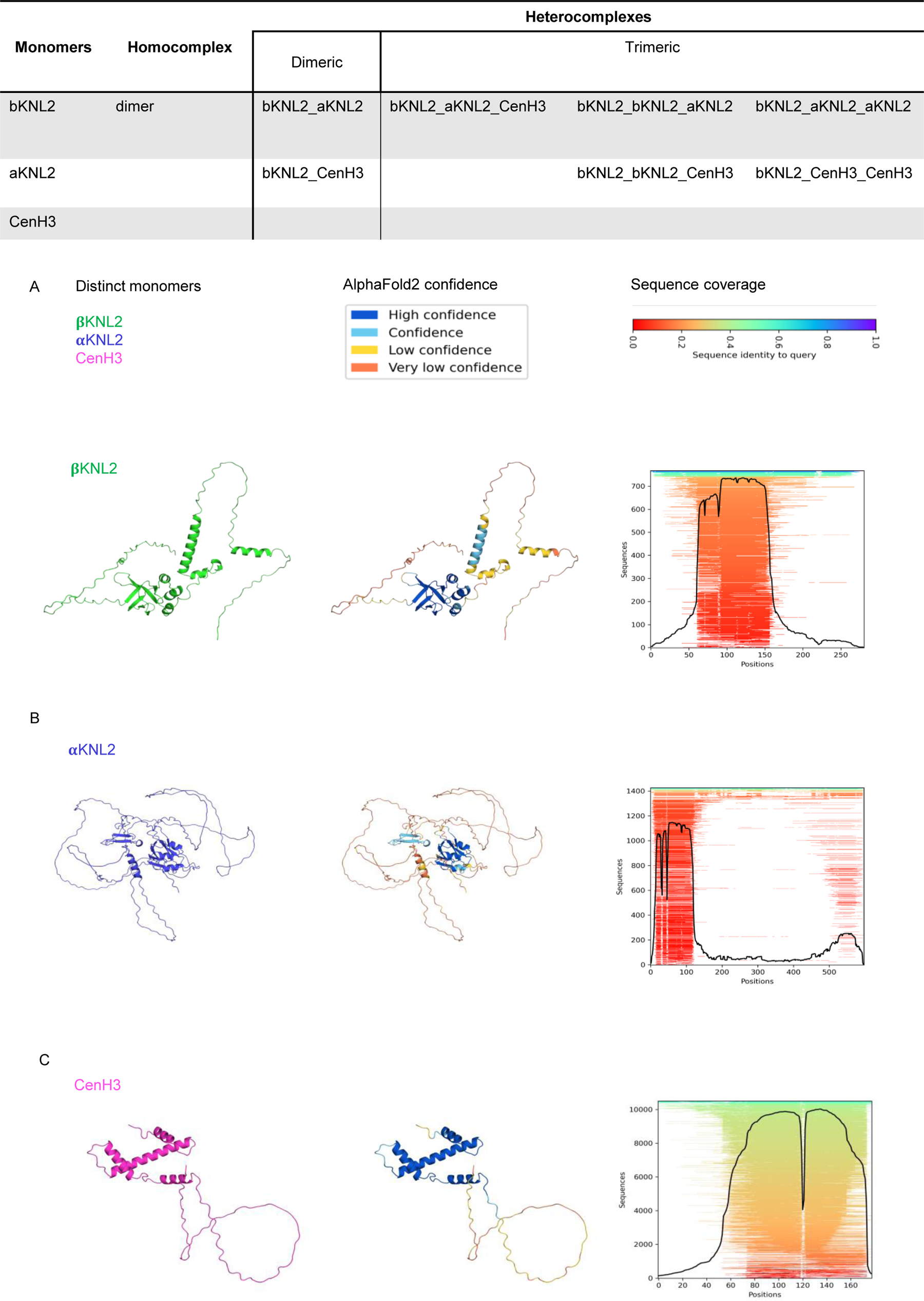
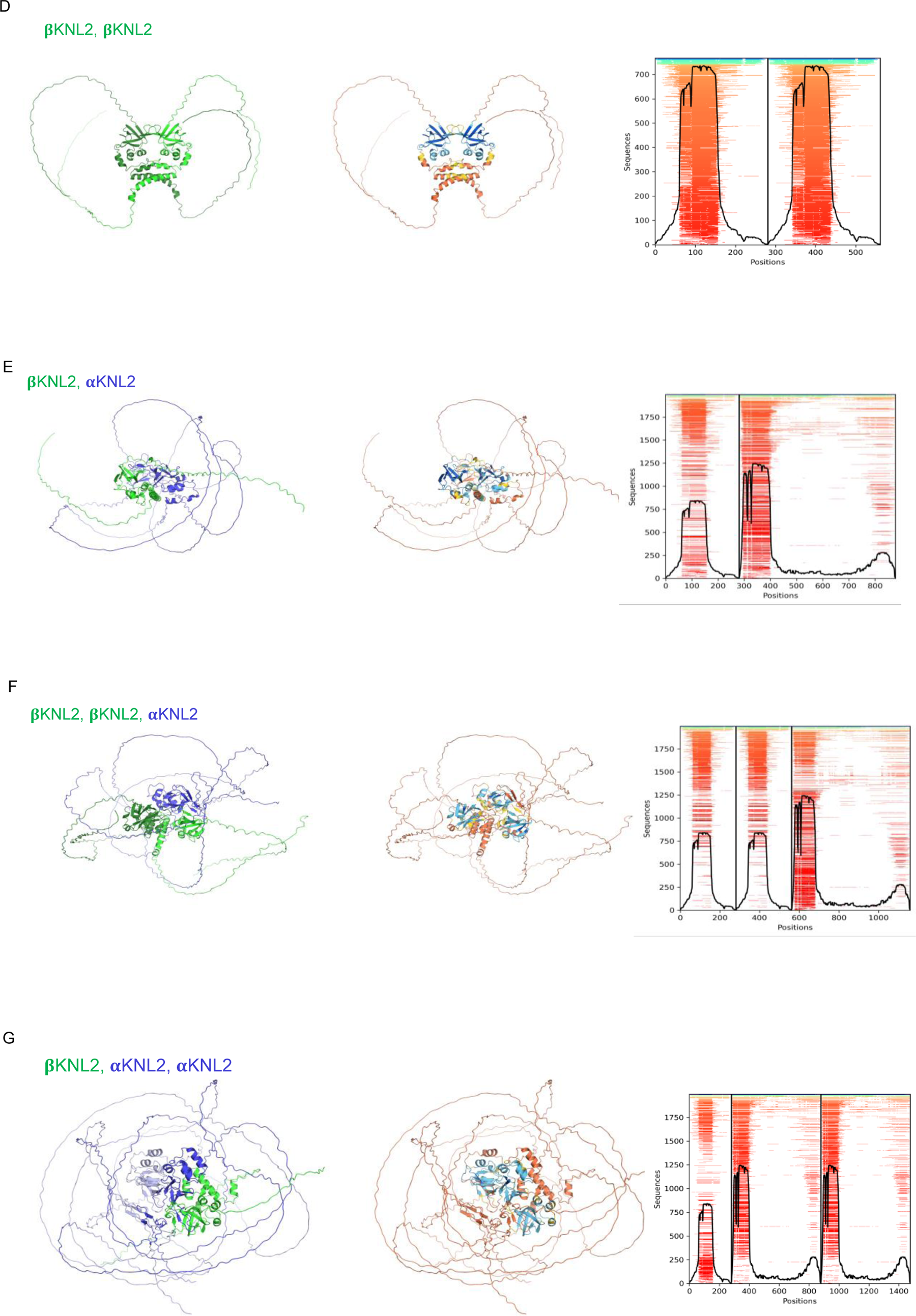
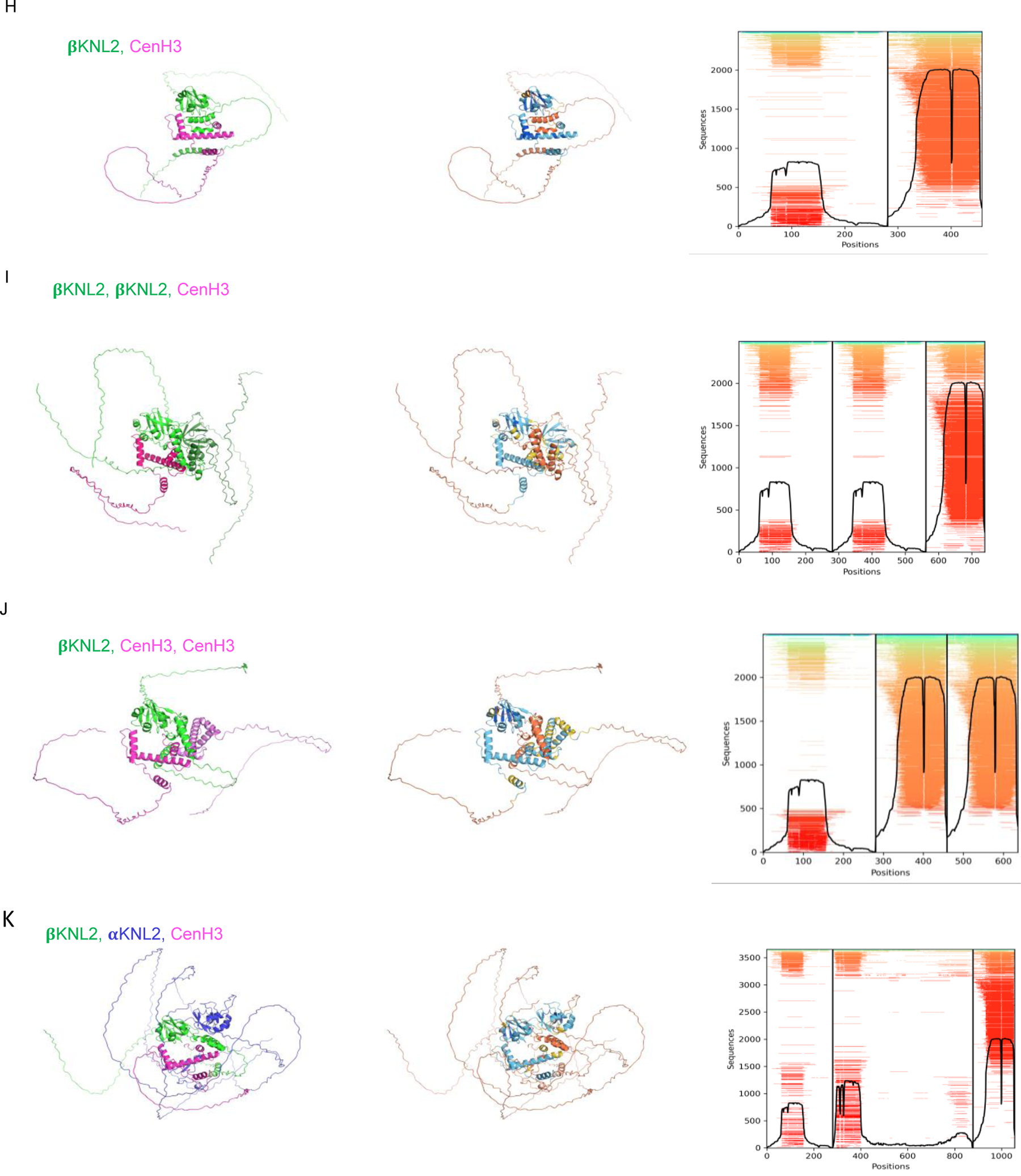
Predicted models of βKNL2, αKNL2, and CENP-A/CENH3 interactions Different combinations of βKNL2, αKNL2, and CENP-A/CENH3 structural complexes predicted by AlphaFold2 are listed in the table. **(A-C)** AlphaFold2 predicted monomeric structures of βKNL2 (green), αKNL2 (blue), and CENP-A/CENH3 (pink), respectively. **(D)** Homodimer of βKNL2, highlighting the symmetrical complex formation. **(E-G)** Various dimer and trimer combinations of βKNL2 and αKNL2. **(H-I)** Different combinations of βKNL2 and CENH3 in dimer and trimer forms. **(K)** Heterotrimer model of βKNL2, αKNL2, and CENH3, showing how these proteins might coalesce in a complex molecular assembly. Each model’s left panels show the monomer colored in above referred color, varying degrees of the same color in case of homodimer. The middle panels, colors each residue according to AlphaFold2 confidence scores, while the right panels illustrate the coverage of multiple sequence alignments for each model.

**Supplementary Figure S3.**
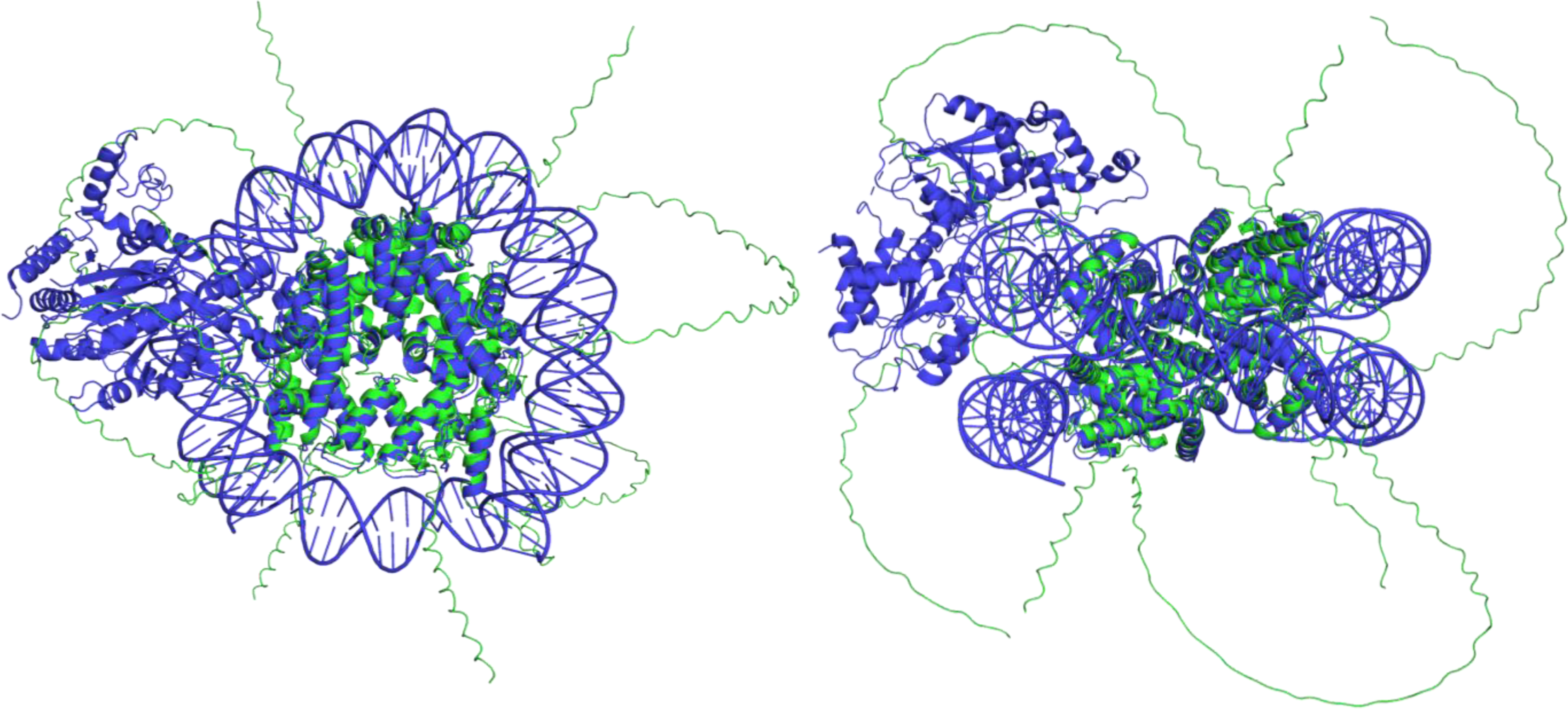
The predicted model of βKNL2 and αKNL2 with the octameric histones (green) superposed with *Arabidopsis* euchromatic nucleosome (blue) (PDB code 7ux9 (Lee, et al. 2023)). The structural alignment was performed with PyMOL, which aligned 391 residues from H2AW6 and H2BHTB9 monomers with the solved nucleosome, with a RMSD of 1.201 Å.

**Supplementary Figure S4.**
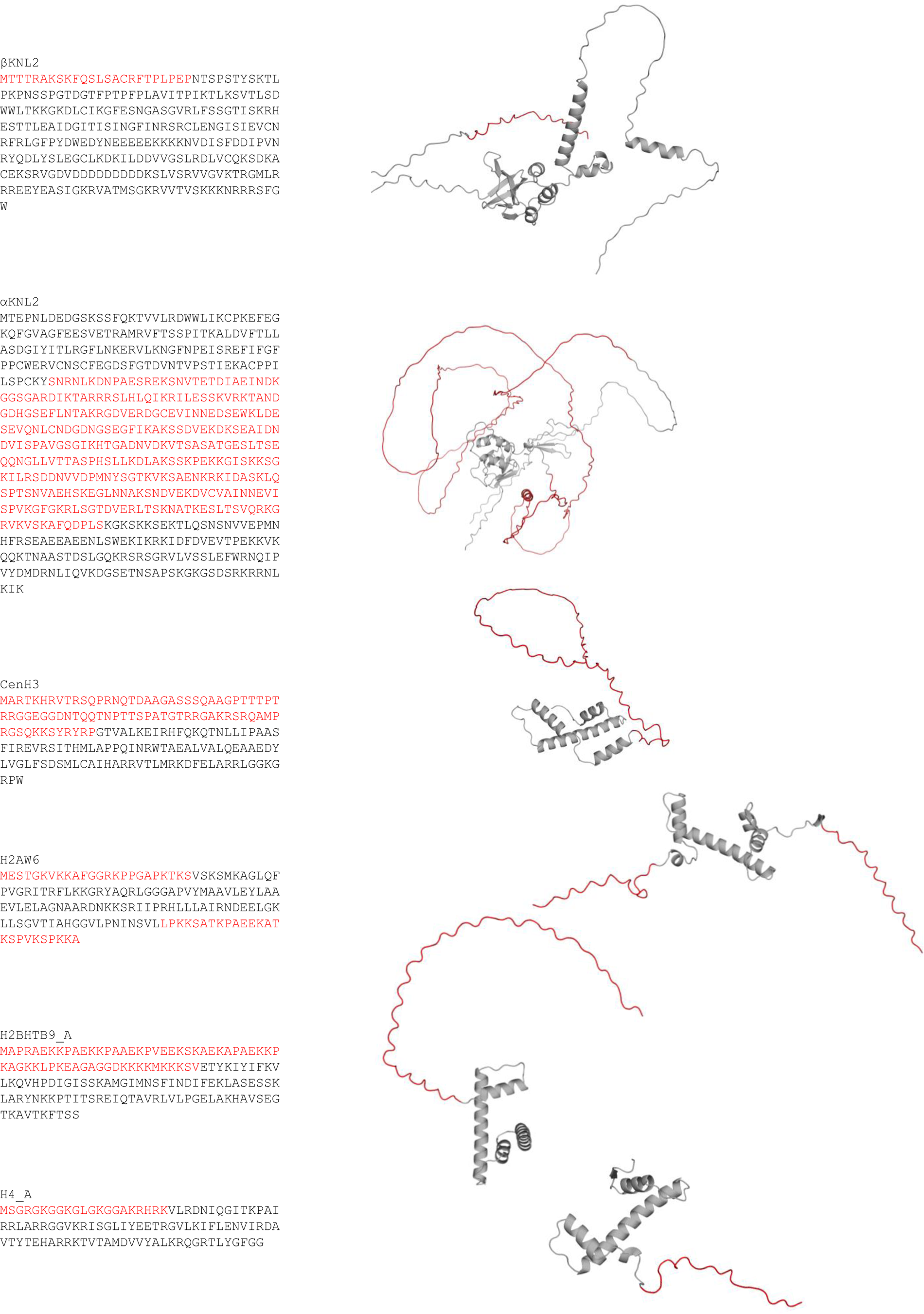
Neglected regions of the monomers in the prediction and modelling of βKNL2 and αKNL2 with the octameric histones. The sequences for each monomer are in the left column, with neglected regions in red, and the correspondent predicted structures are in the right column.

**Supplementary Figure S5.**
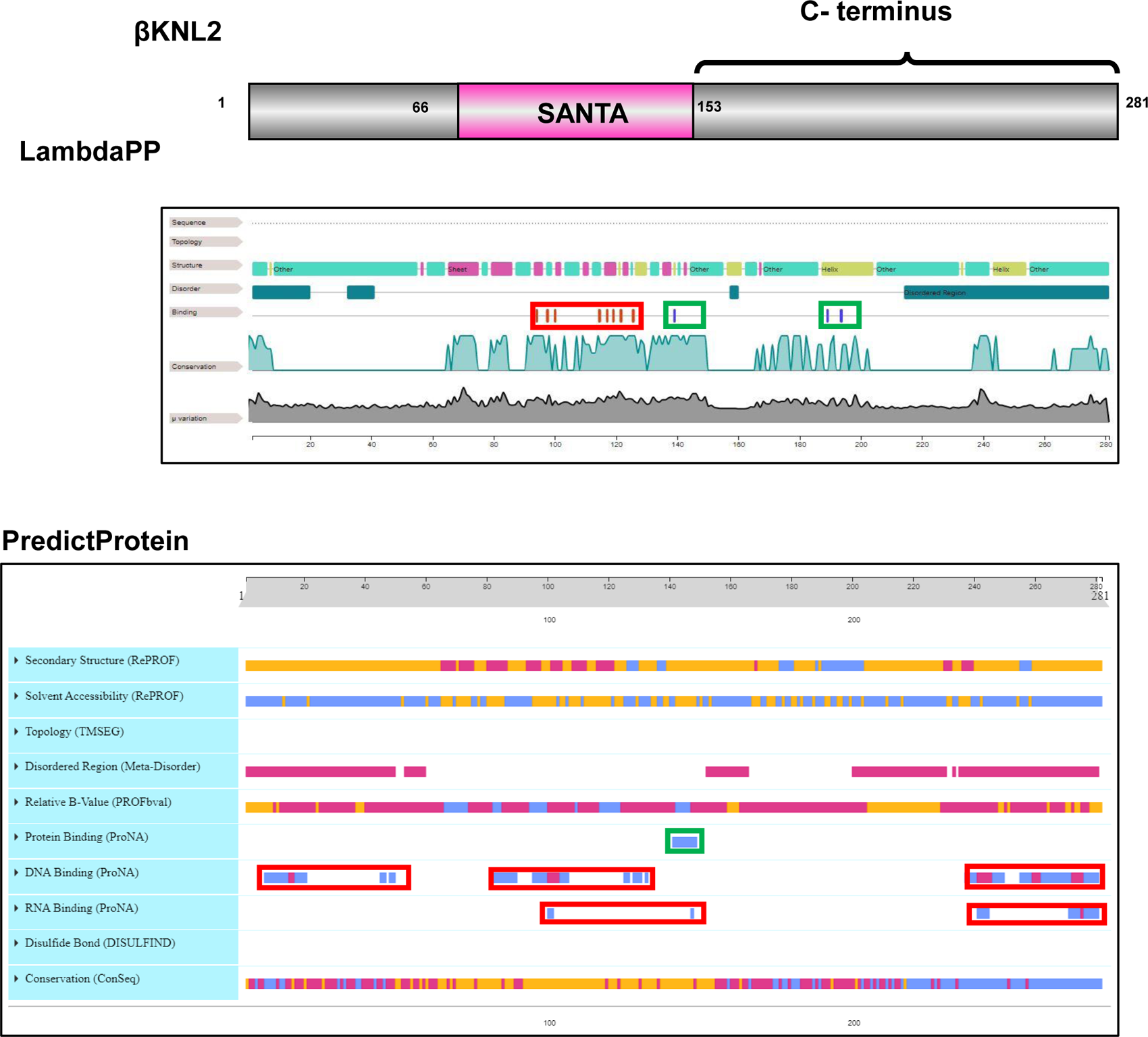
*In silico* prediction of protein-DNA and protein-protein interaction sites of βKNL2 *In silico* analysis of the βKNL2 protein, highlighting potential interaction sites for proteins (indicated in green) and nucleic acids (indicated in red). Computational predictions using LambdaPP and PredictProtein tools suggest that the SANTA domain and C-terminus of βKNL2 have the potential to interact with other proteins and nucleic acids, respectively. Each color-coded box corresponds to regions within the βKNL2 structure predicted to facilitate these molecular interactions.

**Supplementary Figure S6:**
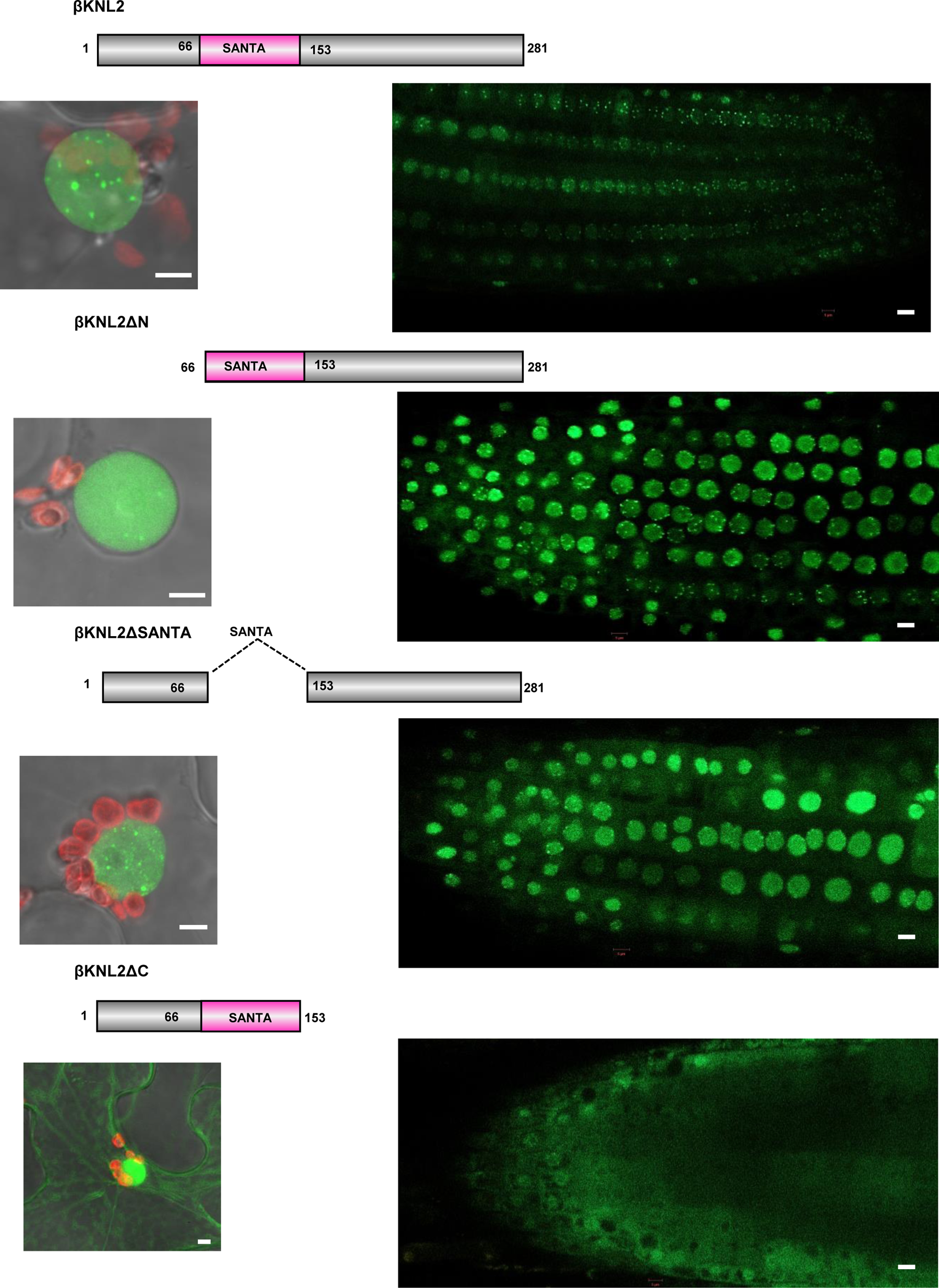
Localization patterns of EYFP-tagged βKNL2 truncated variants in *N. benthamiana* and *A. thaliana* Left panels showing localization patterns of truncated βKNL2 variants when transiently expressed in *N. benthamiana* leaves. The right panels display a representative root tip from *Arabidopsis*, providing a stable expression context from which main Figure 1C was derived. The variants βKNL2ΔN and βKNL2ΔSANTA exhibit centromere-like localization patterns marked by distinct nuclear dots, akin to those observed with the full-length βKNL2. In contrast, the βKNL2ΔC variant shows cytoplasmic localization, indicating a mis-localization pattern compared to the full-length and other truncated variants. Scale bars represent 5µm.

**Supplementary Figure S7:**
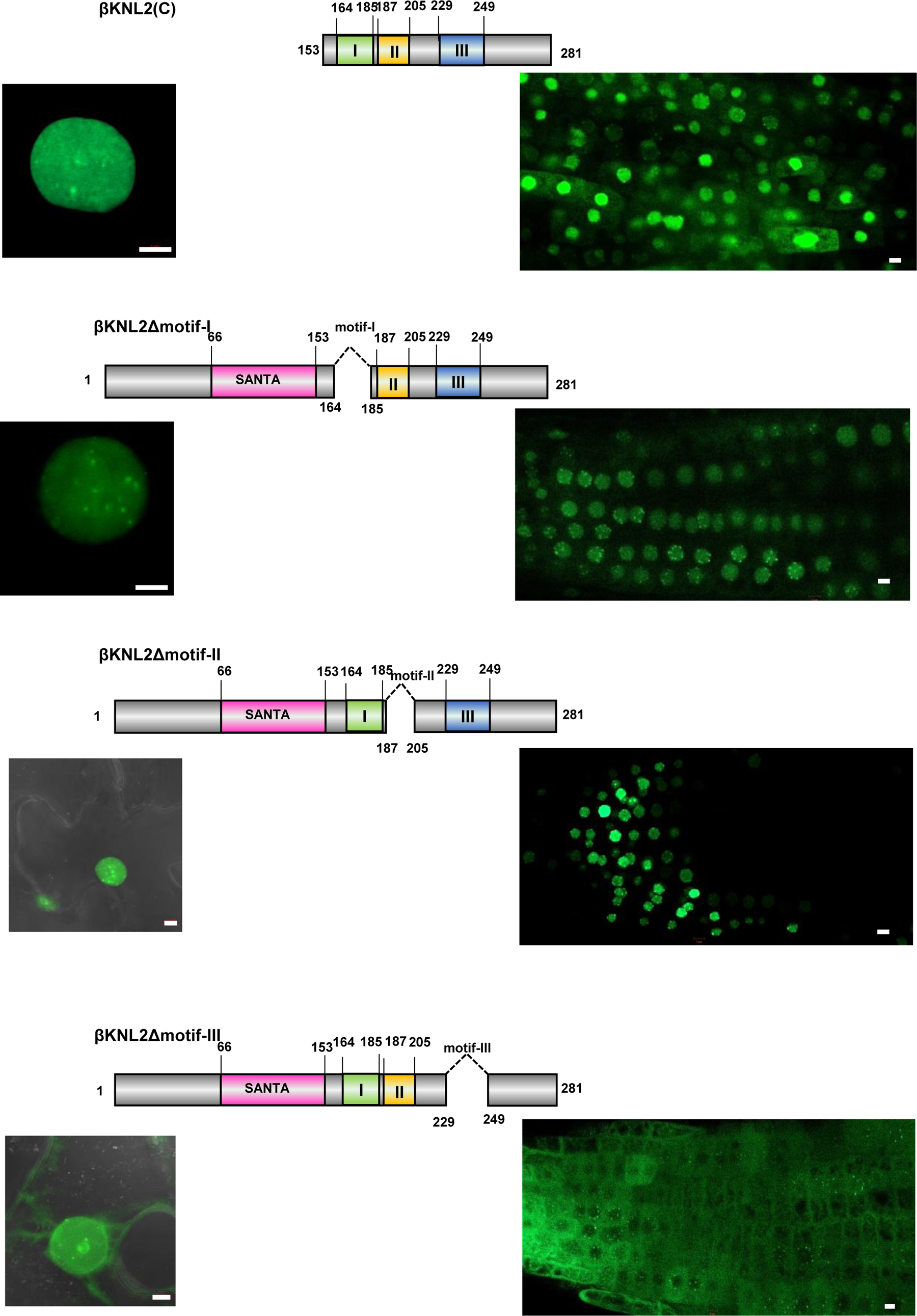
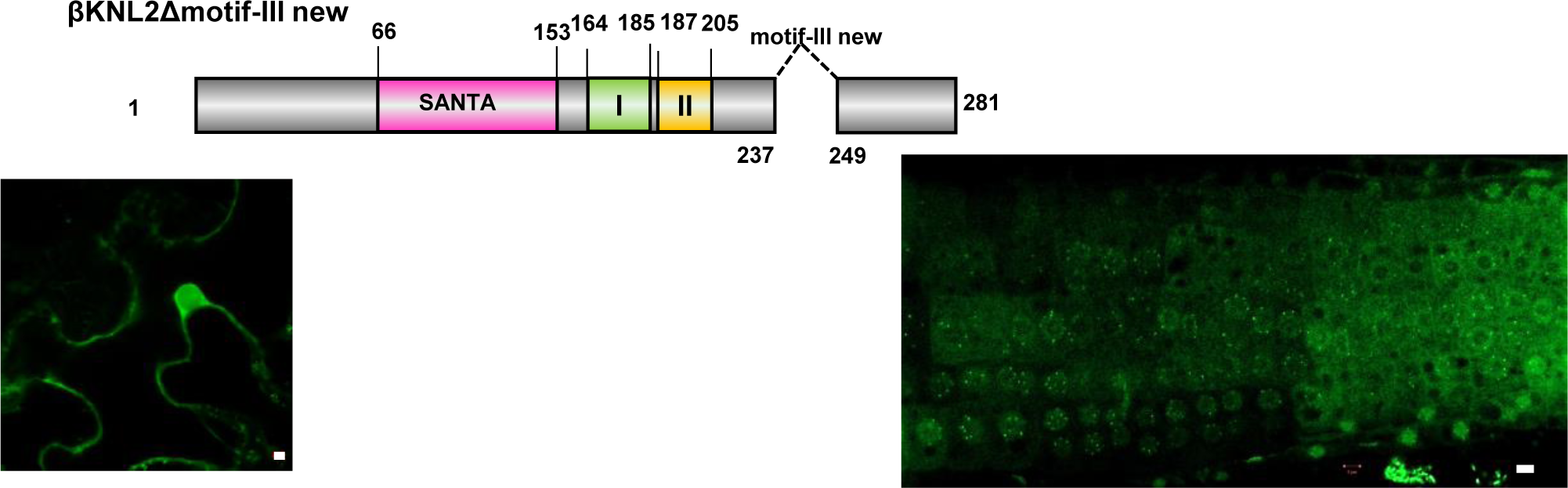
Localization patterns of EYFP-tagged βKNL2 C-terminal truncated variants in *N. benthamiana* and *A. thaliana* Left panels showing transient expression patterns of truncated βKNL2 variants βKNL2(C), βKNL2Δmotif-I, βKNL2Δmotif-II, and βKNL2Δmotif-III in tobacco leaves. The right panel of the figure shows a representative root tip of *Arabidopsis* displaying stable expression of these constructs, from which main figures 2C were extracted. The deletion of motifs I and II retains the localization pattern typical to the full-length βKNL2. In contrast, deletion of motif-III leads to a significant mis-localization of βKNL2Δmotif-III to the cytoplasm. Scale bar: 5 µm.

**Supplementary Figure S8:**
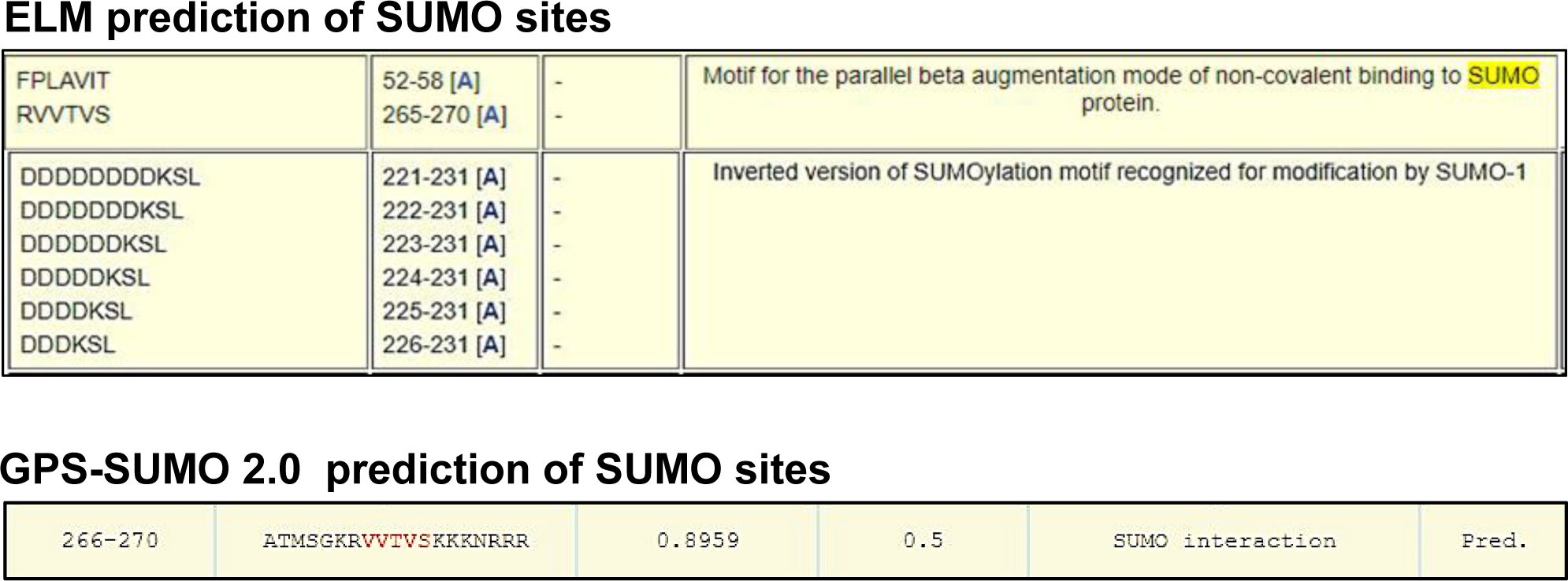
Predicted SUMOylation sites overlap with motif-III of βKNL2 ELM and GPS-SUMO 2.0 bioinformatics tools predict putative SUMOylation sites in a region of βKNL2 spanning residues 221-231. This region overlaps with motif-III, which extends from residues 229 to 249. In addition, both ELM and GPS-SUMO 2.0 identify a SUMO interaction motif spanning residues 266 to 270.

**Supplementary Figure S9:**
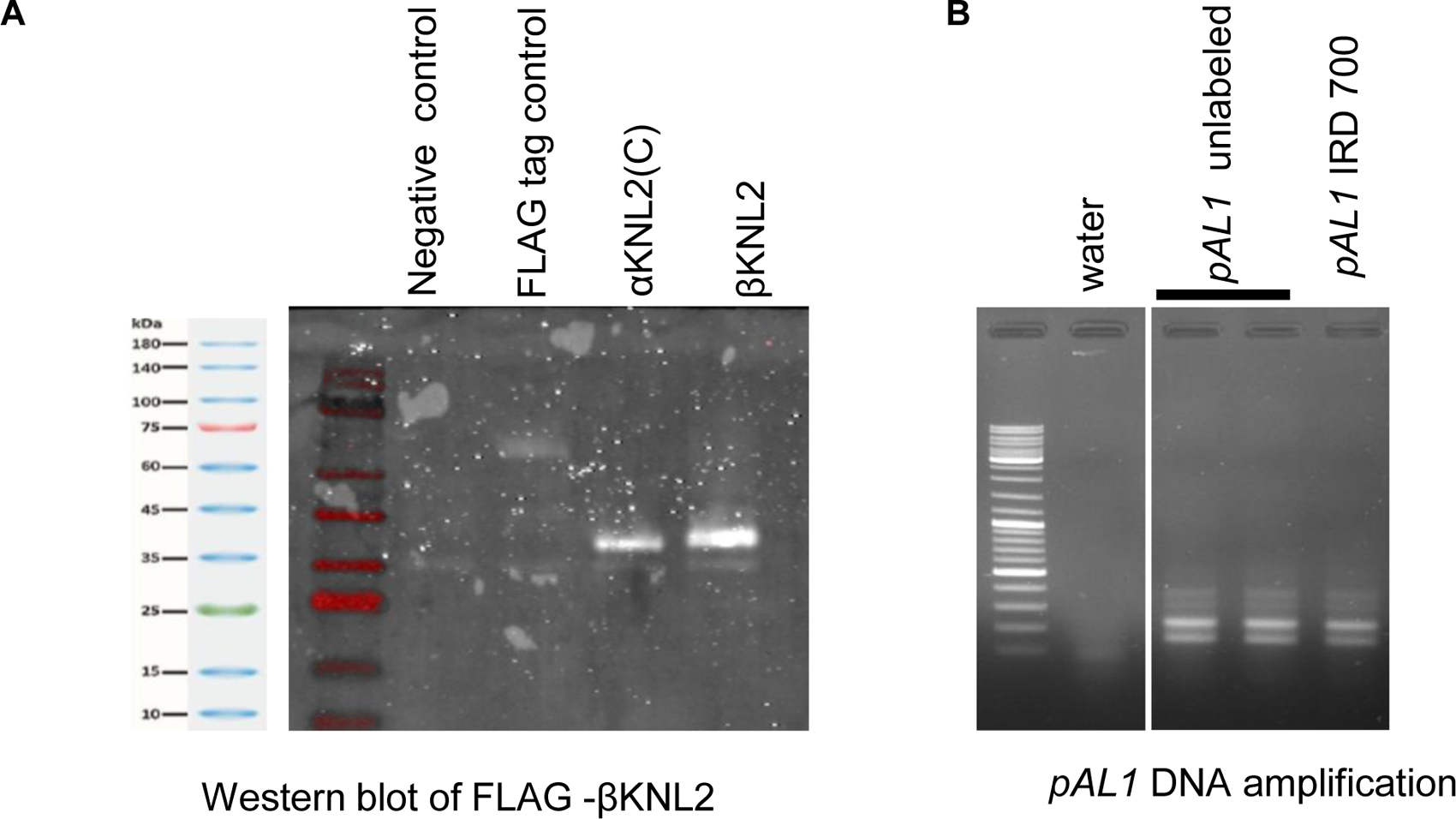
Expression and verification of FLAG-βKNL2 and FLAG-αKNL2(C), and PCR amplification of the *pAL1* DNA (A) Western blot analysis confirming the expression of FLAG-tagged αKNL2(C) and βKNL2 proteins in TNT SP6 High Yield Wheat Germ system (Promega) using anti-FLAG antibodies. Despite their different expected sizes, αKNL2(C) (26.36 kDa) and βKNL2 (31.56 kDa) proteins exhibit similar migration patterns, which may be attributed to post-translational modifications. (B) The PCR amplification of the *pAL1* centromeric repeat from *Arabidopsis* genomic DNA.

**Supplementary Figure S10:**
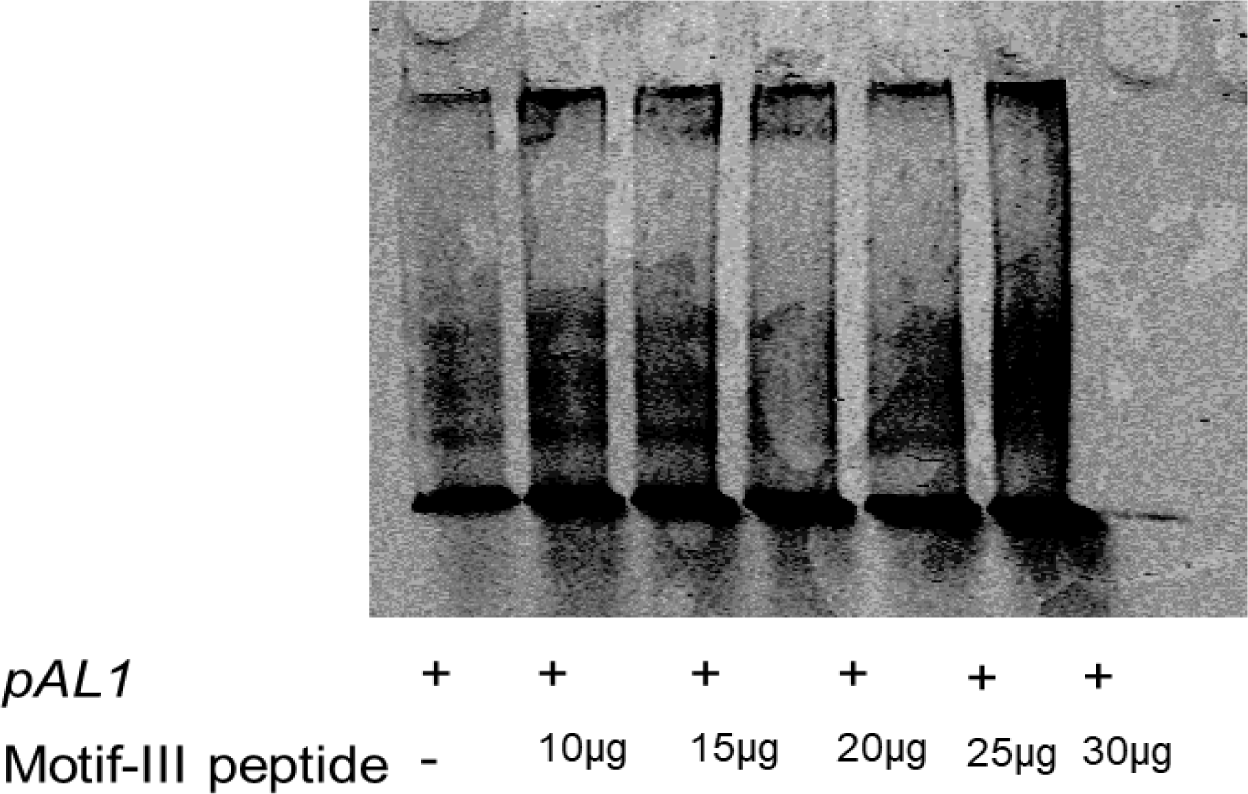
Motif-III of βKNL2 can bind to centromeric DNA Increased upward shifts correlate with increasing concentrations of motif-III peptide (10µg to 30 µg) in IRD700-labeled *pAL1* DNA, demonstrating dose-dependent binding.

**Supplementary Figure S11:**
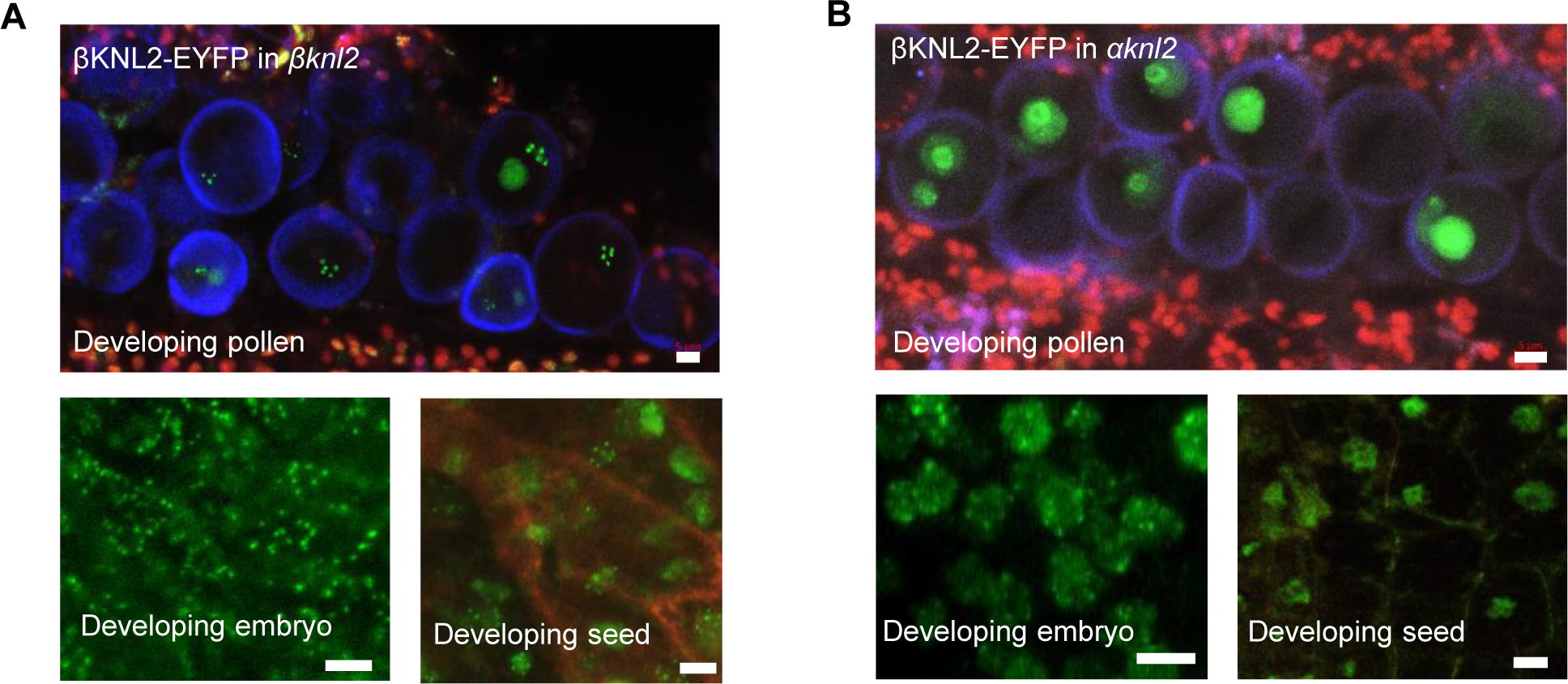
Disrupted centromeric localization of βKNL2-EYFP in *αknl2* mutants **(A)** *Arabidopsis βknl2* transformants expressing βKNL2::βKNL2-EYFP serving as controls showed βKNL2-EYFP signals localized in both the centromeres and nucleoplasm within the nuclei of developing pollen, embryos, and seeds. **(B)** In *αknl2* mutant transformants, βKNL2-EYFP was primarily observed in the nucleoplasm of developing pollen and seeds. Notably, in developing embryos, βKNL2-EYFP targeted to the centromeres, similar to *βknl2* mutant transformants. Scale bar: 5µm.

**Supplementary Figure S12:**
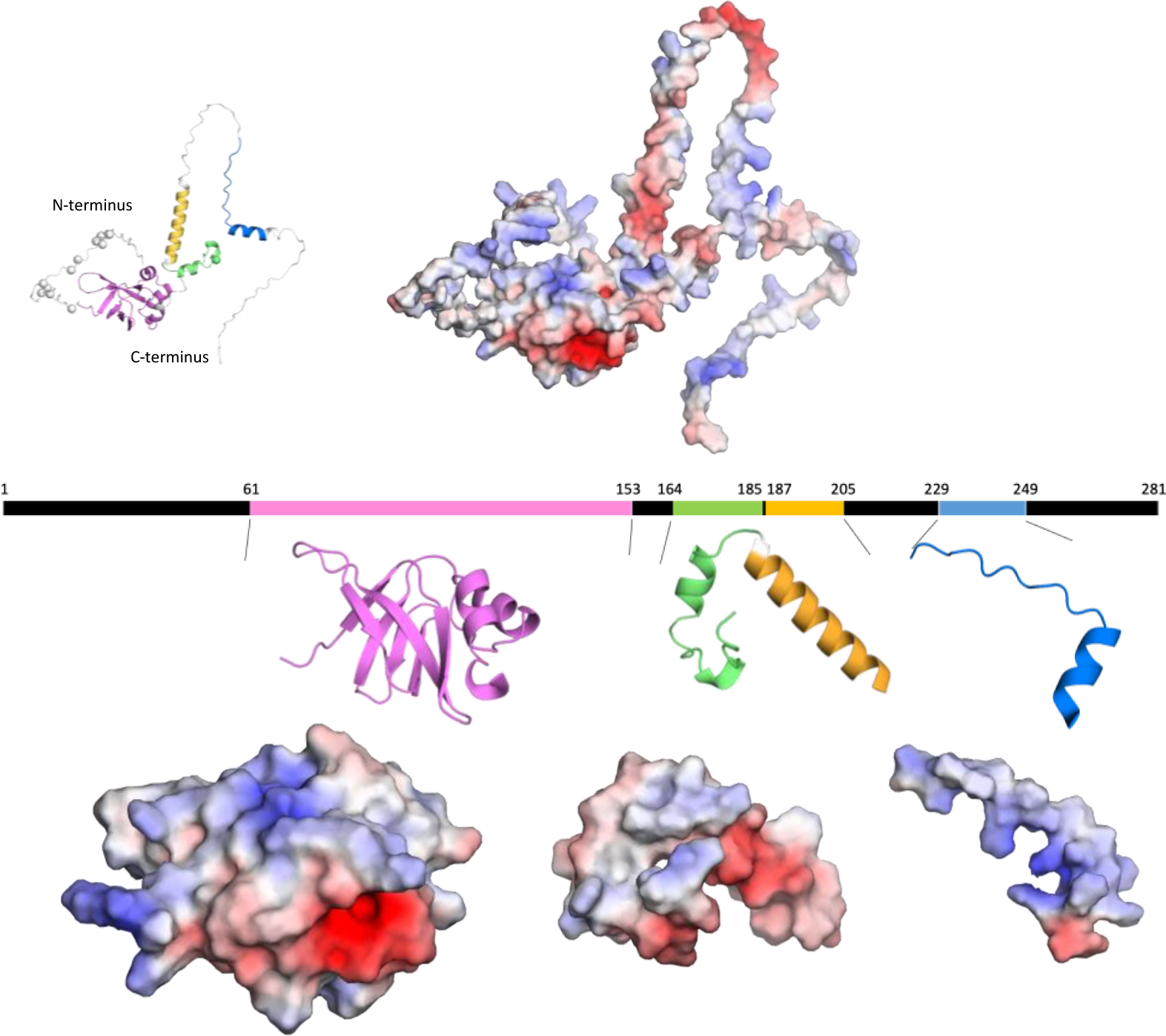
Surface charge distribution of the SANTA domain and motifs I, II, and III of βKNL2 Illustration of the surface charge distribution of the SANTA domain and motifs I, II, and III in βKNL2, emphasizing the electrostatic properties critical for protein interactions. Top: Display includes βKNL2 in cartoon mode on the left and surface mode on the right, showing the full protein with charge distribution. Middle: Schematic representation highlights the relative positions of the SANTA domain (magenta) and motifs I (green), II (yellow), and III (blue) in cartoon mode. Bottom: Corresponding surface views of the SANTA domain and motifs I, II, and III, color-coded to show charge distribution from negative (red) to positive (blue). Notably, the SANTA domain and motif III exhibit a predominantly positively charged surface, suggesting potential roles in DNA or RNA binding through electrostatic interactions.

**Supplementary Figure S13.**
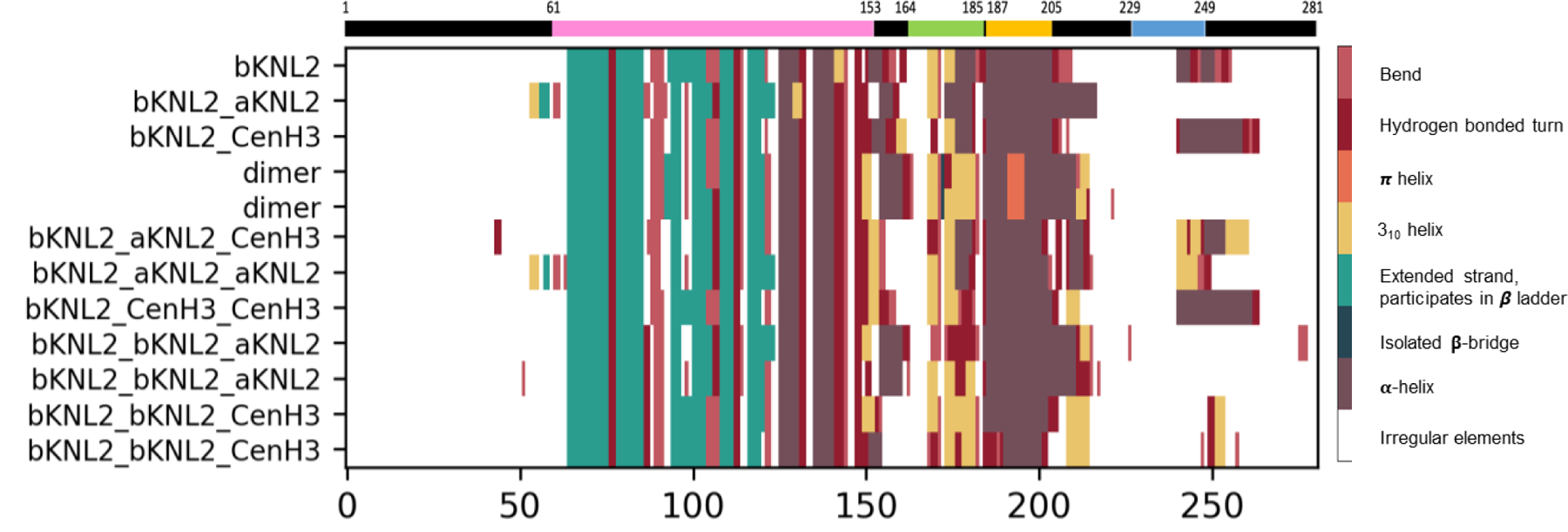
Structural adaptability in βKNL2 C-terminal motifs This figure presents a DSSP (Define Secondary Structure of Proteins) analysis detailing the structural adaptability of βKNL2’s C-terminal motifs when interacting with itself and with kinetochore proteins αKNL2 and CENH3. The analysis, based on models generated by AlphaFold2 (referenced in Supplementary Figure S8). The homodimer of βKNL2 exhibits changes in secondary structure formations at the end of the SANTA domain and within C-terminus motifs I and II. Similar structural changes are observed in heterodimer and oligomer formations with αKNL2. Combinations of βKNL2 with CENH3 show modifications in motif-III. These findings highlight the dynamic nature of βKNL2’s C-terminal motifs, demonstrating their ability to adapt structurally according to different binding partners.

**Supplementary Figure S14:**
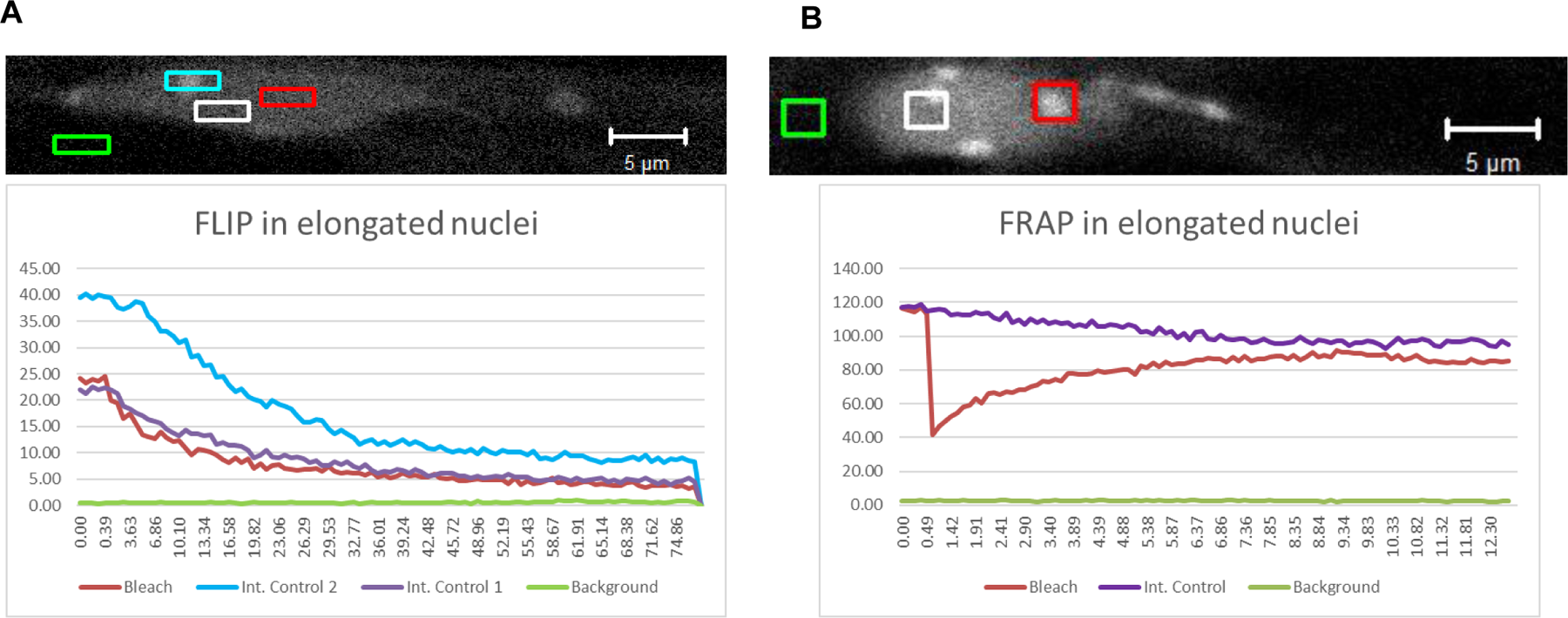
FLIP and FRAP experiments unveil the dynamic nature of βKNL2-EYFP FLIP and FRAP experiments were conducted in elongated non-meristematic nuclei of *A. thaliana* transformed with a 35S::βKNL2-EYFP construct. (A) For measuring FLIP, individual nuclei were scanned three times with a 488 nm laser line (2,5% laser power, scan speed 6 without averaging) followed by repeated bleaching of a region of interest within the nucleus measuring 1,4 μm2, using 100% laser power and 4 iterations alternated by single recordings. (B) For measuring FRAP, individual nuclei were first scanned three times with a 488 nm laser line (2.5% laser power, scan speed 6 without averaging). Following, a region of interest measuring 1.4 μm2 was bleached by more than 50% and followed by 50 additional scans. Each experiment was run over the time scale of 40 sec, repeated 5–10 times and results averaged. In case of βKNL2 fluorescence intensity at chromocenters recovers to about 80% within 10 sec. For both FLIP and FRAP experiments fluorescence intensity was measured in the area of bleaching (Bleach) as well as in adjacent regions. The x axis represents the time scale of the experiment in second, the y axis corresponds to fluorescence intensity (arbitrary units).

**Supplementary Figure S15.**
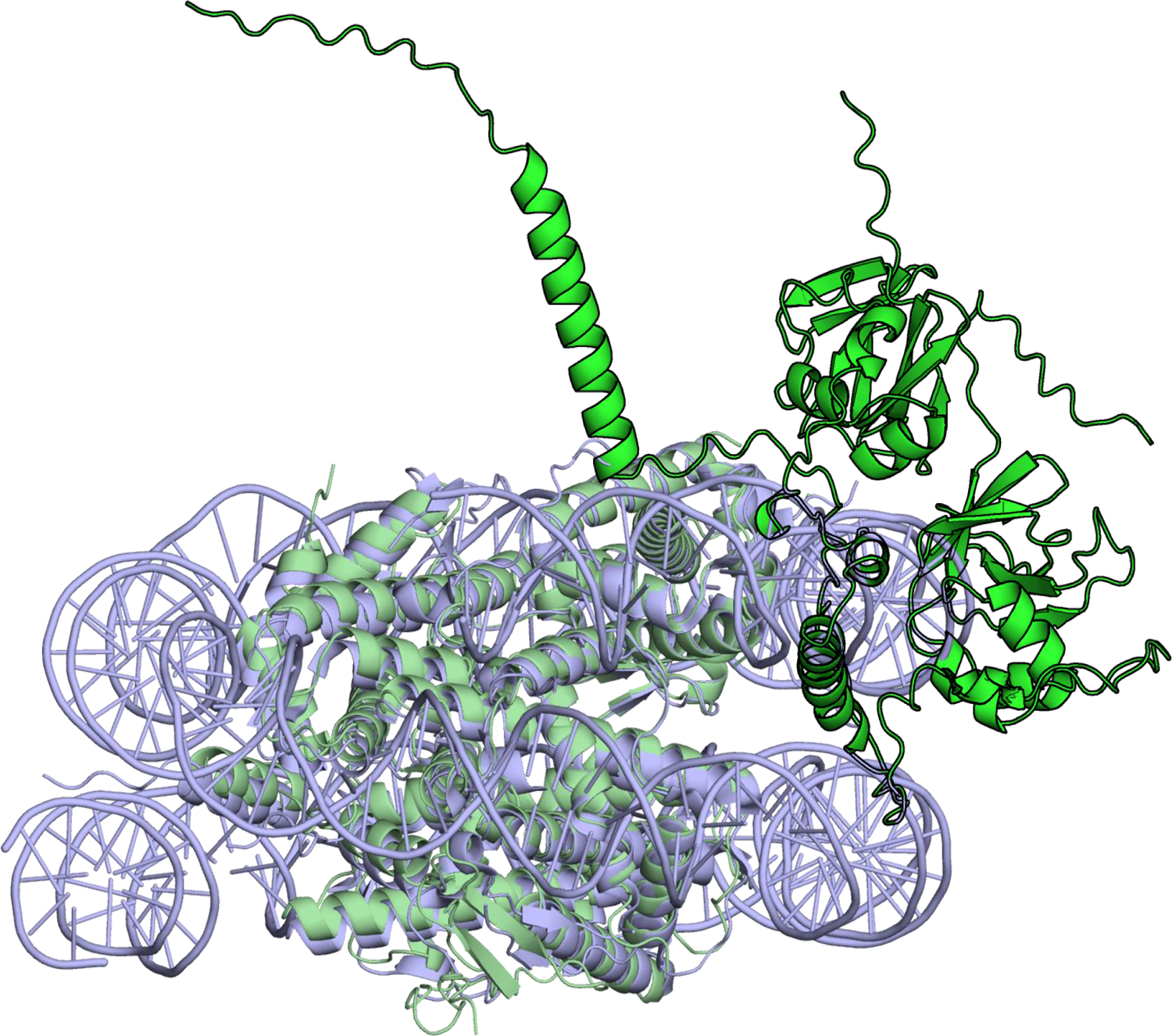
The predicted model of βKNL2 and αKNL2 with the octameric histones (green) superposed with *Arabidopsis* euchromatic nucleosome (blue) (PDB code 7ux9 (Lee, et al. 2023)). The predicted βKNL2 and αKNL2 are highlighted with stronger green color and black contour, evincing its clash with the DNA.

**Supplementary table_S1:**
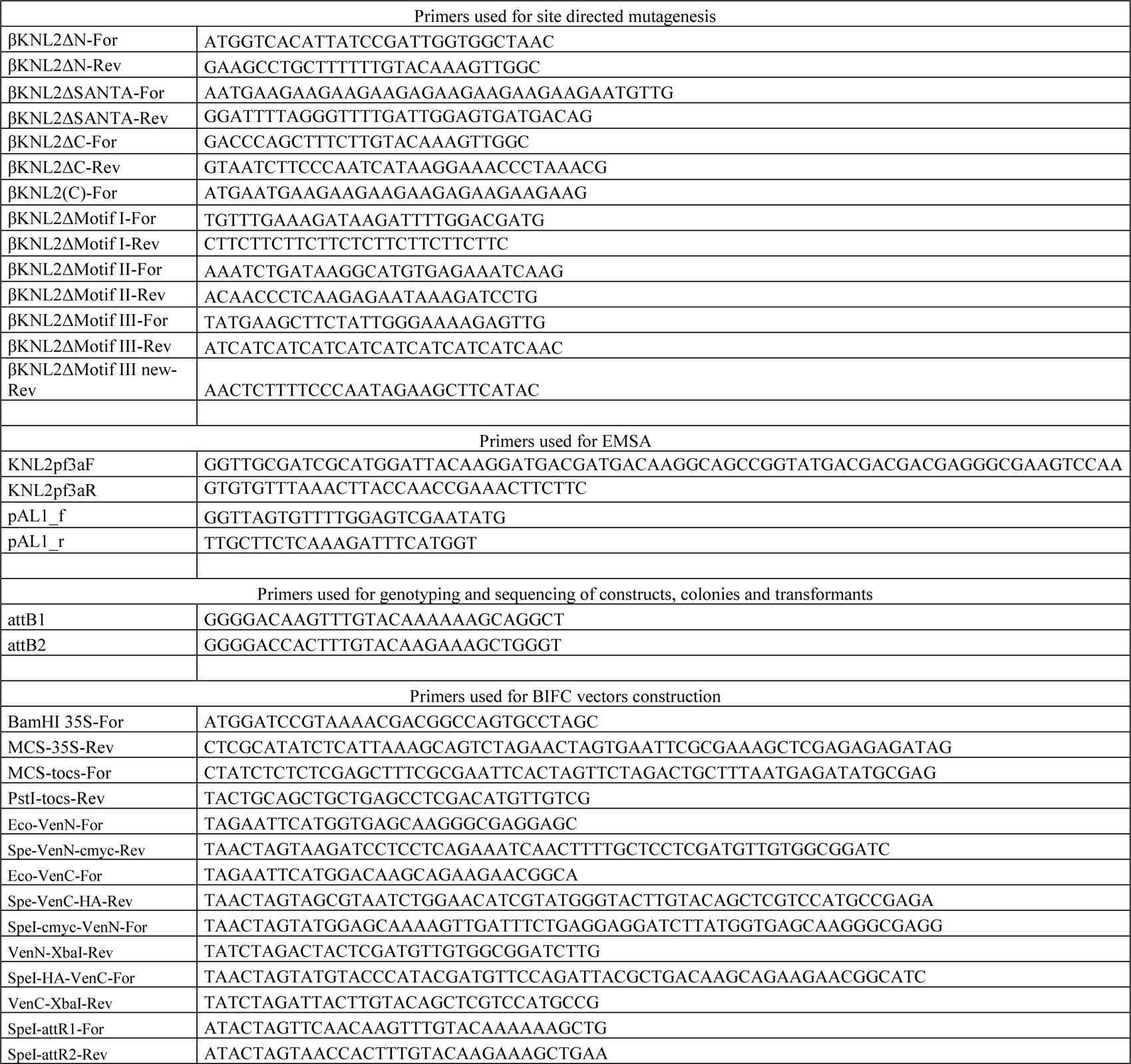
List of all primers used in the study.

