## Supplemental File.2 for "Structural Basis of βKNL2 Centromeric Targeting Mechanism and Its Role in Plant-Specific Kinetochore Assembly"

| Name | *p*-value |
|---|---|
| Ceratopteris_richardii_gene01725 | 6.88e-41 |
| Anthoceros_angustus_AANG012187 | 1.24e-41 |
| Azolla_filiculoides0019.g015039 | 6.09e-49 |
| Azolla_filiculoides0032.g024707 | 8.16e-32 |
| Marchantia_polymorpha_0034s0011.1.p | 7.36e-36 |
| Physcomitrella_patens00338G00350 | 3.26e-32 |
| Picea_abies00016756 | 1.58e-50 |
| Picea_glauca00028859 | 7.89e-53 |
| Picea_sitchensis00004746 | 1.36e-53 |
| Pinus_pinaster00061713 | 8.02e-57 |
| Pinus_sylvestris00003294 | 4.96e-51 |
| Pinus_taeda00042871 | 9.30e-41 |
| Pseudotsuga_menziesii00073202 | 2.40e-56 |
| Salvinia_cucullata0025.g009355 | 2.19e-37 |
| Salvinia_cucullata0183.g024952 | 4.96e-35 |
| Aegilops_tauschii2Gv20621700.1 | 4.55e-38 |
| Aegilops_tauschii2Gv20622700.1 | 1.63e-31 |
| Amaranthus_hypochondriacus008009-RA | 8.91e-28 |
| Amborella_trichopoda_00137.13 | 5.64e-35 |
| Ananas_comosus021414.1 | 3.54e-55 |
| Aquilegia_coerulea1G467000.1.p | 8.55e-33 |
| Aquilegia_coerulea1G467200.1.p | 1.70e-43 |
| Brachypodium_distachyon_3g30800.1.p | 3.86e-37 |
| Brachypodium_distachyon_4g44060.1.p | 2.35e-51 |
| Brachypodium_distachyon_5g25291.1.p | 2.80e-53 |
| Brachypodium_stacei_03G173200.1.p | 1.89e-34 |
| Brachypodium_stacei_05G288200.1.p | 2.42e-49 |
| Brachypodium_stacei_09G247600.1.p | 1.87e-41 |
| Citrus_sinensis_1g023837m | 1.29e-64 |
| Citrus_sinensis_1g045557m | 5.84e-44 |
| Cucumis_sativus_094610.1 | 8.65e-61 |
| Daucus_carota_025649 | 8.13e-40 |
| Elaeis_guineensis_00052_p0043.1 | 1.95e-48 |
| Eucalyptus_grandis_I00627.1.p | 4.59e-49 |
| Fragaria_vesca_22410-v2.0.a2-hybrid.t1 | 1.73e-48 |
| Glycine_max_09G003000.1.p | 2.42e-27 |
| Glycine_max_10G043400.3.p | 1.67e-66 |
| Glycine_max_13G131100.2.p | 1.97e-64 |
| Gossypium_raimondii_004G087600.1 | 1.25e-58 |
| Hordeum_vulgare_2Hr1G067110.2 | 7.22e-48 |
| Hordeum_vulgare_2Hr1G067130.1 | 4.74e-27 |
| Hordeum_vulgare_2Hr1G073590.2 | 5.16e-34 |
| Lactuca_sativa_2_36541.2 | 1.08e-45 |
| Manihot_esculenta_08G078100.1.p | 5.91e-68 |
| Nelumbo_nucifera_010241510.1 | 1.57e-48 |
| Oryza_sativa_01g34610.1 | 3.10e-31 |
| Oryza_sativa_04g28040.1 | 1.65e-44 |
| Populus_trichocarpa04G07690 | 3.44e-26 |
| Populus_trichocarpa07G12430 | 3.03e-52 |
| Populus_trichocarpa17G03610 | 2.71e-68 |
| Setaria_italica_K96788 | 2.12e-32 |
| Solanum_lycopersicum_04g015710.3.1 | 4.94e-63 |
| Solanum_lycopersicum_09g010550.3.1 | 4.83e-45 |
| Sorghum_bicolor_KXG30438 | 1.36e-30 |
| Sorghum_bicolor_OQU89324 | 3.21e-31 |
| Spirodela_polyrhiza_6G0070900 | 3.84e-29 |
| Triticum_urartu_13040-P1 | 2.20e-19 |
| Vitis_vinifera_205s0124g00540.1 | 1.16e-65 |
| Zea_mays_001145421.1 | 5.63e-33 |
| Arabidopsis_halleri_0330s0008.1.p | 1.48e-37 |
| Arabidopsis_lyrata_2G18170.t1 | 1.06e-124 |
| Arabidopsis_thaliana_1G58210_new | 2.93e-123 |
| Arabis_alpina_G656800.h1.t1 | 2.53e-110 |
| Barbarea_vulgaris-Contig491-snap0.32 | 2.57e-161 |
| Boechera_retrofracta_RETT00031638 | 8.56e-168 |
| Boechera_stricta_30057s0098.1.p | 4.61e-163 |
| Brassica_cretica_RQL80743.1 | 2.47e-140 |
| Brassica_juncea_A040432 | 2.60e-143 |
| Brassica_juncea_B007540 | 6.44e-119 |
| Brassica_napus_GSBRNA2T00024219001 | 9.95e-176 |
| Brassica_napus_GSBRNA2T00086338001 | 9.89e-143 |
| Brassica_nigra_B045991-PA | 4.67e-122 |
| Brassica_oleracea_1g087890.1 | 5.08e-143 |
| Brassica_rapa_A01g027290.3C | 2.82e-144 |
| Camelina_sativa_XP_010414757.1 | 4.19e-107 |
| Camelina_sativa_XP_010470274.1 | 1.02e-102 |
| Camelina_sativa_XP_019101020.1 | 1.01e-113 |
| Capsella_rubella_0002s0640.1.p | 3.02e-41 |
| Cardamine_hirsuta_CARHR053810.1 | 5.64e-126 |
| Cardamine_hirsuta_CARHR210840.1 | 4.56e-53 |
| Conringia_planisiliqua_CP11_g33626_DN3_SP0_c | 1.82e-32 |
| Euclidium_syriacum_0020s0140.1.p | 2.42e-45 |
| Euclidium_syriacum_0048s0039.1.p | 9.54e-92 |
| Eutrema_heterophyllum_scaffold455_cov155.49_1 | 2.19e-129 |
| Eutrema_salsugineum_v10023956m | 7.63e-129 |
| Eutrema_yunnanense_scaffold81_cov138.25_1 | 3.49e-137 |
| Lepidium_meyenii_scaffold1913.53 | 6.91e-129 |
| Lepidium_meyenii_scaffold277.39 | 2.06e-129 |
| Lepidium_meyenii_scaffold668.120 | 2.17e-48 |
| Lepidium_meyenii_scaffold96.171 | 1.39e-133 |
| Moringa_oleifera_10005842 | 2.80e-58 |
| Moringa_oleifera_10015199 | 1.40e-43 |
| Raphanus_raphanistrum_RrC35_p2 | 3.58e-133 |
| Raphanus_sativus_018437333.1 | 4.16e-127 |
| Schrenkiella_parvula_Tp2g05410 | 3.29e-171 |
| Sisymbrium_irio_maker2387.1 | 2.61e-159 |
| Tarenaya_hassleriana_010524136.1 | 3.23e-75 |
| Thlaspi_arvense_17942 | 5.54e-141 |

Motif Locations

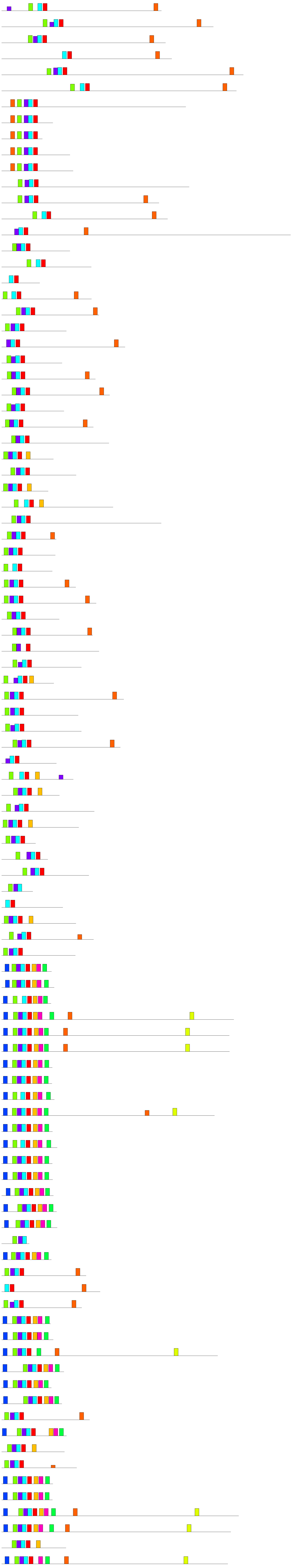

| Motif | Symbol | Motif Consensus |
|---|---|---|
| 1. | | NGFSSEVCNRFLLGFPYBWEDY |
| 2. | | RHDSTTLETSDGITVLJSGFIN |
| 3. | | TSSGTLKSVTLSDWWLTKKEKE |
| 4. | | VGGFESKGGSEVRLFSSAPISK |
| 5. | | FGVSFDDIPVNRLQDLLFTASP |
| 6. | | ESLVSSVVGVKTRGMLRRREEG |
| 7. | | LSARRSSPRTRSGALPEPNSSP |
| 8. | | KILDDVVDSLRDLVCPKTEKED |
| 9. | | SRSGRLLVPPLAHWRNQIISYB |
| 10. | | FWLRFSTSVHQIQKYQTTVQDL |
