## Supplemental File.1 for "Structural Basis of βKNL2 Centromeric Targeting Mechanism and Its Role in Plant-Specific Kinetochore Assembly"

>Ceratopteris_richardii_gene01725

MATHGLRLDASARNRFCSDVTHTENYCNHCCGHHSLSSPTRSKFSSPEISWVHQQKQFARDPDDAFRRGFCSARYFVQSSVQPPTPHQDYEFLYPASTSMSEGTPGGASRSFEPCSSSAWLSSARGFNLERNSEREDLCTIGYETCSVQSNQVTLYSWYLIKCEGASDSVGVSVGGFLGTGGIEKFESASIVQRMEKCILMAQDGVKIRLAGLINGSTSVANGFSQNLVESFLSGFPYTWNHLLYSELRASKMKQPDHQGFHEGLSETVEVSQLSFKGSHADQLKEIPFSNQVKDHHGYPEVLAEIELYAKQTSRKIQIEDKNVEFSNQTVSTDNDDFKQGLSKNDVVMHSEMGASLQQCSLDASLTNIEMKGKNEELNNQAITTDHGNSEIGTFKDDLLITSDTVTPLEKDILNSSHSIEKDLEASRYNGQFTDHVRSNSRRTSKAQSPHTRKKGGTVNNQSCVGSSPAIQTNQCTRMTRSMHKVSMHQDHSKNNDSLLPDRVDNGGNAGSYSGKNVAGDCHRQASFTMSQKNNKGKSDHTKTLNTTSCASKDGDEDITQETESKLNVDANLDPKPVCLEKVVTISSDKNDDDKDSDRQDMPMLARKSETNANVECSTLKANIKKHRKSNKESDVAVKDPGQITRHLEKEVGTASSDENDDDKDHDRQDTPMSVRKSEANANMEGSTVKVNTKKHRKSNKEYDVAVKDPGQATRHLKKEVVTVASDKNDDDKDPDRQDMRTSVRKSEANANMECSTVKVNTKKHRKSNKECGVAVKDPGQSTRHLDILPALPSPRVPVPKQLVSEAFGLKTSRSGRLLVPPLAHWCGQSLLRDKDGGLLQYLMVQKTRKIIL

>Anthoceros_angustus_AANG012187

MLERGGDGVPPCGPRGRQWSSRLPLFTKPSSRCAGLRPHAPAQHDRRRDACSREFDCGEDSYDAFRQQQLEPQPEHQQHQQNQHCGFLEQSGYDEAEGMHGSARAPHSPFERRGEGCRCWSCSSRVRFRSEDSEGKSPEEHRSWRRDAGRRPKECEHDGCRCRAGLRGRGSVSPCQSDSFRHFSRLGPNTPSDTSTSGSGRQGLDELRARDVPLMAWHGRQRPAPAEKQVSLRRWFLVRCDDKEAEDVVDATNPRVAVGGMHTSESFELQTVRSTMIVERIEKSRLRTKDGVEIRLMGCIDAETTIANGFSVKVVHHFLTGFPFVWKHFLELFLSNDTTPKSDIPLYLPETPVTPDAAATGFKFGGREGVSCRGVGSNEVESPKTVAILSPLPATPTPATQTCVESTETQELDNLNIVAAMLSPAPATPTTCLESTEKEELENPNVVATSSPLLGAQVLEDLGCHDAHLDRQEETGACTSQLKGKRSRKGRVPPVRVYKLRSSAGELEHGLPDRLGTTRKRGVKRQRDIDTATGYRIMDDNDEGPAGGSSTNGKEKEDQVERSGNNSCGGLERQMEAPAGASEGSDLEKNIGPSSTALEKDGRERVVRPAEGELSNCDILIHVNEVEVEQPPKARGNCMGGVELQIEAHMVHTEVSTNLETLGPSIIASSKDEIGLEDHLVTQLEDEAQLEEMLLDRELQEERTGDGEEEDRTELVGVHPDIEQGDSRELEEVHPGRQEEDGRVLEEVPRDMQQDAGRGLDEVPLDIQRDATSEVEDLHVHIQQNAGTELKDLHLDIAQYAGIELKELHFDVAHDADRELKDVCLDRLDKVGRELDEVRPDRGIEDGIEFEELHPDRQGASGEELGEVRTDLEQEDGSHLEEVHPDNGWEGGLALVVFHDRGSEGGAKPEEHPHAGKITSSNELGDSHPEVGGRKGVKRPREETFDAPKSAPTEDIDGVDSAGSDKVSRTGNRYTRRGSKKRNCKVTPKNPTIERKSDMQKQLASSGCQSRRSKTSMIPPPLPTPCVSVTTDLVSQAYGLKISRSGRLLVPPLAYWRNQTTARDKDGGIIAIMDGFDENETPDTGCFNFEPPKEKRAVKLQRQLQQAAEERQQVTRAKASKKRRQAVAK*

>Azolla_filiculoides0019.g015039

MEALLTTIFQHHQQNHIPSGSPNFLKPTEIHQSFRQRGPSDLCFSSTPSYPSEERRRRISSGRSSEPSFSKFSASEGITAYNVSENAYIPRTPQTEDFGMASSSPYVQPASNAKFIPSAYTQPRNCERQDFRMASAVNGSDHHMTAPQKQVKLYRWYLIKCGTSSVEIAVGGLLSPGTVGSRRQEMAAITKRNDKCRLMTKDGVEVRLTGLIDDCTTIANGFSVEIVHLFLSGFPYTWNHLLQNEFQTCTIKNPICLGSGTDALNSIETSELNFEGCANKEKCLPLLYNSQMDTSAPSGKKVDATAYQENNTKLPLDEILQLSASSAMGGQDLQKYGTHTAPSTHNTMELHETTSSSQKSKDELHENAPVEKTDNVLGVPREIGEAHVLDSPVIPKENVRKTRDGREKSTQKSQLPKRVTRSMQRASRHMDGNENKIMNGATNCADASENIQRTNLEAKDVNASFKVITDTPIKEEAVAHLKKSMRKSGKNNRQQNDMKEDFVDSAQNLNSFVDKDVDIIMDTIEHLEGDEHTPQESEHAAEVLIDSSNGTIDQGKGTVKNSAQHSINIESMPDGEMEGKIHSRSEGRRLSSAEVIDFPNCFQEICTESQSCKEKQTEIFPPIKRKMQSRPGEKRLSSSLYFPNSSQETETERHGEKEAEMIPSLKGKAQLRSGERTSCNSESDLTNSLEDEKTGTGSQSHEDNEMHGSCRLGLISLTKGKIQSISEERRLSSSEVIDLPDCLQEIGDESQSHAREMRTPCRTGMIPPPLPSPSVPVSKNLISAAFGLKTSRSGRLLVPPLAHWCSQSIVRDKDGGIIAISNGSMNMDTSESGYSKFRPPAETNLQNLQKRLHEAAVEGLKKKTMNKKKKVKRN*

>Azolla_filiculoides0032.g024707

MQTQKLIQISKPFDQSRSQPSFLRLLLPLLLKTKQDCDHLHCDHNNRSQKNKNHSNRAPSRNMDSAGNSSPSTTKNKAHCCHSTPISTHSMAAHTPWCQNRGQNCAQSTDINTGDGIEPPYRSLLRPNQSSQSPFADYILSPQYDHHHQHRQSSWGDISPPARERFASRDSNAVGALGFASHECDDLGSRGFAPSSKHLSRDFDDLGACGFAPTRKGFSCEINDSGSFGFTPGSKIFSRELGPRGFMSHEFNGLGPHGFASRDIDNLGFRGFASPAVEEIYEGICAENYLYILESNLVLEKANYASNLYRAIDNETFETTAIIKRLEERKLLSKNGIVVNLVGTINDCATIANGFPHQMVQLFLSGFPNIWNLLVGMAFPVKQLGATSCSEFLGGGVMPESLTLERDTKHARQSLFSECKEKLPSSCLDFGNRAQADTEKNSLVDGNINIQITPKRRESNLDSETTRDPNEGIMQDNPLHIRMECYNENKEEPVDADIAININTLEEKFLSKNSKLDVDKCEAVTTCADIGAAADSCGVLNQSLKVACIESPIRDACETEQDMGAVASVDGSRKRKVRFRSKNESVAGSSSQTQKSSETHSKRVCQEPFDKLDQEPESTPAQNMDMETQLLSKGISHKKGISKKRGKPKKKVRLCLSKPRRSNKCMISKQKQGMVKNPVSLHEEEKMDTSENPKVASGEQEDCRGEMQQTKNIDDDRNNGNDPASCNEENYMEADQGSQWENKNCREMAHQKVDTKARIESCVSRCSLSTINSPEDECSVKDTPTDISACKVARNRLPPPLPMPTVSVSNQLVSKAFGLKTSRSGRLLVPPLAHWRNQSIVRDSDGGIIAIEDGSADTMGVDTGRFNFKPPLEARARKLQKWLSVTASKCFMQKLNSRANKKKRNVT*

>Marchantia_polymorpha_0034s0011.1.p

MDNIQRSHALGRREGSCKQRATLMAMHGGLHSRRDSGTLGRSEPCCEATFSRNNAGVTANSVATPFHLPNGRHGSHSSFNCRSSCHFHERFSVHDEHAESARRAAKFAQTSCHRRTATQSSCDCGHCQGRDLREDSEAQVSEENVRYRRGASVPDTQAGTGPSHAEEGCGCCHPSGSVLGRNDGGDDFDDRIRALGGTPCFSLSDDLPYRATANAFMSKPSSVSRKGSPDLSRIESMHRALAATKASVKQVILYGWFLKRPSSSNGSASCSAVPQVVIGGYKSAGMSESDYFESGLIVQRLERWRLQTSHGLEVGLMGMINVSKSVANGFPEALVQCFMIGFPSAWSYMISNFGMGSLSDDKTDLPSSSAVPSNATGQAQDPRNYRREVLLNTAPLEGDPVRIILSSSLFIAPDEARMIALRRLSSSNNLQGGVESDATSAPQSVNTQEGLDRARNDRNSCYGAINTGNHSASHDPMPSRLVERAVTQEVLPEGGEEHSVRLGDHNKTRNGSDSCYDAINTRNQVSSHDPRPSRLSERRATQVVLVEGDEIGDGLEDVVSSPTPKDEVSVEGAVAVDLDVSERTPNEEAQPNIDWGEFLGDGNSRDQEYEAGLKLCDEANVTQEGTVHVAVVGAVDTDCNCPGVSYLKLTCASCGCTGIDLDVTTGASQDGESKADLNIKVSSIEFPNRGNEPAAGCAELSEDSSDFTTKRPFQGPVSEEGEEKPSKMLCETHASSLEIRLPATAAIGNDANGSDSGQPIVVGTEAGLMQDSIPCTPLQESKKRSRNSTETPGPTEETILMCNILSHAGPSVLPSQSRTNAPVSEILLNDIDRESVDTAVEPCQEEVIVSAKELAVDESTTRKKRKKSVGAALGRPSIHSVDSDPGLPRDDGLLATEVFSSNSQMTSFVSLRRSRRRSVAESSSRSQSGKESSEVLVVGDEGGKESIHVLKSNREPKGGADVTYFHKDVKEFRVPVVACRNEAGTESNHAMKSDQKLETDEENGCKENKQVVTPGNNKGSRKSRKYYKGRRKKSTKRVGFVEKDTGMNHHSGGSRPEELKSKLHKDSLGTSACGETHIEGGTDHCLQVPREIPAPISQKHQRSSSRRNYRMRRKDHDRETDLQTSGEGQGDAADGGLNQGLEDADKAAKSTKRLKNTAERSFSGSRRASPKVLNRVIQKSQPQRRKNKFSASIVPPPPPAPRVVDKTKVPEAFGLKKSRHGRVLVPRLATWRNQSIKYDMDGSIIAIAEGFEGAEGESGTGCYNFKPPTDKEARTMQRKLQKAAELART*

>Physcomitrella_patens00338G00350

MSCQLQPGAEGSLSESVHFPPTSSFLEHQGYQGDVRMGMAPPLSSFSRAPPPRSPFLHDDGRAGSRYVHSGGFDRQGYDYAVHRHSCGDHLELHSGSLLEHLLLQENQKALRIASLVQSQASGCLSGCLSCGLGEKRLRYKDSEERMVADQRHCCPRSVGEEPSRSVNLGAEADPVECADEWFRQPRPRESVGGVGERSQAKISFFEDELGRSLCQERRTVFPVSRPVLSHSQASHHIHDHQDHLVGAQESSVPIYGFSDGDGRVRQRWPSQRPAYSGNGVEVDGWQRATRSVSRVGSVTATEQDRLSLPDIAAGSLPTVLVGSPHRPVGDASRPTQSARTTSQNLLRMSNSVEPSSRDKPLQEASTQRVVTQGPVSLTGWYIMKVQTADVGRVVETKVAVGGRLVQGGEHVKTSPIVNRLDFHKVVTEDGVEVSLEGSMDLETSTANGFSPGIVQCLCNGFPYMWKQLLRVRPVGMSSSLAVSESVGGLHPVASKCVSEDESQGVVDPNICKTIPKGIPGGSRDCEGNSTVGDVGSDAVAAVDEPSVIQRIASDAVGSKRVETEAVVTAGAETSTAVETEPVDPGGVERCEPPNAFEIEAMILGGGTTVRDEPPNADGKGTVEPEGEGIGRANTSNAIDEGCPLDVPSQQACDPCPTNFDPVPVSNPLDTDSLGKEAESGPTVDTTFVTMRRGRKNKKGPPQPVRSSARLQQRRTKSAESMPNLFKSSGPAQVEQVTVSGEVTRDNSGEQELGEILPDVQRAANGREKITRTSDIIGEVLKDLQEEDMEAANQGFVGIVVDPQPIQVEISVQVQDSGKNLLKDRDDVEVPKLNENDATEDRMEIEENKEMAPAASSDFENEKSEETAQAAPAATFKLVNEEMEEAAPVTPPELIQRKVQSRVDSRYKPSSEALPQASEERVSPELNARMTRSLKRRLRLSAISEPVADVAPVTRSSKRLRRPVPESNSEPSISHQVGDPLVQPVSNPDLSHQVEGPSMEPISNPVTPHHGEVNGFHARVRDLSTRRAKAVNCSNCKKPCSSQEVIKYKNRIASIGATPEHAVERTMKTEQEEYPPLEPQRTESTPGTSSLPKKGETNTSSAKRRSGRPKKTKSANLPAERRTTRRLKLDDSGEGPSEGHNPVPSKQSSRRTSIMPPPLPSPRVQGGLKGKVPEAFGLKTSRSGRLLVPALAYWRSQSIEYDKDGGIIAIFDGFQATPSDTGCFNFTPPQEKHAKKIQEKLCKAAATVKKRK*

>Picea_abies00016756

MEDPFAFQAENSAHSGVTCGCSDEAEYCTQNDRRFHNGFPGNSIQRPYSSPFRTLFNSPVHHRNQLFRRSTRGSNLATGGSDYVHSVSSQKRVNLYKWYLIRVVKDCPQNDMINDEAQVAVGGYTTPDLLDCEKFRTAPIESRIEKCKLRTVDGLDVLLMGTIDELCTNDNGFPFQVIHLFLHGFPFNWNHVINISNGKPSLNMHILPSSKMKVCASKEKLEKPDSECPVVQVDDTVTDKTSLLEPEQSLYKAGSPRESSVGCSKRVRHKPQPARLSKRSICGKLSTGGKPEKLFTVVGSQEKSDILKESETNVSSGAPNKSVKQEAEITPKRRRDKDPVKTEMNVSQRYKRTKESKKQIDSGSSLDEAKLVQSTGTVNIKDCNAIPTIGYFPDSEGDAENRADNQEIEQSSQNKVHLEESIDNCTPVEQTVEIHISVSEFAVTGSKRRRERPSKSNTKRNRNSRNAYSQDTRDIICDEATVMDGIKNNSDAKDMTHCEELHQEITLTSPLDKACKGTDERVHAEEETLADVGVVHFQKIFSSGSAENMPFSSGDSHINNDVIRVGKDVNKSKTKREGKGKSRTGDTNSICDVSKGLVCNDITMKDGLKLNPDDINEIENGDLNPEAKLVQSTGTLNIEDCNANSTIGRFPDSEGDADPRADNQEIEQSSQNKVHLEKLIDNCTPVEQTVEIQIPISEFAVTGSKRRRGRSSKSNTKRNINSRNAYSQESRDILCDEATVMDGIKNNSDVKDMTHCEELHQEITPISPLDKACKGTDKRVHAEEENLADVGLVHFQKNVSSGSAENMPFSSGDSHINNDVIRVRKDVNKSKTKRKGKGKSRTVDANPICDVSKGLVCNEMTMKDGLKLNLDDINEIANGDPNPVSGEIEHDMSAEHDTQIGDNFLKENVDKNKDSLQTQKNDTERTMGTPIGKRKNLHLKIPPPLPPRRVQTTPEEVSKAFGLKTSRSGEISGTCLAILNYM*

>Picea_glauca00028859

MEDPFAFQAENSAHSGVTCGCSDEAEYCTQNDRRFHNGFPGNSIQRPYSSPFRTLFNSPVHHRNQLFRRSTRGSNLATGGSDYVHSVSSQKRVNLYKWYLIRVVKDCPQNDMINDEAQVAVGGYTTPDLLDCEKFRTAPIESRIEKCKLRTVDGLDVLLMGTIDELCTNDNGFPFQVIHLFLHGFPFNWNHVINISNGKPSLNMHILPSSKMKVCASKEKLEKPDSECPVVQVDDTVTDKTSLLEPEQSLYKAGSPRESSVGCSKRVRHKPQPA

>Picea_sitchensis00004746

MEDPFAFQAENSAHSGVTCGCSDEAEYCTQNDRRFHNGFPGNSIQRPYSSPFRTLFNSPVHHRNQLFRRSTRGSNLATGGSDYVHSVSSQKRVNLYKWYLIRVVKDCPQNDMINDEAQVAVGGYTTPDLLDCEKFRTAPIESRIEKCKLRTVDGLDVLLMGTIDELCTNDNGFPFQVIHLFLHGFPFNWNHVINISNGKSSLNMHILPSSKMKVCAKR

>Pinus_pinaster00061713

MEDPFAFQSENSAHSGITCGCSDEAEYCTYNDRRIHNGFTRSSIQRPYTSSFRNLFNSPVRHRNQFFHGSTRDPKPVTGGSDYVHSASSQKQVTLYKWYLIRVVKDCPQSDIIDDEEQVAVGGYTTPDLLDCEKFRTAPIESRLEKCKLRTVDGLDVLLMGTIDEQCTNDNGFPFQVIHLFLHGFPFNWNHFISISNRKPSLNMHILPASNTEDCASKEKVEKPDSECPVVQVDGIETGKSSLLEQEQSLSRTDSPRESSLRRSKRVRHDSQPAKLSKKSICCKLSTEDKPEKIFTVVESQEKSNILKESEANLSSGAPTKSLNFKSSDMDSHVQIVQEDKLTASGKEFVGSQDNPNILKGTEA*

>Pinus_sylvestris00003294

MEDPFAFQSENSAHSGVTCGCSDEAEYCTYNDRRIHNGFTRSSIQRPYTSSFRNLFNSPVRHRNQFFHGSTRDPKLVTGGSDYVHSASSQKQVYLYKWFLIRVVKDCPENGTIGDGEQVAVGGYTASDLLDVDRYITSPIESRIERCKLRTVDGLDILLMGTIDDQCTYENGFPFQVIHLFLHGFPFNWNHIVNICKGQPNSNIAPLLARKTEESVSKECPVVQVDDTGAGKLGSLEPEEYFSRVVSPRESSVRSTKRVRHDSQLYRSSKESKCGKLSTESKPKMHFTGVGSQENSNILKESEANLCLGASSKSIEKEGLMPHLRSSDMDIHGQINHKDKLTDLRSELGKTKPLNFSVSHIVKEPVQNDNKIMKGLLFTKN*

>Pinus_taeda00042871

MNISSQRSEQDSDTENLSEMGNPFRVFQSSNPVNSAGACGCLRQTECCTPKGRQFHSNRFNSPAHSRNQFSLESTPCSKLVTGSGYGIHSTCSQKQVYLYKWFLIRVVKDCPENCTIGDGEQVAVGGYTASDLLDVDRYITSPIESRIEKCKLRTVDGLEILLMGTIDDQCTYENGFPFQVIHLFMHGFPFNWNHIVNICKGQPNSNIAPLLARKTDESVSKECPVVQVDDTGAGKLGSLEPEEYFSRVVSPRESSVRSTKRVRCDSQLYRSSKESKYGKLSTESKPKMHFTGVGSQENSNILKESEANLCLGASSKSIEKEGLMPHPRLSDMDIHGQINHKDKLTDLSSELGKTKPLNFSVSHIVKKPVQNDNKIMKGYNLRRTRGTEELKKQMDSGTSLDEAKFIIENMKFEDTNENTSARFTNSEDYADCRLHNQEVEQSPLTNEHLEQSSNICSRLGEQTQEIQTPISECAKLNTKKTSSKNVFSQASRNLLCDDGTVKDGIESNSDGKDTTYSKKSHQKIIPTRPLHEVSKIYSQFPLDPDFLEFGSKKTRTGTDGKAQAEEDSLADVGFVQLQKNVSFHSAESKSLLPESSHNNKIIGSEKDVNKSKTKRKGRGKRRTTIRHTNPFPAVVEGLVCGEKMKDDVNLKFDDIDATSNGDPNLVACEIDNDTLAEHDTEIGDNCLKENVDGDKGRLQTRKHGTLENTGTDGKAQAEEDVNKSKTKRKGRGKRRTTIRHTNPFPAVLEGLVCGEKMKDDVNLKFDDIDATSNGDPNLVDCEIDNDTLAEHDTEIGDNCLKENVDGDKGRLQTRKHGTLENTGTDGKAQAEEDNLTDVGFVQLQKNVSFHSGQTKPFLPGNSHINNNVMGSEKDVNTSKTKRQGRVLEGFSWGEKMRDDVNLKFDDIDATSNGDPNLVACEIDHDMLGEHDTVTGDNCLKKNVDRDKGSLQTQKHGTLRTVGSPTGKKQYSHARIPPPLPRRVQTTPEEVSKAFGLKTSR

>Pseudotsuga_menziesii00073202

MNISSQWNEQISDPENLSMGDPFGVFQSSNSVNSRVTCGCLRQTDCCTPKGIPFHRNRFNSPMHHGNEFSPGSTPCSKLATCSSDVIHSTSSQKQAYLFKWFLIRVVKDCPENGIIDDGVQIAVGGYRASDLLDSHKFRTAPIESRTEKCKLRTVDGLDVLLMGTIDEQCTYDNGFPFQVIHLFMHGFPFNWNHIVNICKERPSSNLAPLPVSKIEENVCKECPVVQVDDTETGKLSSLEPEENISKVVSKRESSVSCSKRVRCESQISRSSKGSKRGKLSTESKPKMHYTCVGSQENYNISKESEANMCLGASNKSIEKERSITRIRSSDKDIHVQINQQDKLTVLGSELGRKKPLNFSVSHTVKRPVENEKKIMEGYHLRRAKTTEEFKKQIDSGTSLDEAKLITATMNIEDASENIMSDRFTESAADAHHRLDIPEVEQSVMSNEHLKQSKDICSPLEEQTQVIEIPFSEFADSCYKMRRGRISGKSNTKTNRSRNAFSQASSDLLCDDATVKDAIENNSEGKDMTYSKELHQEIIPTRPLHEVSKVHSQFPLDSGFLEFGRKKTRIGTIGKAHTEEDNSADVGFIELQKNVSFHSAQTKSFPPGNSHINNNVTRSEKDVNKSKTKRQGRGKRRATRHTMPVVLEGLVCSEKMVKDNVNLKLDDIDATNGDPHLVGCEVDNDMPAEHETEIGDNCQKENIDRDKGSLQTEKHSTLKTADTPTSKRQHSHANIPPPLPRRVQITPEEVSKAFGLKTSRSGRLLVPPLAHWCNQKIAYDLDGGIIAIFDGISEQKEDNGGYNFKPPTEAEAKKMQRKLCKAANDVIKTPKSGKKIRKG*

>Salvinia_cucullata0025.g009355

MHNHHHPLAQPPSISPLPLYHGSPAPDRCPASADRPFPERISPGCGCNLHHLHASSGSPHFLRTMETAHACCQPGTSELRRPSSAEESRSVFRDPCRSSELSRKFSVAECLFPSHNLSENAYVPRTPQINDFGATPVMQYEPTSTAQFFSWAYKQNNFGMVSEPGHCMMRQQNQVKLYGWYLINSSNNTAEVAVGGSLTPGAVSNGRKETTTITKRLEKCRLLTKDGVEVRLMGLIDDSTTISNGFSIVHLFLSGFPYTWERLLENEFRMYKTKITAGPGPGPSQVLENVNLRDSEISKLSAEHVENSRKNASSLNGNQVDKFILDGQTMDVTQCKQNSSETMPDVKLQSIAGDEQGRVNVGDIGMPMNIMEKTQFTEESHEIIHSAQATKDKPDGMPFLQDDTSGLQKETEKLKELDSPVLRMEHMRKSIDSLEKTDQMMRASRRITRSMQRSSKHMGATENETINGVDKFPQLSVSQEPEIENLTKELRGINASSESITDTFLKQRGIHSMRSTRKPVKDNRCRSSTKESSADDPINIKCSDNKEMEEVVIVDDIEVEETALESKQEAESLTETIEKNCSQRIEEEGEIESQNTNGVGTFPQLGVSHESEAESLTKEQNGITTSPESTTDTSLKATDVHLMRSIRKPVKINRHQSCTKESSVDDPINIKCSDNKGMEEKLVTVDDIEVEQTAVESKQNAENLARTIAKTCSQRIEERAVQDPRNVESFARHDADMEEKTESASIESLLSDVGAMNTTIGNEFREGKETHTSCRAGMIPPPLPCPRVPVSKRLISTAFGLKTSRSGRLLVPPLAHWCSQSLIRDKDGGIIAISDCSMNTSALDDGSLKFKPPAGANLQKLQKRLREAAVEGLKTKTTNKKKKST*

>Salvinia_cucullata0183.g024952

MSQNSHSEGSLLARSFQMGADTELQRHPHRVSYQSRVQYPEEQIVTHIGVHQMRWYLTRPVGCQTSDIGVGGFLSTRAVTSERVETAAIIKRLEERKLLTQSGVVINLIGTIDDCATLTNGFSIQMVHLFLSGFPHVWSLLVGTMSGSVSIEKNSNTSGKVEECQSLFFDSKEESPSNFHAVGNRDQVDAEHGPHLDGSKNMHFTSTTRKFPEIIETHLEPCEKRVLDDDAKCSIEKKGGVDVADMTTGTGIHTETLDDAKLSQGEVLATTARRKMEVAVNVVNGKNDNGKDQLALLENIMEHNEKNNDENESGNDQEAPSSLNECVGTDQKSILDDEIQQIDKAEYGGAETSVSSEENEDKESNENSEIECDLKARMESCISRNSGSTINSPEDACSVRDPPTDASTKKMMKHILPPPLPSPTVSVSNQLVSKAFGLKTSRSGRILVPPLAHWRNQSIIRDSDGGIIAIQDGSEDTMRVDTGIFNFKPPAEARARKLQKWLSVTAADCFTKRLKSLHARSVKPGGPAEPHVGPSSVFSRHLRAMGISLSALPLQPLTKPPLSSPLLSNKDTLSPASAKKRQRGIDSSTDDGTPSPSKSCRRSPHTPEIQERPSRLRPRKLGFDASVQQTDVQSSLPSIAPSENDAIKQIAEKSSGQFSVLNQNSALSPSQHTDKGTAILRTPTRRASPRLLEKLSTKDAATSVVKPRSLNFSPVTPSRKSVDKRLPPPSPNVTKTKVPTTPKRNFLDHVTVDRVKFSVGDDVYVKRTDDDIDEEAEGCLICGKAGKLIECDNCSRGVHLKCTDPPLKKVPDGEWLCPKCELPSKASDSKQNGFQKNGTEALKTARELLLSCKLWAARIERIWKNEDGSLWFQGRWYLIPEETSIGRQPHNLRRELFLTNDIDENEVVSILRKCYVMGPEEFRNSGRDGDDVFLCEYEYDTQFHTFKRISDMDADNHSENDLSEEDEDEDSDGEESLRYGKKSHSQTPPKAANSRGSLLKIGTKSIPHNARIKPATVFERAKAALRLTATPGSLPCREREMTEISSFVTDAVVSGNKGLGHCLYISGVPGTGKTATVMEVMQKLKRRYEEQNANPYRFVKINGLRLTSPEHLYTVLYEALTGHHVGWKKALQLLDERFSNPNPSRRADARPCILFVDELDLLVTRNQSVLYNIFDWPTRPHSRLFVIGIANTIDLPERLLPRIASRMGLQRLSFSPYSHEQLQTVISSRLEAINAFEKQAVEFASRKVAAVSGDARRALELCRRAVEVAESRCTGDSNRGNSAVKGNLVSIKDVEEAIKEMFQAPHIKMMGRCPKQAKIFLVSMVYEQHKTGMAETIFEKVASTYTFLCRNNNEMSSDWDTLLSVGCALGACRLILCEPGSYHRIQKLQLNFPTDDVSFALKQDPEIPWIGKAREYFFLLVVFLELPPAAMVVAKKTKKAQESINNRLALVMKSGKYTLGYKTTLESLRSGKGKLVIISNNCPPLRKSEIEYYAMLSKTGVHHYTGNNVDLGTACGKYFRVCCLSITDPGDSDIIRTMPTE*

>Aegilops_tauschii2Gv20621700.1

MAKKTPSPPPRARSRRGAAPTSPTPAAALSPPFSPAPLRTRLGAAAVAAAAAAAAAASSSPVEHPCVTLCEWWPVRVEGEERKLAVSGFTERNDAFTSAPIAHRYEPLTLQDEGGVVVLLHGSINLLRMRENGFSVQICEQFMIGFPFWWETWDSHMESYPNCFIDPREGSAQFYLEKFQLGNFIQKFGPSFIEDLLNNAKNFPIDHLDAFTESSRFQEYICGNDASTKENSAASDDARPATVANVEIGLTASSISQERDHVDIECNVSLAPAETYTGDETCKEAGNQNDTMHPDAREEDAGSHLFNSDWTCTMCPDHMPNDSEGGNENSVELLAKYPLAIVPPENANCCSEIPGASQSVEPSSM

>Aegilops_tauschii2Gv20622700.1

MRTRSMASKPEPVPSTHGTAARAPAPASVSASTHGTAARAAARASVSASTHGKAARAPARASVSASTHGTAARAPAPASVSASTHGTAARAPAPASVSASTHGTAARAPAPVSVSAPTHGTAAPAPPPASVRAPTYCATVQRCVALLDWWLVRGQGGKIRVAGYIDNVEKNRAGRVFSSGSITVRHADGTLETADNKIVLTRGPLNIEQMHWNGFSREVSEQFRLGFPIQWEKYANSNMKQANEHILSPAKSTEYCVEKFLRSSFANSMEHTLTGFDFRTSKESTGNTDGPGLPNYVKPRIQEPSGNSVGYDNSVSNMAASEGLCNDRMGTPDESFEDPGPGETCNGQASRADNSHEDIQTDASGQRIVTHSADSTLVNNDIDKIEEERGSSKLGNSSVCPGTEHVSEALNQGASPEHGSVQCSRRLRSGKVYGMSNGASLKRRYSKRKTMQHGTLSMKVIPTEETTPPAGPTCHKKVG

>Amaranthus_hypochondriacus008009-RA

MVKISWSDAYRCNDAQRKFHSAPARRRYASREFHSTKIVKRHGCTTLETSDGFIVSLCGFINKSRSLENGYSTEVCRDFIYGFPYNWEHYAASVPEESSQVNLTDSTKCFPSADLDAYNGAQLHELSFSLSDPERELFCNKVYDDLMELSSLIDENEAQPSKPTRSQVDVDVLTSESRNTKVFTFQAGKCECITEVLLEHEDS*

>Amborella_trichopoda_00137.13

MGLTALSSPKFFEKKPVVLKNWYLMKVPSETSGKRLAVGGFHSKTHFHSCPIEERISAFRLRTSDGAILLLEGCIDEQWTLENGFPFEISGLFRIGFPYTWKILVSQFLSEDSSGFSIREVETNNGSSTFLPITHLFGRNLFPSETERFHKQNEEERKACESERQTTKSKSSSVARIQQYEVCLGQANVNVNDSSSSKVRFGSINEETISKGIDNEQPKCAKGFGSLYSNSKVDVVITPEGVKYHGTKLSDWDVPSPSGVSLKVLESNKKASQVAGSKVRRKRTAAILDPELNCSVNLVKKSRTSSSKELKALQTPLSREFKTLYSPSKELKALRIPVSKEFKTLHFPSKESKALRIPVSKELNVLQTPSKGLTALQTPSKESKLSISRSGRIIVRPLAYWCNERIVYGKDGSITSILDGDKYDEQCTRGSKSAPPKRRHTRNQNSSKSKQGERPSERKLTRNEKPSKPKGSPMKLRRR*

>Ananas_comosus021414.1

MGANASPRRSQRGEEEELVASPTPSPSPTPTPTPAPKSPTPFSPSLPPPPKRSFPTTTRFPFSPSPVAVSSAAAAAADASSTDKACVSLFDWWLLGVERECGGKKLAVGGFTTRKQATRIFTSAPIVKRHDAYTLETEDGIIVIIQGMINRERMQSNGFPPEVSKCFLIGFPYNWDQYANEYLQKNPASTNSLGSFFSVSEQHKDSIKGGDFGAKITRNFIDSIKNLFQLTSVDSIVQRSQCLPNEAETHKYKDANKNVDLMPHSVEGNLSAKTIESSEISGMKSVLISSSLNVEDVIPGEQNHSGNLRTDDRNVLDNTSVQKGATIIDAGLLKVRMNVARTLLDSCGSTSRKISTRKAGARTKSQVSSLNREVEPSDLSQENTGSYCVKGESTSLINVEAINSEAPVVEKRSKDEVLSTPRKRKIKQPTSSVIHLEASPVEKKTKEKVLRTTKRTTKTPTSQVYQAPITRQHAKQLSFASPESLNLRRTRSGRIIVPPLAKGSQQIIYDAVCEIFSLL*

>Aquilegia_coerulea1G467000.1.p

MATTPFFPDTVTTPIKTATTATSTYRKTVTLYDWSLIKSDKDYKGRILAVQGLTSEGNSAIRIFSSAPIVKRYDLVTLETADKMRVILKGCINRTHTHENGFSFEVCSRLFLIGFPSIWEHFSYHCEEGFNFENTFGGSYEHDTPKGGDSKRNLSKERSEDAVAVSVSQVTAKNWGPSNTELHKIENTLSENKVGREVVSDTLEDHLVPTAPSVSTLKVEDYYGEKASPYMRSSERAKYLSSRIKAILKSKMKSSNLCDSVPQEYVNDVVTPVEKKSCAPICLSSSEGNIISLEEKNGLDTYARKGALEKSSRKRLFGRISYDETGSEKESTSPVELSRSSRLRK*

>Aquilegia_coerulea1G467200.1.p

MSDKTVTLYDWWLIKSEKDYNGRRLAVGGSTSKENRATRIFSSAPIVKRYNLITLETADGMTVFIKGNFNRTRTQENGISFKVCERFQIGFPCIWEDFSYHCEEDSISSNTSKGLNDCGSLKDGESNGKSSEEKLEDAVVDLASQMTAEVMGPCDIEQQKPDGSLYEYNVGSEIILNTLENGRRDCFTKMSNSNTTLENHLVPTGSDEKDSSTLFKLDKDSSEESLCIFSFEQQIACVSGGNTESKKCTTGSFKPRSSSRLINMRKGTDNSQEIITNSLTCNNETTCHDASKDSLIELGSGVSVCNPAEVNMVSSVEAPSMSLKSFSRKGTFEKSGGDYLTRYDSPNECFTSRGNNNSKKESMGPFEVRRLSKQKKGNANRQSTIRKTTVLVSDVFNISENSLVCKYDTTQHDTLKDSDVSGCNPSEVNMVSSVGTPNQYSRSLSRKGTRKSVQKHPNRRNFLEPSVSACGKTDSKKELRGLSEARRSTRLINTKNGSDSRHGTVGKKSVHVPLGLPSVTITVDEINCVPEGENDLDRTFIGPDETMRKSSLAGIKHPNHGSQSKTRKRLEYITDAKETPIAKNKKKESFVLSPKESNRKRTKSGRLRVPPLAFWHNQRIIYDRNLQIAGVLNGLPAVESLSITGSNSNPVRKKPKFI*

>Brachypodium_distachyon_3g30800.1.p

MAPEPQPGPSRHGTTIQAPPPASVDAPIYRATIQKCVALLDWWLVRGQDDKIRVAGYTERNRAARVFTSDFITMGHADGTLETADHKIVLTRGPLNIKQMHRNGFPYEVSKHFQLGFPAQWEKYANSNMKQSKHQAQSPSKSIDNIWKFLSFMKYNFEETDFNSSKGSTGNTDGTPIQGLSNLSNGTPRFQDSSGPGPGETCSSAQGDNQHQDMHLDAEDAGQVPAKGMSPEFGVVQGSEDSTGRRLRSGKVYGMSTSSSASLKRGRSKRKTIQHDTPNRKILNEETILPVDPTNHGNGGSPVTRSIAAAKLQSPHPFQKGM*

>Brachypodium_distachyon_4g44060.1.p

MASPPEPAIPVLSPLSSRLSAAADSSSVSSSAPVEHPCFTLQNWWLVRVEGEERKIAVSGFTQRGDAFTSAPIAKRHESLVLEDEDGVVVRIDGLMSLCRMRRNGFSLQICESFLIGFPSWWESWDSHFESQPTSSSNSQEDSSQIYLKIFQLGNVVQKSVASFIKNPLHDAKIFRRYVADAFTQCSRFDEYSFDNDTSTKGKTVASNDASEGPAAVANEVDNMEIDLIVSSTSQERGHVDISCNASFAPTEKCTSDETYKEAENQNDSMHPDVTEQEAGNHSVNSDLICNRSRDRMPSDLEDGNTNAGNSTDVALCHLATAQPERVNCCSEIPGALQNIQPLSNQRNPVASLKNQSHPKRTEDISLNQKAVPIEDTSTSIRSHVLSSEKTVGPSKKQRSAQDKLLSPARLRGTRNPISYVHHSPHTRGKAQSLSISTPESLEMTRTKSGRVVVPPLDLGCERILYGNNHLVLGVAPVKLHSPPIKGSKPETPARKRRAR*

>Brachypodium_distachyon_5g25291.1.p

MAAKTPQTPDPPFGARSRPSVPRPPGSAIPVLSPSFSRAPLRSRLAAAADTNAAASSSSVEHPCVTLWDWWLVRVEGEQRKIAVSGFTQRDDAFTSAPIAKCHGPLTLEDEDGVVVLIYGSVSLSRMRENGFSPQICEKFMIGFPYWWESWDSHIESQPTSFSNLQEGSSQFNSEMCQLGKFLEKVEPSFIKNLLNDAKNFPRDYEDAFTECPRFEEYTFDNDISTKEKSGVSNDASEGPAGVANEVDNMEIDLIVSSTSQERGHVDISCNVSFASTEKCTSDETYKEAENQNDTVHPDAREQEAGSRPVNSDLICNRSSYRMPNDLEDGNTNAGNSTDVPLCHLAAVPLGRANCCLEISGALQNIQLLSYQRNPVASLKNQGHLQRTEDISLNQKAVPSEDTSTSICSRVQTQEKTVGPSKKQRSARDILLSPARLPVTRSPMSSKKQIFAHDKLSSPTILPVTRSPMPSKKQRSAQDKLSSPATLPVTRSPISSVYHSPLTRGRAQSSISISTPESLNMKRTKSGRLVVPPLDPGCERILYDNNHLLLGVAPVELHSPLKGNKAGTPAKKKSAR*

>Brachypodium_stacei_03G173200.1.p

MASEPQAGPSNHGTTIQAPPPVSVNAPIYRATIQKCAALLDWWLVRGQDDKIRVAGYTERNRAARVFTSDFITIRHADDTLETADHKIVLTRGPLNIEQMHRNGFPYEVSKHFQLGFPAQWGKYANSNMKQSKQQAQSPSDKSFVKSMDYNLKETDFNSSKGSTGNTDGTPMQGLSNLSNGTPRFQDSSGSGPGETCSSDQGDNQHQDMHLDAEDAGQVRAKGMSPEFGVVQGSEDSTLSRLRSGKVYQKSTSGSASLKQGHSKRKTIQHNTPDRKMILNEETPPPVDPTNHGNGGSPVTPSTAAAKLQSPHPCQKGIFTKRLKTVDRTRET*

>Brachypodium_stacei_05G288200.1.p

MASPPDPAIPVPSPLSSRLGAAAVEHPCFTLQNWWLVRVEGTERKIAVSGFTQRDDAFTSAPIAKRHEPLVLEDEDGVVVRIDGLMSLCRMRQNGFSLQICENFLTGFPSWWESWDSHFESQPTSSSNSQEDSTQMYLKFFQLGSVVKKSVASFIKNPLHDAKDFRRYVADAFAQYSKFDEYSIDNDTSTKEKTAASNDASEGPAAVANAVDNMEIDLIVSSTSQERGHVDISCNASAPTENCTSDETYNEAENQNDTMHPDATEKEAGSHPVNSDLMCNRSPDCMPSDLEDGNTNAGNSTDVALCHLATVQPERANCCSEIAGGLQNIQPLSYERNTAASLKNQGHLKRTEGISLNHKAVPSEDTSTSVSSHVQSLEKAVDPSKKERSARNILLSPTRLPGTRILISYAHDSPLTRRRAQSLSISTPESLKMKRTKSGRVVVPPLDLGCDRILYDNNHLVLGVAPVELHSPLIKGSKPGTPARKRRAR*

>Brachypodium_stacei_09G247600.1.p

MATKTPQTPDPPLRSGSRRSVPPPPPRSVIPVLSPSFSRVPLSSRLAAAASSSSSSVEHPCVTLWNWWLVRVEGEERKIAVSGFTERDDLFTSAPIAKCHGPLALEDEDGVGVLIYGSMNCSRMGENGFSRQICDKFMIGFPYWWESWHSRIESQPTSFSNSQEGSSQSTFQQLGKFLNRESSFIKNLLNDAKNFPRDYEDAFMECPRFEEYTFDDDISTKEKSAVPNDASEGPAAVVNELDNMGIDLIVSSTSQERGHVDNSRTVFAPTENCTSDETYKEAEKQNDTVHPDAREQEAGYQRNPVASLKNQGHLQRTEDISLNQKAVPSEDTSTSIGSHVQSREKAVGPSKEQRSAQDILLSPTRLPVTRSPMPSKKQRSAQDKLSSPTRLPVTRSPMPSKKQISAQDKLSSPTTLPVTRSPLPSKKQRSAQDKLSSPTRSPMPSNKQRSAQDKLSSPTRSPMPSVHHSPLTRGRAQSLISISTPESLNMKRTKSESFALRRGSCGVAFTSQRKQDRDSYEEEKGSLSDFQTQDAAAQPEDADNLTTAACCRCTRNSEGSGCYASLYMQPGV*

>Citrus_sinensis_1g023837m

MDPTKTPRFTKLSSHSFKSVTLHDWWLVRAEGNGLAVGGIASRESQEVRAFCSAAIAKRHNTTTLETADGITITLSNLINRSRTNQNGFPSQVCNRFLHGFPFYWEEYADQCCGQEYTNRGSPFGAANISLPLVSLDDIPATRLRDLLISPRGHADNGALQSEILDVLRPCGCSSVRDKTPSNHQSKAASGRKYRAAVKRPPSRGVVTRSMAKLQTLGSQQNKDSAEFSTQIEGMVTGNGSQGCHAATFSDNDKEPPKSPGNSAVRRSIRLKNQAK*

>Citrus_sinensis_1g045557m

HFDHFPSGELWSMISLINRLLSPKSSTPAPSRASENSNKSDDDDNGDVLASSRFQKTVILYNWWLVKANTDHEGNRLAIAGFTSRELQAKRVFTSAPIAKSYDPFSLETSDGIYIIIRGLINKSRTLENGFSSDVFSHFNLGFHYDWKAYAEKCFMKEVEAALDPIFAADAGTANLDSNAIPVDHKKVKGKRKRNAGDINKSFEYFSENTLASASKSSDSSRHAANHKDTVNTNNHASTNFPQLMKSEKDNVSSLEDNLAVADTAEPSGYCVRGSSKRMANSTSRRKGNTGTAAKSKTEERSKLDGNTMKNVALGTSPVPEGSDTDNANPSNLDPAVSLGDSEVALGIPRTSEKHKENDRSKDGLDYQNINSSPASLPQDFDTNIRCPGEKSSGIASS

>Cucumis_sativus_094610.1

PKTPATCSTSIIPSSLKSIFLYDWWLVKANDGEGLAIGGFASRERSGIRAFYSAAISKRHETTILEATDGIIISISGFINRPRTHENGFPPKVYNHFLLGFPFNWKDYMSSGSIRKSTVEFFKASTSRSNDQGTSHYLEPDLDNLAVTRLRDLCLSTYGESSHGHDLFMKNSNSSCCPTQSFSNEGKNDDVIKDSLHARQEAKKLDIDLQIRRGQGVCTRSMTKLKNTRNRSKESLISDSRKKKKSRK*

>Daucus_carota_025649

MFTPPASSARKHQNNNSNSNSPPTLTPIFNSLLKTNWRKRCNTTRRTPYRFASNNATPLASAPPEQPHFVSTSKNVVLLDWWLTKLQIEGSTSPKAFKFGVGGKAFDGRESTHFFSGEIVKRQDEITLENVEGITIRLDSLLNRFRTLENGFTSQVCDHFFLGFPFDWEEFGAQYFGGQSIHGARSSKGTRTSSDDIDERRCLLSSFDDIPVTRLFDHMMVTSGDYNECSLTRSIFDHILSEYGSSSAELKEEDTDRIPAEDFLVDETKTPLDVSGENNKPLVAGCSSLDKASRDELEHKDDNVILDDVSMGTTNNLLTVHSQSKMDEDVRALSKFVPTRNKTRLETLTSQQQEGLPSNTTTNPDTTSRTSTRQYADTTVTHVASVSQSAIPNAANIKVDRVLDSSSSKNCTTCNHLGKNDSAKRGTMSEMLMNSQELNLFAATLDPGVTAPIPAPEMSRNRIKVPQVDMESRTLNKSGSKKLRRNLNTGSGLLTRSQARGVLTRCRAKLKKLRTDSKDITAEHDVTAEEDAAHFAQTTKISDSCPISEAEKLEKDESRRHVPDKVLEVNNRHLDRPEVRRSGRRRNVVNYRER*

>Elaeis_guineensis_00052_p0043.1

MGAGEKTTPGNQDEPATAVLGSSVTPMARSSSSTTPDPGGSNPTPCTPFTRSSSSSSSEKNTVLLSDWWLVKAERGAGGKRLAVSGFTTRQQAIRVFTSAPIAKRYDAYTLETEDGITVIIQGMINKALAHDNGFPPEVCNRFLVGFPYNWDYYADEYFRRSSTCSSNSSSSAGFGELSKDSADGAGHTSSVCVKESPIGRICDILISSGVTLSSNFTETLKKFFRDPSLNPICQKPLLAVEKCGEHEMDKSDQKHENDKGCATTINHDAEGHVTSHAEEGGHRNDLHAGDVPEENSKLNHDVSYDSLMHESRETDSQPAVEITSQTETGTAMRRETLLTKTRDVEKDNDLVVSVAMDCDNVDHVAACIDGKESAESRLESCKVGMKYLLKDSLPKESNLTNHEHDEELGHDAVKRRHFVEILRVPNVGVSDDLLVSMAQDDKARSKISSQNQVMVHSTILVAQDDADSHPVEGRGMDLNNLKSPNVNITGISKDFSSNEIDDMCRRIFKKIESCKFPAARDSIPQIGSAVDMKSTRASEVKESPTNKHLTDVVKGGRKKENIQKEDQKCAECINSKVCLHFSMPAAQKSANEEKLTGKFSLCGTEELSSTGDVQDDVSLIVLRTRNVPRNSSASSEGKPVKSSTNVKADALSTRMKSSKKKGDDNQMARCSPEVEKIGVSSVPMDHNIVNAANDAKRESRHKHTQAVAKVGKVSLESLPTEHSSEDVNYGIQPCIISTPAATVVMHRQGDRHTNDCIGMNDDNENVTLKFPVKTTGHLANSVKEQSSNDKTLAMTGENIPNGPNDQNEGEGHPLASAALRSLGALTHRPLESSRQHRCPSRGLRGPLVIR*

>Eucalyptus_grandis_I00627.1.p

MDRSGRRQEEGQVHKVGSMMKSLPSQTQVPPSLPPNNRVFLNEWWLVNANGKGLAIGGFAHRDSERVSLFCSAPIVKRHDITTFETADGMAIMVGGLINRLRTLENGFSTKVCNKFLLGFPYDWAEYSTSSCGEYSSVQAGSVKAAASEKTDFWSRSTTNYFEATSLDDVTVPKLRDLFSLGDLDDSFLMKKIYDDVVGMSGSGAEHDGEFLRSCRKSSPSTAAESQTNEAQINDKKSKEDVLCKHKEDIWSTRKIVKKVNHKSGRQLRASYVFRRREGVSTRSMTRFKGL*

>Fragaria_vesca_22410-v2.0.a2-hybrid.t1

MTSTPPPPFSAPRTVSASLKSVLLDDWWLVKAQGNNALAVEGFARSLRPAIRTFSSAAISKRHTATTLETIDGIIVTLCGLLNISRTSQNGFPPEIYDRYLLGFPFDWEEHAAALLGQGSTGKSASARNSSQKYMSLEHSESNQLPFSISDVPATATRDFIMSCVGDSEHLKTILDDILRTLGDNVFGDTTPQINSNMEDSDPVEKVESGCNETLTMAKKVHIDGDRNKSANIHSKGSQKSKKGMYSGRNLTGKISVPRRSSARLKNKNEMQDSSMTLDSELEVMH*

>Glycine_max_09G003000.1.p

MAASKCNSATPLIPVPPKSLIFLHEWWLVKQRKGLAVGGLASVEIADRERVFLSSVIVGREETNVLHSEDGITILFRGFINTSRSSQNGVPFQVCQHFLVGFPHDWKKYSAYSFGDAFGDSSVCSKNICHGLGISSGNHSKQSQSTDNMECECNNTASQLQVGEKGISDVAVESQAGNKKSFMSMFDSNKCTSTSGKGTVKPLTSKEKMDRKEKQQIRQKIVLDASNSCNRMVTRSISKRSHTMPKEDEKATIATVVSPVRRSPRLYSRR*

>Glycine_max_10G043400.3.p

MADSTPTATPSNDTVSSYFQRTVTLYDWWLIKAKNDFQGKRLAVAGVSSRKDEAMRVFVSAAVIKRYDVFSLETADGICVMISGFINEQRTLENGFAAEVFNRFLFGFPPDWESYALDCFREESTTGTDLGSAVPDNVPASSLEILSDGVENSIPTSLASPKEAPGDHEKSFPGNECNVSKEMGGVNVACSSGGKRRSARLHDIKVHQQKKKPASGGSLKNPNNENSTLVALENCDVEGLKSPVTPIQSQSSRQLSTSSGQLVKKSASRISRTLSPKTEGCYKKKRVTVERKVGRPKGKLNKSASAVKNPQEKDLSHLTKGSQQKISTVTPESPSFRKSRSGRLLLPPLEFWRNQIPIYNADHEITEIQDGASLISPCKGFSPSLSRFSNQKRGA*

>Glycine_max_13G131100.2.p

MADSTPTATPSNDIVSSSFRRTVTLYDWWLVIAKNDFQGKRLAVAGVSSRKDEATRVFVSAAVIKRYDVFSLETADGICVIIRGFINEQRTLENGFSAEVFHHFLFGFPPDWERYALDCFKEEPTTDADLGSVVPDNAPASCPKILSDGVEKSIPTCLVSPEEASGDHEMSFPENECNVSKEMGGVHVACSSGGKSHSFKLHNIKVCQQKKQPASECLPNHPDNENSSSVALENCNVERLESPTTPIQPQLWSEYSNDACVENAIPTSLASEEAPGDHEKSFPENESNVSKEINGVNVACSSGGKSRSARLHDIKVYQQKKPASGGSLKHPNNENSTSVALENCDVKGLKSPATPIQSQSSRQLSTSPGQVIKKSASKISRTLSPKTEGCYKKKRVTVETKVVMPKGKLNKSASALKNPREKDLSPLAKGSQQKISTFTPESLSFRKSRSGRLLLPPLEFWRNQIPIYNADHEITEIRDGASLISPCRGFSPSLSRFSNQKRGA*

>Gossypium_raimondii_004G087600.1

MGKRNRERRKSEKPCKSDQLVPVPIGSPTPLANLSLNSVLLHDWWLCMVQPKGLAVGGFECRGRQGQRVLCSAAIAKRHDATTLETADGITVAISGFINTSRTLQNGFSPKVCSHFLFGFPYDWEEYASHSNEQSFCSSTEASITTSLLHSNAMSLPLPSLDNLHVPAARLRDLLMLSAGDSPNSVFDHMLLQKLSTHDSQNAAITTADSNMGNKDPKVCPYSVADGENSNCHKEKVIQNHVDDNNILCSRSKTTVGSRNEDVGILTPTNTRGVITRSMTRLKRRTSERLKRLRLSSSELKEKVLRR*

>Hordeum_vulgare_2Hr1G067110.2

SLRSAPNMARKTRNPPSRARSRRGAPPPPSPSSSTAALALSPPFSPVRFCGRLGAAEAAAADVEHPHVTLWEWWTVRLKGEDRKLAVSSFTEKNDLFTSAPIAQRYESLTLQYEDGVVVLLYGSFNSSRMRENGFSMQICERFMIGFPYWWETWDSHMESYPNLFIHPQEDSIQFYLEKFQLANFIQKFAPFLIKDLNDAKKLPINDLYAFIGGSRFQEHNCGNDVSTKESSAASEDARPAVVAYVEIGLNPSSTSHERDHVNIEGNVSLAPTETYSGDETCKEAGNQNDTMHPDAREDDVGSHLFNSDWTCTMPPNHMPNDSEGGNATNAELLALDPLARVQPKSSNCCSEIAGAFQSVEPLSYQSAPVAPLKNQQYLERNEHITLTQKAVSNKTVPSSIHSDVQSPKKTVGSAKKQRSAKQGLGRPTRLNSAPYARLTRSRVRALSISTPESLKMRRTRSGRVIVPQLDSVRSWVVYDRSKIKL

>Hordeum_vulgare_2Hr1G067130.1

LRSAPNMARKTRNPPSRAKSRRGAPPPPPSSSTAAPTLSPPFSPVWFCGRLGAAEAAAADVEHPHVTLWEWWTVRLKGEDRKLAVSSFTEKNDLFTSAPIAQRYESLTLQYEDGGLWCSFMVHLTHRECVRMGFSMQICERFMIGFPYWWETWDSHMESYPNSFIHPQEDSAQFYLEKFQLGNFIQKFAPFLIKVLNDAKKLPINDLDAFIGGSRFQEHNCGNDVSTKENSAASDDARPAAVANVEIRLNPSSTSHERDHANIEGNVSLAPTETYTGDETCKEAGNQNDTMHPDAREDDAGSHHDLDAFTGGSRFQEHNCGNDVSTKENSAASDDARPAALANVEICLNSSSTSQERDHVNIEGNVSLAPTKTYIGDESCKEAGNQNDTMHPDAREDDAGSHLFNSDWTCTMFPNHMPNDSEGGNATNAELLALDPLARVPPKSSNCCSEIAGAFQSVEPLSYQSAPVASLKNQQYLERNEHITLTQKAVSNKNVPSSIHSDVQSPKKTVGSAKKQRSAK

>Hordeum_vulgare_2Hr1G073590.2

MRTRSMASQPEPGPSTHGTAAPAPPPASVSASTHGTAAPAPPTASASASIHGTAAPARPVHHATLQRCVTLIDWWLLRGQGGKICVTGYIDPIFRRKRSVRLFISSPITVRHAEGTLETADHRFVLTRGPLNIKKMDCNGFPSEVSNQFRLGFPIQWEKYVNSNMKQANEHTLSPEKSTEYSVEEFLCGSFANSLEHTLPGFDFRTSKESTGNADGPGLPNYVKPRVQEPFGNSGDYDNSMSNMAASEGLCNDRVGMPDESFEDPGPGDTCNGQASRADNSHEDTQTDASGKRIVSHSVDSALVSNDIDKIEEEHGPSKLGNNSVCPGTEHVLEVLNKVVSTEHGSVQCPKKLRSGKVYEMSDGASSKRGYSKRKAMQHEALSMKVIPNEETAPPAGPTSRKKGASVAQITALDKLQSHGPGRKGN

>Lactuca_sativa_2_36541.2

MPNLSSASKTPISSSNYNKSVLLHKWWLIKVESESKLGIGGYVNRETLGSRGMRLLGSASASGGKRCNNNNNLNTESKVFYSAAISKRLDNNTLETVDGFTITIIGSIHSSRTLSYGFSPQVCDHFLSGFPFNWEGYFGNFSHEKCERSLPLSFNDLPVTRARDILMSTAQESEACAFTSMILKDILKQTELDHKQDGSSLKIMEEDCGNSRETEAARVSSRAETLPESPSDTFSSLSDIKGEKCNKTRVLGSKWMPLRTHEGYPLRSKRLKKSNETD*

>Manihot_esculenta_08G078100.1.p

MAASASLSNPSNIDNTCSSYFQKTVSLCDWWLIKADEDFQGKRLAVAGVTSREQRAVRVFHSAPITKRHDVFTLETADGIVVIFQGFINRTRTIENGFPSEVFSHFLFGFPPYWEEYADNCFKQAVNSDVNSQNTLDVDEPITDQGLEDVRPTPCKNKDTSDNECYIKVCSLDVSKKYIVDASKDSDLDKDAIELMSSVKNISTYSGVLNDKLIAKLSISQNESHPSIFVCPRDNDDTVDNMEHDAAKNLPPSTSCLTNVVDSMNDNIAVEARLLSYHSIGSSGMIYDTRSGKESQKMCTRKSKNKESVSLESNTIKENASEECSVSNSAYMDNLIVSEFDNTETAGAEGLVLTLCPKSLKKENQNKGKKGGMKSGSKNFSTGSMPRDLEAVLGCIGEGEVICEESLLRSTTAVNSDILKQPSNNFQVTVCSTVQTVAKENEDAPSMVESSPVNAIPSDSGSRKCSTPLRDLKDNKKQKSISGCSVNHKSKELISEVDEHLDGSKLKKRSHSTFQAEKEISQKMLSAARILRNLSGTVGSKEKSEQVSVNKGGSSMKKARRKIIFDAQATPLTGERKEKTCVFSPESLSLKRSRAGRLLLPTLEFWRNQIPVYDADRNITGIQEEFRASRGCNSEPQMRSSKRKGKSPKRH*

>Nelumbo_nucifera_010241510.1

MATTSPTTASNQILDTRTMSSASLRMVCLQDWWLVKSEHNFHGKRLAVGGFATRQRLAGRVFCSTPIIKRHDSFTVETADGVVVTVKGLINNARTCQNGFSTEICSHFLFGFPFDWEDYTSYFFGNECTTDSGGALTNIKGFDETEVHSYNSTNHDMPFHLDKYPVTEIRDCLISITGNSGNSALPSSIFADIIEKCTNSISEKAGELVHMFINGSNTSTSLSSLSAEASRECKKANLRQNEIDNDIFPATNGRLECENVKLEDQGNLNHDGCNETKSETLKDKFTVQSDDVLENAPNDSLFRVKTELVRLSKVELVESRSTSNRNGVRGSGLDVLKRKKRSFSTVESNYLTRSKRLQNREELSENPHGKEGEEQMVCSQVSREVSVLLKWCMKDKVKSNVGTRHGKVD

>Oryza_sativa_01g34610.1

MATQPRAGDGAAAAAAAKEAPAVSYLQACVELDDWWLERVEGEEGKVRVVGSNTTTSRAGRRFTSASIKTRHASGDLETEDGIIIMIARPPNISKMHLNGFPDEVSKHFSLGFPVQWENIINANMAEMNKQPQSPLKSTEYYIEKFLRGNLKYSMGLFSWDGLNIYQGSRSDADRFPSERLSNSSNGRPTVEDPTANTDCNVNFMGTLATSEEFCTGRMDMPEEPRATPSETCGNDQENNQHLCMLMNTCENGNKVQHGTSSVGPSVVPAEKYVRSKAEQDALLVNDSTSHVSSVLGDCATPKCGKSLTHLGTKDALETNEGMNPQFGVPQGSEGSTVRRLRNGKVIVISTSASTKKVYKRARMQDNTFSENVIPNKNVTCPTGLISQENVGSVAVTAAAKLQIHDTPRKGRRGRLRKSEKRKRS*

>Oryza_sativa_04g28040.1

MGTEPSQTPTPPPPPPPASPRRRAPQPATILSPSFSRNPLRATAASASASAGPVPSPSSSSDCEEHPCVELFDWWLKRVEGDDRKVRIAGHTERNHKPHLFTSAPIVKRHKACMLEAEDSIIVLIDGPLDLSQMENNGYSLEVCEKFMTGFPCLWESYNLGSQQSCSYTSISRDGRTKFYLERFQIGNFIDKVGSSFLANLLNNSRSSSGNDADSFEKGSYLSNKKPRFEEYTCDLDISAKEKTTAFNEGSTGSLAVCNKVGNQQIDLVVKSFSKERGHGNIDLSASLTSIEETTRDKTSEDAGNQNEFIHSDAEYQEAGSHLVNSDSIYGMSTESGNQNEFIHADAEHQEVGSHVVNSDSNFDMSTDNMICEMGDGSANAGSAVSQGSKEVLATVLPERANLSPDSCLDNILPISTCNSNNCLENQGFPEIAQHMTLNEEVVPNEDISTSVHSDVESLGNPVGPAEVQRSECDILQGAPRSPKQNVGSAQEQRPEQSMSQGAARSPMIRTPIPDGAPSLRNQHLGSAQEQRSEHFMLKGATRSPMIRTPIPYGHYSPLTRGKAKSSSVSTPESLKLRRTRSGRVVVPTLDPGCQRIVYDRDGLVSGVAGLEFESPPLKGNESRTPESKRRVR*

>Populus_trichocarpa04G07690

MIVIKRNPNSILIMSTISLSSFSTVTSLLSKKPVRVFHSAAIPERYDVFTLQTADGVNVLLQGYINKTLTIENGFSSQVFRHFCFGFPPDWEECGTKLLNSNCESAAEPPVSQNECRPIFLLLPVDDGVNNLKNEDSKNLSPLSSCRVNDVNWVKDSVVVPVKPSGHHVDVALSSEKIGSKKSKTRSFGKLMAKRSSSLERISIKNDASGECSALTDYNVGNITKSNFDQTRSVRTSGVVLAVSLEVSSSLKKKKRENEGNKDGLKSGSDFSMPSLPQGPQVMLRCIENNTT*

>Populus_trichocarpa07G12430

MSTDPKRKGSSSATTPASLTKILHRSLSHLKSRTPNNITPVSSLSLKSVWLYDWWLAKAEGDGLAVSGFTFREFVFGHKLFYKLGSIFLLMITIVRRHYATILESKDGITVTISGFINRDRTRENGFSFQICEHFQLGFPYSWLELATRLGGEESANRGSPPGKSGFDEPKMSSGTSASAASVSFDDLPVTRIRDIATHPLGDSKDCALADILDHFCSNDAKLSPLATSPGSKNPVTVANAVLDETPRKNKRRADRKYKDGGIIPRRVDIVTGEHITPSRGIVTRSMSRRRNFREIREESPSSNPTVSCKASRNYPEKVLPSSAFLSTSETTKNLAGGTMSTSARRDSWATTSTEKIMEVPDVPLVRRSSSRPNIRKDYREK*

>Populus_trichocarpa17G03610

MADVPTHLSINGAKKEKTRPGKMSSEAKEREGFSSSTTPASLTKILIRSLSRLNPRTPSNTVPLSSLSLNSVWLHDWWLVKVEGNGLAVSGFTSREGVGTRLFCSAAIVKRHYTTILEAKDGITVTLSGFINRDRAHENGFSFQICDRFQLGFPYSWEEIAAKLRGEESANGGSPRGKSGLVELNMSSGISTNTASVSFDDIPVTRIRDILMHPLGDPKDCALEDILGSFCSNTMEHTPMLTDPFSNSKSPVTVARKRKRTKADQKHRDGGKITHTDDTVMGECITPRRGVVTRSMSRLRNLAKNNPG*

>Setaria_italica_K96788

MASEPPPGSTGEATRTPPAPSAAAAPRVSYVQQCVVLVDWWLERVEGEEGKIRVAGIASTAQMRHLLLPKGASSSTGNRNVAGRVFRSAAIGRRHDQHAIETEDGYKIQIGRLLNVPRTRDNGFPEKVCKCFEFGFPIQWLKLVNPKMEQQNEQAQSESTADAPRHSVKYWMEEFLSDDLTNLKKYASEENDSYSSAGYTSNTDGPAIQSLSNLPDGNAGNMAASGGLYGGRTNMPGKPLARPRETSCSGQESDQHESMQIDTSEQGLDNHSISSVSVNQNTGSFCPNSKVDDSILATSKIMSVEKESYRRRVGSSKADEDADIQHENMQSCSNEHEIVTLPIDSAIVNENPNSTSSDLEKPGTPKCGKASMNLGSTDALELPTERMTPQFGAVQGSEDSPVRRLRSGKVFGMPSGGLMKSGHKKRKIQHEASSQNMIPNEGDTSTADLTSHENDSSAAGGVTKDKQESHDSHRGISAKKAKKKRESSKLFWNWC

>Solanum_lycopersicum_04g015710.3.1

MAEANNFTLSSSFLKQVYLYDWWLIKVETGDGSKRLGVGGFTAKERPDGRVFHSTTIAKRHDTTTLVTEDGITILLSGFINRCRTLQNGFSSEVCKQFLLGFPYNWEESAAVSFGESTNENAASRISDFSESANASADCTSSSFTLSVDHLSPNVLRDLLISAAGDPEGGMLRKSIFNEIVQKYGNNAFNVDEASSLNQKSGNQVTPQSPSLNGSPSQKKKAKTNRKQEDSCIADAKSGKEKLPEATPKDMPEKRVLDRLLHRSGDDVPIVGENSCLNQKSGNQVSSRGPSLDETPYKKKKTRANLRKEDDKHVPNAQCRKEVLPKCNDESGTDIDKNSSSSSALTRDKASLYKKTKIYLTQEEKRDVHKVSGQGDFGIVNITNSSNGPLTRSRAKMKRVKEQGQEGHRYL*

>Solanum_lycopersicum_09g010550.3.1

MASPSTCQTQRVEEKKEPKSSNSSKSCFQPTVSLKDWWLIRAERDSQGRTLAVAGRTSREGQALRGFTSAPIHKIYDVFNLETIDGICVVLKGFINRSRSEENGFPSEVIEQFLFGFPPQWETFNEKFLGRDSKGKASASYALGFEKPSGCSEKVKDLKNLDQNDYVETTGETIQDHNGRKD

>Sorghum_bicolor_KXG30438

MDSQIWPIPPDTSRSGAISIVHIMKSQKSRSRGMPSQPPPGSTDRSPRTPAPFQPAPDSTGGSPLGSRRRSPPRPAPATSRDMQVTLVEWWLERVESKEGKIGVAGEACVPQMRHQSKEGKIGVAGEAYVPQMSEGAFSSKPNKARRLFRSCAIVKRLGYCTIESEDGYHIRIDGPLNISKTRENGFSEKVCECFKWDFPEQWQGLVNPKMVPDYNEHARSPAETTTAAPSPHEDVDMSIIAGFFSS

>Sorghum_bicolor_OQU89324

MPSQPAPDSTGGSPRIPVASAEAVRRVVDFFKTNGKLRSVTGAAVSSPLLPRGAATPPLAGMPSQPPQGSTGGSPRISAPFQPAPDYTGGSHRTPAPGSIGVVPRIPELSAQAVDRIARKCIILVDWWLERVEGEEGKIRVAGTTFTPRMAEQMRKGASSSNMRMAVRVFRSSAIVKRHDYTSIESEDGYQIEIGHCLNIPKTRENGFSEEVCESFDFGFPDLWQRLVNPKMVPDDEHALSPSETTTGPPSPSVEDYMAKFLSDSFSSKIGFDFTENDFDSGSISSDSKVDGNILAPSKISSVVNEGYRSTVGCGQAKKDANIQQENMPSCSSEHAMVTPKFGKDSVNLGTTDALELPTEGMTPKFGAIRGSEDSIGRRLRSGKVLPIGGPMKKQKKIQQQMVNQGATPAADLTSHENDFSAAEVVVKENLGSDDSCGKVTGQGRIAEGKGKRKRKRVWCRFSYYP

>Spirodela_polyrhiza_6G0070900

MGNSPSRGFRMSEVDKLGRVEDAAVAGTPPSISSASCVSSMEKTVLLHDWWLIKVRDECDRERLAVGGLTAFGKAARIFNSAPVAKRYDAYTLETTDGITVRIQGLINILGPNPFLIDHFFRAVENPRRWHQMKISLTKRHIMELILHRKGEKPHGCLRVTLDRLL*

>Triticum_urartu_13040-P1

MSRLKKRNRAGRVFSSGSITVRHADGTLETADNKIVLTRGPLNIEQMHWNGFSREVSEQFRLGFPIQWEKYANSNMKQANEHTLSPAKSTEYCVEKFLRSSFANSMEHTLTEFDFRTSKESTGNTDGPGLPNYVKPRIQEPSGNSGGYDISVSNMAASEGLCNDRMGMPDESFEDPGPGETCNGQASRADNSHEDIETDASGQRIVTHSMDSTLVNNDIYKIEEEHGSSKLGNSSVCPGTEHVLEALNQGASPENGSVQCSRRLRSGKVYGMSNGASLKRRYSKRKTMQHETLCMKVIPTEETTPPAGPTCHKKLLLGGSDITKACY

>Vitis_vinifera_205s0124g00540.1

MGKGESSNTKATIISASSFLRSVTLHDWWLLKTNANRLAVGGFATRERQGIRVFSSGAIAKRHDATTLETADGITITIVGFLNKSRTHQNGFPSEVCKHFLFGFPYHWEEYAVQCFVGESTKSGVSKKPSGCEEFNLPSTSSENNLLPASLDELPVTRVRDLLMSTLGDSKTLTNSIFSDIVGRVSNLSNMKGNSPTTVDTELDRTPRNHKKAEVEKFEDDNSILDVSDMRTEECKKNICQSSGVMNDSTPSRRVSTRTMTRLKNLKSQLEWNLSSSTSKKQKTRENSEKRLLTNSSNDVLRRSSGNDFKHARKDMKTNGPIKTMDSDLDKFIKHKMAKVEQKRTDNCVSMDTMDIRKEECQKYFCRSEVGMHALSPDRGVATRSMARLKNVKNNP*

>Zea_mays_001145421.1

MPSQPQPSSIGGTSQTPAPFQTAPVFTGGSPRTLATLSAQSVPRVTRRSIALVDWCLERVEGEEGKIRVAGTTYTPQTSPQTRREASSSKGSRKVAGRVFRSSAIVRRHDHFRIMSEDGYLIRIGCLLNIPKTRNNGFSEEVCECFEFGFPIQWHRLVNPKMVPDNKHALSPSETTASAPSHSVGYYMEKFLSDSFSNSNRYSFTENDSYTSVVCTGSRNGLTTQTLSNLPDDNAGNITVSWGLYGLGMDMSEKPWTPPAEACNNRQESDQHESMQIDACKQELVNRSMSSVSVKQSISSISPNSKVDGNRIVPSKIMSVVNESYRSTVGCGQAEEDVGIQQEKKHSCSSEHGMVTLSINCTSSQLGAPGIPKFGKDSVNLWTTDALELSTEGMTPKIGADSTDRRLRSGKILGMPSGGLMNRGHKLKKIKQEASSKQMVNQGATCTVDLTSHENDFSAAEIVVEEKLESHSSCLKGRGGPAKGKRKRERW

>Arabidopsis_halleri_0330s0008.1.p

MAEPNPDDDGSKSYFQKTVVLRDWWLIQCPKEFEGKRFGVAGFEDSVETRAMRVFKSSPIIRALDVFTLLASDGIYITLRGFLNKERVVNNGFTPEISREFIFGFPPCWERFCSSCFVGDSFGTDINTVPSTIDKAFPPILSPCKYSNGNVEDNPAESRDKSSVTETDIAEINDKDGSRARAKKTARRKSLHLSEEEERKLESSYVQNTTNEGDHGSECLSKAKSGDVEKDGCEAINNEDNEWKLDGSELQNRTNDGVHGSEGLIKAKSSDVEKDECEVIDNNVKSPAVGCGIKYTDADNVDKVTSASATGESLTPEQRKGVLGTTASPQCLLKDLDKSSKSEKKGISKKSKNATKESLPSEQRKGRVKVTNASQDPLSKDLINSSKPGKKGKS

>Arabidopsis_lyrata_2G18170.t1

MTTTRSKFQSLSARRFTPLPEPNPSPRTFSKTLPEPNSSPGTNGTFRTPFPLSLITPIKTLKSITLSDWWLKKKSKGLSITGFESNGGSGVRLFSSGTISKRHESTTLEAIDGITISINGFINRSRSLENGVSNEVCNRFRLGFPYDWEDYNVEEEEEKKNVVDISFDDIPVNRYQDLYCLEGCLKDKILDDVVSSLRDLVCQIFDKECEKSRIGGDDGESLVSRVVGVKTRGMLRRREEYEASIGKGVATISGERAVTTSKKKKR*

>Arabidopsis_thaliana_1G58210_new

MTTTRAKSKFQSLSACRFTPLPEPNTSPSTYSKTLPKPNSSPGTDGTFPTPFPLAVITPIKTLKSVTLSDWWLTKKGKDLCIKGFESNGASGVRLFSSGTISKRHESTTLEAIDGITISINGFINRSRCLENGISIEVCNRFRLGFPYDWEDYNEEEEEKKKKNVDISFDDIPVNRYQDLYSLEGCLKDKILDDVVGSLRDLVCQKSDKACEKSRVGDVDDDDDDDDDKSLVSRVVGVKTRGMLRRREEYEASIGKRVATMSGKRVVTVSKKKNRRRSFGW

>Arabis_alpina_G656800.h1.t1

MTNKSKSQSLSTRRTSPRIFSRSLPEPNSSPRTLSRALPKRNFSPGTDGIPRTPFSLGAITPVIGTLKSVSLSDWWLTKKTKENGLGIEGFESKSGSVARRLFSSGAITKRHDSTNLETFDGITVCISGFINQSRTLQNGISLDVCNRFLLGFPYNWKEESVETKKADLGISFDDIPVSRLQDLLVTTCSTCLKSKILDHVVDSLRDLTCPTNNTQKSDKKCEEKLLPMVVGVKTRGMLRQREDNEASSSSIGKLVLSKSKKKR

>Barbarea_vulgaris-Contig491-snap0.32

MATATKSKLQSLSVHRSSPRTRSKSVPESNPSPLTFSRASMKPNSSPGTNSIPPRTPFSLGAITPIGNLKSVTLSDWWLIKKGKEKNLCISGFESKGGSEVRLFSSGTITKRHDSITLEAVDGITICISGFINRSRSLQNGVSNEVCNRFLLGFPYNWNEEEEEDKKKKVEILLDDIPVNRLQDLCFVEGCVKDKILDDVVSSLRDLVCPKSDKKCEKSRIGGDESLDSDDVVDSVRGLVCSKSDKGCEKFRIGGDDESLVSKVVGVKTRGMLRRRQDEDSIGKRRAASNAYSWWWASHIRTKQSKWLEHNLQDMEEKVEYTLKIIDEDGDTFAKRAEMYYRKRPEIVNFVEEAFRSYRALAERYDHLSRELQSANRTIATAFPEHVQFPLEDDSDEDYEGKPHKHLHLIPKGTNIPEVPEIPKKKDFRSQSMMLSRKGPGSLKGAVASALAKREAAIVSSGLTKEEGLMEIDKLQKGILALQTEKEFVRSSYEESYERYWDLENEVAEMQKRVCSLQDEFGLGASIDDSDARTLMASTALISCKDTLARLEEKQKQSVEDAEIEKERIITAKERFDAIRNRFEKPESDDHDDVIRTEDEEVEEADVEEVEEADVVQESSYESEREDSNENLTVVKLAEKIDDLVHRVVSLETNASSHTALVKTLRTETDDLHEHIRGLEEEKASLISDSTDMKQRITVLEDELRNVRKLFQKVEDQNKNLQKQFKEANLTVDDLSGKLQDVKMDEDVEGGGIFQELPVVSGSDDLKSFSKETERSSVEERKNKAIVGKESEDDEGAQEEKPEMKDSFALSETASTCFGTEAEDLVTEDEDGETPNWRQLLPDGMEDREKVLLDEYTSVLRDYREVKRKLGDVEKKNREGFFELALQLRELKNAVAYKDVEIQTLRGKLDTPMKGSPHQVEGNNQLEHDQGQRESVSISPTSNFSVATTPHHQGLDMKRTPGRAKTNEVRVKFADVDDSPRTNIPTVEDKVRADIDAVLEENLEFWLRFSTSVHQIQKYQTTVQDLKSELSKLRIESKQHQESPRSSSSNSAVASEAKPIYRHLREIRTELQLWLENSAVLKDELQGRYASLANIQEEIARVTAHSGGNKVSESEISGYQAAKFHGEILNMKQENKRVSTELQSGLDRVRALKTEAERILNKLEEDLGITSATEARATPSKSSSSGRPRIPLRSFLFGVKLKKNRQQKQSASSLFSCVSPNPGLQKQSSYVKQPGKLPE

>Boechera_retrofracta_RETT00031638

MATTKSKSQSLSARRSSPRTRSKSLPEHNPSPRTHSGALPKPNSSPVTNVILQSPFSLGAITPVRTLKSITLSDWWLTKKGKEKKGLSITGFESKGGSEVRLFSSGTICKRHNSTTLEASDGITICISGFINRSRTLENGVSNEVCHRFLLGFPYNWEDYNEEEEEEKKNVDTDFGVSFDDVPVSRLQDLFSLEGCLKSKILDDVVGSLRDLACSKSDKECDKSRIGDDSLVSRVVGVKTRGMHRRREEYEASIGKRVRHTKRCCREQRAMHIHGGGPATSVQSNPNGSNTIFKVEYTLKIIDEDGDTFAKRAEMYYRKRPEIVNFVEEAFRSYRALAERYDHLSRELQSANRTIATAFPEHVQFPLEDDSDENEDYEGNPRKPPKDLHLIPKGSNIPEVPEIPMKKDFRSQSMMLSRKGPAGLKRTVSSALAKREAAIVSSGLSKEEGLAEIDKLQKGILALQTEKEFVRSSYEQSYERYWDLESEVTEMQKRVCNLQDEFGLGAAIDDNDARTLMASTALSSCKDTLAKLEEKQKQSVEEAEIEKGRIKTAKERFDALRNKFEKSESDDHDEAIKTEVEEEEGDVVQESSYESEREESNENLTVVKLAEKIDDLVHRVVSLETNASSHTALVKTLRSETDELHEHIRGLEEDKASLVSDSTDMKQRITVLEDELRNVRKLFQKVEDQNKNLQNQFKVANRTARDLSGKLQDVKMDEDVEGAGIFQELLVVSGSEDSRHDLKSILTETETRSSLEETKKDATVVRESENGGRSQEEKSEIKDSFALSETASTCFGTEAEDLVTEDEDEETPNWRQLLPDGMEDREKVLLDEYTSVLRDYREVKRKLGEVEKKNREGFFELALQLRELKNAVAYKDVEIQSLRQKLNSPGKDSPHQVEGNNQLEHDQGQRESVSISPTSNFSVSTTPHHQGGDVKRTPGRTKSNEVRVKFADVDDSRRTKIPTVEDKVRADIDAVLEENLEFWLRFSTSVHQIQKYQTTVQDLKSELSKLRIESKQQLESPRSSSNPAVASEAKPIYRHLREIRTELQLWLENSAVLEDELQGRYASLANIQEEIARVTAQSGGSKVSDSEISGYQAAKFHGEILNMKQENKRVSTELQSGLDRVRALKTEVERILSKLEEDLGISSATEARTTPSKSSSSGRPRIPLRSFLFGVKLKKHRQQKQSSSSLFSCVSPSPGLQKPSSYNRPPGKLPE

>Boechera_stricta_30057s0098.1.p

MATTKPKLQSLSARRSSPRTRSKSLSEHNPSLRTHSGALPKPNSSPVTNVILQSPFSLGAITPIRTLKSITLSDWWLTKKGKEKKGLSITGFETKGGSEVRLFSSGTICKRHNSTTLEAIDGITICISGFINRSRTLENGVSNEVCNRFLLGFPYNWEDYNEEEEEEKKNVDTDFGVSFDDVPVSRLQDLFSLEGYLKSKILDDVVGSLRDFACSKSHKECDKSRIGDDSLVSRVVGVKTRGMHRRREEYVASIGKRVRHTKRCCREQRAMHIHGGGPATSVQSNPNGSNTIFKVEYTLKIIDEDGDTFAKRAEMYYRKRPEIVNFVEEAFRSYRALAERYDHLSRELQSANRTIATAFPEHVQFPLEDDSDENEDYEGNPRKPPKHLHLIPKGSNIPEVPEIPMKKDFRSQSMMLSRKGPAGLKRTISSALAKREAAIVSSGLSKEEGLEEIDKLQKGILALQTEKEFVRSSYEQSYERYWDLESEVTEMQKRVCNLQDEFGLGAAIDDSDARTLMASTALSSCKDTLAKLEEKQKQSVEEAEIEKGRIKTAKERFDALRNKFEKSESDHHDEAIKTEVEEEEGDVVQESSYESEREESNENLTVVKLAEKIDDLVHRVVSLETNASSHTALVKTLRSETDELHEHIRGLEEDKASLVSDSTDMKQRITVLEDELRTVRKLFQKVEDQNKNLQNQFKVANRTARDLSGKLQDVKMDEDVEGAGIFQELPVVSGSEDSRDELKSILTETETRSSLEETKKDATVVRESEDGGRSQEEKSEIKDSFALSETASTCFGTEAEDLVTEDEDEETPNWRQLLPDGMEDREKVLLDEYTSVLRDYREVKRKLGEVEKKNREGFFELALQLRELKNAVAYKDVEIQSLRQKLNSPGKDSPHQVEGNNQLEHDQGQRESVSISPTSNFSVSTTPHHQGGDVKRTPGRTKSNEVRVKFADVDDSPRTKIPTVEDKVRADIDAVLEENLEFWLRFSTSVHQIQKYQTTVQDLKSELSKLRIESKQQLESPRSSSNLAVASEAKPIYRHLREIRTELQLWLENSAVLEDELQGRYASLANIQEEIARVTAQSGGSKVSDSEISGYQAAKFHGEILNMKQENKRVSTELQSGLDRVRALKTEVERILSKLEEDLGISSATEARTTPSKSSSSGRPRIPLRSFLFGVKLKKHRQQKQSSSSLFSCVSPSPGLQKPSSYNRPPGKLPE*

>Brassica_cretica_RQL80743.1

MATKSKLQSLSARRSSPRTRSGAVREPISTRAASCSRFVPKPNSDEIPRTPFSFKSITPIGGTLKSVSLSDWWLTKKANEKGLGVAGFESKGGPEVRLFSSATISTRHDSTTLETSDGLTVSISGFINRSRSLQNGLSSEVCNRFLLGFPYHWRDYTEQGFLEEEEEKDYGFSFDDIPVNRLQDVLFTASPRFQDKMLDDAVDSLRDLLRSTTEKPDKECRTPRMDGGDKESLVLSVVGVKTRGMVRRREEGGEASIGERVLRSSKKKRDQ

>Brassica_juncea_A040432

MATKSKLQSLSARRSSPRTRSGAVREPISTPAAPCSRFVPKPNSDEIPPRTPFSFKSITPTTLKSVSLSDWWLTKKANEKGLGVSGFESKGGPEVRLFSSAAISTRHDSTTLETSDGLTVSISGFINRSRSFQNGFSSEDCNRFLLGFPYHWKDYTEERFVEEEKDHCVSFDDIPVNRLQDVLFTASPRFQAKILDDAVDSLRDLLRSSTEKPDKECRTPRMDGGNEESLVLSVVGVKTRGMVRRREEGEASIGERVLRSSKKNKFLLN

>Brassica_juncea_B007540

MATTKSKPHSLSARRSPPRTRSGALPKPLSTPATRSRFLPKPNSDEITPRTPFSSKSITPIIGGTPKSVSLSDWWLTNDGKGLGVAGFESEARLFSSATISTRHDSTTLETSDGITVSVSGFINRSRSLENGFSSEVCNRFLLGFPYHWKDYTEEGFVEEEDEEEEKDYGVSFDDIPVDRFEDVLFTASPRFQDKILGDAIDSLRDLLRSGNEECQEAEKECEERSDKTPIRMDGGDEEEGLVLSDEGVKTRGMLRRREEGEASIGERLHRSSKKKRDQEKR

>Brassica_napus_GSBRNA2T00024219001

MATKSKLQSLSARRSSPRTRSGAVREPISTPAAPCSRFVPKPNSDEIPPRTPFSFKSITPTTLKSVSLSDWWLTKKANETGLGVSGFESKGGPEVRLFSSAAISTRHDSTTLETSDGLTVSISGFINRSRSFQNGFSSEDCNRFLLGFPYHWKDYTEERFVEEEKDHCVSFDDIPVNRLQDVLFTASPRFQAKILDDAVDSLRDLLRSSTEKPDKECRTPRMDGGNEESLVLSVVGVKTRGMVRRREEGEASIGERVLRSSKKNKLIKGSHQNTRTKTISLFLSDQCFVFYASIYAHHHIRPLTKTCCREQRAMRIHGGGPATSVQSNPNGSNTIFKVPSIFHHVFEYTLKIIDEDGDTFARRAEMYYRKRPEIVSFVEEAFRSYRALAERYDHLSRELQSANRTIATAFPEHVQFPLEDDETEDFEGNPRKQPHLHLIPKGSNIPQREAAVVSSGLSKEEGLEEIDNLQKGILALQTEKEFVRSSYEESYERYWDLENEVAEMQKRVCSLQDEFGLGAAIDDSDARTLMASTALSSCKDTLAKLEEKQKQSVEEAEIEKERITTAKERFYALRNKFEKPESDDHDKFIKTEAKVDVVQESSYESEREDSNENLTVVKLAEKIDDLVQKIVSLESNASSHTALVKTLRSETDGLHEHIRGLEEDKAALVSDSTDMKQRIAVLEKELSEVRKLFQKVEDQNKSLQKQFKEANWTADDLSGKLQDVKMDEDVEGAGIFQELPAVSGSEDYLKSITKETEREKDEDEETPNWRQLLPDGMEDREKVLLDDYTSVLRDYRGVKRKLGEVEKKNREGFFELALQLRELKNAVAYKDVEIQSLRQKLGTLEKDSPHQVEGNNQMEHDQGQRESVSISPTSNFSVRVKFADVDDSPRTKIPAVEDKVRADIDAVLEENLEFWLRFSTSVHQIQKFQTTVQDLKSELTKLKIQSKQQQESSRSKHAAASEAKPIYRHLREIRTELQLWLETSAVLKDELQGRFASLANIQEEIGRVTAHSGGSKVSDSEISSYQAAKFHGEILNMKQENKRVSSELQSGLDRVRVLKTDVERILSKLEEDIGISSATEARTTPSKSSSSGKARIPLRSFLFGVKLKKQTKQKQASASLFSCVSPFPAPQQESS*

>Brassica_napus_GSBRNA2T00086338001

MATKSKLQSLSARRSSPRTRSGAVREPISTRAASCSRFVPKPNSDEIPRTPFSFKSITPIGGTLKSVSLSDWWLTKKANEKGLGVTGFESKGGPEVRLFSSATISTRHDSTTLETSDGLTVSISGFINRSRSLQNGLSSEVCNRFLLGFPYHWRDYTEEGFLEEEEEKDYGVSFDDIPVNRLQDVLFTASPRFQDKMLDDAVDSLRDLLRSTTEKPDKECRTPRMDGGDKESLVLSVVGVKTRGMVRRREEGCEASIGERVLRSSKKKRDQ*

>Brassica_nigra_B045991-PA

MATTKSKPHSLSARRSPPRTRSGALPKPLSTPATRSRFLPKPNSDEITPRTPFSFKSITPTIGGTPKSVSLSDWWLTNDGKGLGVAGFESEARLFSSAAISTRHDSTTLETSDGITVSISGFINRSRSLENGFSSEVCNRFLLGFPYHWKDYTEEGFVEEEDEEEEDDYGVSFDDIPVDRFEDVLFTASPRFQDKILGDAIDSLRDLLRGRPDQECQKKPEKECEERSDKTPIRMDGGDEEEGLVLSDEGVKTRGMLRRREEGELSIGERLHRSSKKKKGEKGENFLLRVCFRNIVV

>Brassica_oleracea_1g087890.1

MATKSKLQSLSARRSSPRTRSGAVREPISTRAASCSRLVPKPNSDEIPRTPFSFKSITPIGGTLKSVSLSDWWLTKKANEKGLGVAGFESKGGPEVRLFSSATISTRHDSTTLETSDGLTVSISGFINRSRSLQNGLSSEVCNRFLLGFPYHWRDYTEEGFLEEEEEKDYGVSFDDIPVNRLQDVLFTASPRFQDKMLDDAVDSLRDLLRSTTEKPDKECRTPRMDGGDKESLVLSVVGVKTRGMVRRREEGGEASIGERVLRSSKKKRDQ

>Brassica_rapa_A01g027290.3C

MATKSKLQSLSARRSSPRTRSGAVREPISTPAAPCSRFVPKPNSDEIPPRTPFSFSLKSITPTTTLKSVSLSDWWLTKKANETGLGVSGFESKGGPEVRLFSSATISTRHDSTTLETSDGLTVSISGFINRSRSFQNGFSSEDCNRFLLGFPYHWKDYTEERFVEEEKDHCVSFDDIPVNRLQDVLFTASPRFQAKILDDAVDSLRDLLRSCTEKPDKECRTPRMDGGNEESLVLSVVGVKTRGMVRRREEGEASVGERVLRSSKKNKFLLN*

>Camelina_sativa_XP_010414757.1

MATKSKPQSLSARCSSPPRTRSKPLPETNPSPRTRSKPLPETNPSPRTHPEALPKPNFPPTPRTPASLGAITPIVKTKSVTLSDWWLTRKGKDKEKKALCITGFESDVRLFSSGTILKRHNSVTLESVDGITISISGFINRARSMENGVSEEVCNRFLLGFPFNWEDYNEENVVDEDRGFVVSFNDVPVNRIQDLSFVDGYLKDRILVDVVASLRDMVCPKSDEKKKSVVEDESLVSSAVVVGVKTRAMRRRDEFESSSSGKRPVCTRSTKRKKKLA

>Camelina_sativa_XP_010470274.1

MVATKSKPQSLSARRSSPPRTRSKPLPETNPSPRTLSKPLPETSPSPRTHSKPQPETNPSPRTHPQALPKPDFPPTPRTPASLGAITPIGTRKSVTLSDWWLTRKGKDKKKKALCIIGFESDVRLFSSGTILKRHNSVTLESVDGITISIGGFINRSLSIENGVSEEVCNRFLLGFPFNWEDYNEENVVEEDRGFVVKFDDVPVNRIEDLSFVDGCLKDKILVDVVASLRDLVSCPKSDEKKKKSVAVGEDESLVSSAVVVGVKTRAMRRRDEFESSSGKRPVCTKSTKKKKLA

>Camelina_sativa_XP_019101020.1

MKKKIAMTTKSKPQSLSTRRSPPRTRSKPLPETNPSPRTRSKPQPEPNPSPRTHPKALPKLNFSPFPPTPASLGAITPIGTRKYVTLSEWWLTRKGKDKEKKALCITGFESDVRLFSSGRILKRHNSVTLESVDGITISISGFINRARSIENGVSEEVCNHFLLGFPFNWEDYNEEIVVDEDRGFVVSFDDVPVNRIQDLSFVDGCLKDKILVDVVSSLRDMVCPKSDEKKKSAVVVDDESLVSSAVVVGVKTRAMRRRDEFESCIEKRSVLHKVYKEEEEISLVVSFTLFFNSHFV

>Capsella_rubella_0002s0640.1.p

MTKTTAKSKSQSLSAPRSSPPRTRSKTLPESSNPSPRTHPKPKRNFSPLPRTPFSLGAITPIGTRKDVTLSDWWLTRKGKEKKKGSLCITGFESSKHGSEMRLFSSGTIVKRHNSITLEAIDGITISISGFINRSRSLQNGISNEVT*

>Cardamine_hirsuta_CARHR053810.1

MATKSKIRSLSAHRTSPRSRFKSLPESNPSPFTFTRASSPGTNGTPFSLGAITPIGNLKSVTLSDWWLIKKGNEKTLCISGFESKGGSEVRLFSSGTITKRHDSVTLEAIDGITICISGFIDRSRSLQNGISNKVCNRFLLGFPYDWNEEEKEEEGTVEEKKNVDVSFDDIPVNRLQDLCFLDGCLKDKILDDVVSSLRDLVCPKLDKKCEKSRIGGGDESLDSGMDDESLESQVVGVKTRGMLRQRQDEDSIGKIVRVTTSKKRR

>Cardamine_hirsuta_CARHR210840.1

MSDHQEPNLDGDGANSCSSSSFQRTVVLRDWWLIKCSNEFEGKRFGVAGTETSFESRAMRVFTSSPIIKALDVFTLQASDGLCITLRGFLNKERVFKNGFKPEICREFIFGFPPCWERICNDCFQGDSDINTIDKACSPILSPCKFNRNPAESRDHSTVTETNIAEINSKDGSRAVRRKSLRLQPKSGVNSAKGERKLESSKVQNSTNGGDHGSEGLSKAKSSDVEEDECEAINKEDSYDSKVQNCTSDEDHGGEGLDEAKSSDVEKDECEAINDEAISPGRKQNGADNVDKVTSVSASGESLTSEQRKGKRKGTKTSLHSLSKEINNSSKPGKNRKSKKSDSNVVEPMNHSESEEAEEDLSWGKTKRKIDFDVEVTPEKESKNNVVSTDSLGQKRSRSGRLLVSSLEFWRNEIPVYDTDRNLIQVREGSDTNSKSAPSKGKGSNSLKPRN

>Conringia_planisiliqua_CP11_g33626_DN3_SP0_c

MDRTRAMRVFTSSPIIKAFEVFTLQASDGVCIILRGFLNKERVVQSGFIPEISREFIFGFPPFWEQICNNCFRGVPDATGFNTLPSVIGKASRPILSPCNNTRGNLVDCSAESRDRSIVTEKNTAEINNGRSGGSRAIDKNTASKKSLRLQSKSGGKSSQDEKKLEVSKVQNITNVGDHVREGLNKAKGCNDDVEEDEREAIGNEGNEKKLDESEVHNGTNDEDHGSEGSDKAKNNDVEKDECEVIYNDVTSLADGCGKKHSGADNVDKVTSMSATGELLTSEQGKGELGVTRASPHSGTDSKKLKSKNATKESLTSEQRKGKLKVTKTTVHSKSKDVSNSRKPGRKGKSKISENTLKGDCDVVEPMNHSASKVKEAEENMSGGKINRKIDFDEELMPPLLCNAKVTPDKDAKKQKTNAVSADSLGQKRSRSGRVLVSSLEYWRNQIPVYDMDRNLIQVNEGHETISTPSKGSFFLKAVFDKRGLHCKSNTEFCLALVLDRKGIEFSKAKKMKIKHTASNYFGNY*

>Euclidium_syriacum_0020s0140.1.p

MVDPWEPNLDDDCSSYFQKTVIITDWWLIKCSNEFNGKRFGVAGTEITDSFDQKRAIRVFRSSPIIKAINVFSLETSNGVSIILRGVLNKERVVKSGFNLEISREFIFGFPPLWEQICNKWFEGISLSNDIDTVPSSIIIDKARYSVLSPCKSKNTKRNVEDSVGKNRDKNTVTETNKVKVNDKDGRSVGSRARDENNARRKSFRLRSKPVEEETEFEVLDNEVDDGCGIKHTDDAESMAFEQRKDEPKVTTRAALRKKSENDQSVVVEPMIHTTEDGREVEKSKNVTTEQGKGEVKVTKELDKRSKSGKSKRNVVVTEPMNHSRPEVKQAEKILSMGETKRKIDFDAELTPEKKSDKKQKKSDASSSSIGSFNRSRSGRLLMSPLEFWRNEIPVYDMDRSCIKVKDGDNETPSKAGKRSDSRKPRR*

>Euclidium_syriacum_0048s0039.1.p

MATKSKSESVSMHPPRTRSGYVPEPNSSPRTRSGCVPESKSAPRTDRVARTLFPPDIVTPVGTLKSVVLDEWWLKKGKEKGLCVSGFEIKGGGAIRKFSSGAITKRHDSNTLETIDGITVTLSGFLDRTRSLQNGISFDLCNRFNFGFPYDWNEDEDESVETKNKAFGFDFSFDDIPVRNVNDLLLTSNSSLRSKILDDVVEGLRGFAYGSTQESEKECEKSGMDDNDYNGDDESLVPRVVGAKTRSMLKRVHETSKKKRS*

>Eutrema_heterophyllum_scaffold455_cov155.49_1

MTTKSKSKSLSARRSSPRTRSGAVAEPISSPCIFSRFVPEPNSSPGTDEIPRTSFSFGAVTPVAGTLKSVSLSDWWLTKKTNQKGLCVTGFESKGGSEVRLFSSAAISERHDSTTLETFDGITVSISGFFDRSRTLQNGFSSEVCNRFLLGFPYNWKDHDEEAEEKKQFSVSFDEIPVNRYQDLLFSSYHNEILADVVSSLRDLVCPSTEKSDKKCKKSRMSNDDDDKSVVPRVVGVKTRGMLKRREDSEGEAFSVVERVHTTSNKKNRSREKTKR

>Eutrema_salsugineum_v10023956m

MTKSKSQSLSARRSSPRTRSGAMPEPISSPRTSSRFVPKLNSSPRTNEISRTPFSFGAVTPIAGKLKSVSLSDWWLTKKTNQKGLCVTGFELKGGSEARLFSSGAISRRHDSITLETFDGLTVCISGFINRSRTLQNGFSSEVCNRFLLGFPYNWEDNHDEEAEKKKQFRREEYKKCGKSRMDDDDDDECLVPRVVVVKTRGMLRRREENETRDKKRCCREQRAMLIHGGGLATSVRSNQNGSSTIFRKVEYTLKIIDEDGDTFAKRAEMYYRKRPEIVNFVEEAFRSYRALAERYDHLSRELQSANRTIATAFPEHVQFPLEDDDDNENEDPQKPPKHLHLIPKGSNIPEVPEIPKNEFRSQSMMLSRKGPAGLKRTVSSAQAKREVAIVSSGLSKEEGLEEIDKLQKGILALQTEKEFVRSSYEQSYERYWDLENEVTEMQKRVCSLQDEFGLGAAIDDSEAKTLMASTALSSCKDTLAKLEEKQKQSVEEAEIEKERIDTAKERFDALRNRFNKPEINDHGEVIKTEKEKDVVQESSYESEREDSNENLTVVKLAEKIDDLVQRVVSLETDATSHTALVKTLRSETDDLHEHIRGLEEDKASLVSDSTDMKQRIIILEDELSKVRKLYQKVEGQNKSLQNQFKEANRTAEDLSGKLQGVKMDEDVEGAGIFQELQVVSGSEDSKSISKETERRSSVEEQKKDDIVVKESEGAQEEKPEIKDSFALSETASTCFGTEGEELVTEDEDEETPNWRQLLPDGMEDREKVLLDEYTSVLRDYREVKRKLGEVEKKNREGFFELALQLRELKNAVAYKDVEIHSLRQKLDTHGKDSPHQVEGSNQLEQDQGQRESVSISISPTSNFSVSTTPHHQVGEMKRTKSNEVRVKFADVDDSPRTNIPTVGDKVRADIDAVLEENLEFWLRFSTSVHQIQKYQTTVQDLKSELLKLRIESKQQQESPRSSSNNTSEAKPIYRHLREIRTELQLWLENSAVLKDELQGRYASLANIQEEIARVTAQSGGTKISDSEISGYQAAKFHGEILNMKQENKRVSSELQSGLDRVRALKTDVERILSKLEDDLGISSASEARTTPSRSSSSGRPRIPLRSFLFGVKLKKHRQQKQTASSLFSCVSPSPALQKQSSYVRQPGKLPE*

>Eutrema_yunnanense_scaffold81_cov138.25_1

MKSKSKSLSARRSSPRTRSGAVPEPISSPRTRSGAVPEPISSPRTRSGAVPEPNSSPRTFSRFVPQSNSSARTFSRFVPDSNSSPRTRSRAVPEPNSSPVPDEIPRTPFSFGAVTPVAGTLKSVSLSDWWLTKKTNQKGLCVAGFESKGGSEVRLFSSAAISERHDSTTLETFDGITVCISGFINRSRTLQNGGFSSEVCNRFLLGFPYNWKDHDEEEAEEKKKHFSVSFDEIPVNRLQDLLFTSSACLKNEILDDVVDSLRDLVCASTEKSDKKCKKSGMGNDDDESVVPRVVGVKTRGMLKRREDNEGEASVVERVHTTSKKKRSRVKTKR

>Lepidium_meyenii_scaffold1913.53

MATKNSKSQSLSARRSPPRTRSKSLPEPNSSPCTCFRSSLKPNFSPGTDCILKTPYSVGASTPIGTLKNVSLSDWWLTKKGKEKALSITGFEAKGGTEVRLFSSVAISKRHDSTTLEAIDGITICISGFINRSQTLEKGISNEVCDRFLLGFPYNWKDYNEEEEMEMNNHTDFSISFDDIPVSRLQDLSYLEGSLRKKILVDVVDSLRDLVFTKADKQCEKSRIDVDESSVSMVVGVKTRSMLRKREMNEASIVKRVQTMSKKKKR

>Lepidium_meyenii_scaffold277.39

MTTKNSMSQSLSARRSPPRTRSKSLPEPNSSPRNRFKSLPEPNSSPCTRSKSLPEPNSSPRTRSKLLPEPNSSPRTCSKPLPEPSSLPRTYFRSSLKPNSSPGTDGIFKTPYSLGAVTPIGTLKSVSLSDWWLTKKGKEKALSITGLESKGGNEVRLFSSGAISKRHDSTTLEAIDGITVCISGFINRSLTLQNGISNEVCDRFLLGFPYNWKDYNEEEEMEKNVHTDFSLSFDDIPVSRLHDLSYLEGSLKKKILVDVVDSLRDLVFPKADKECEKSRIDVGESSASMVVGVKTRSMRRQGEANEASIVKRVKTMSKKKKR

>Lepidium_meyenii_scaffold668.120

MADNREPNFEEDVASSSTSFFQKTVVLRGWWLIKCSNEFEGKRFGVAGVEESVETKAMRSFKSSPIIKALDVFTLQASDGIFIILRGFLNKERLLENGFNREIAREFIFGFPPCWERICNNCFKEAPSLIEKASSPILSRCKYSKENLEDNPAESRDKTSLTETNTAEINRIDGSRARDKKTASRKPLPLQSKSGRKSTEDERLGLHLDLNKATNRSIDGDRGRKRGEDRVDKVTTRSASSKSLTSEQRTDELKVARACPHSLSKDSEKTSKPGKQGTSKKSGKILKCDSNHSVNIVKSSGNKRKLNASKVQNPTTSDGDHDSEGLNKRYSATDSVRKQKSYSKPEKKGKSKKSVKNLKSDCNIVTPMNNSGSEANKAEEHLSWEKTKRKIDFDLEVTPDKPKANAASTDSFGQKRSRSGKVLMATLEFWRNQIPVYDRDRNFIQVKEDRETNSTPSSKGKVSDSRKRRK

>Lepidium_meyenii_scaffold96.171

MSQSLSARRSPPRTRSKSLPEPNSSPRTRSKPLPEPNSSPRTRSKPLPEPNSSLRTCFRSSLKPNSSPGIDGILKMPYSLGAFTPIGTLKTVSLSDWWLRKRGNEKALSITGFESKGGTEVRLFSSGAISKRHDSTTLEAIDGITICISGFINRSQTLENGISNEVCYRFRLGFPYNWKDYNGEEEIDKNVHIDFSISFDDIPVNRFQDRSQTLENGISNEVCDRFRLGFPCNWKDYNGEEEEEEIEKNVHTDFSISFDDIPVNRFQDLSFLEGSFKKKMLVDVVDSLRDMVCPKADKECKKSRIDVDESSVSMVVGVKTRGMIRQREANEASIGKIVPRMSKKKKR

>Moringa_oleifera_10005842

MAKRAGRKSLIAKSDPEANQITRTPFTSAAIIPTQSIKYVLLHDWWLSRAEDKGLAVAGFESRGKLGVRIFSSGIIAKRHDAVTLETVDGIRITISGLINRSRTHQNGFPFEVCDIFLFGFPYDWKDYASQCYSVESAERDFQLRKPGCFEFNMVLGSSGNTSGPGSLYDLPAAKIRDLLMSPDGGWENCAESNLLDDVSVHSNMKEESPRTDVNSILCKSAGNHKRAKVDIKLYRGSREILHTKHTATQELQNTGKSVLRADILTPYAGVTTRNMTKLIKEQEANLKNLLVEDENQLVTRGQRLAVRCNEWRGEIEEEETAETPGSGMKHVEKEA

>Moringa_oleifera_10015199

MAPTSAPNCHNNRDVGDSSCFQKTVCLRDWWLIKADKDFEGKRLGIAGVTSARDLQPVRLFTSAPIVKRFDVFTLETADRICIVVNGCINRARTSENGFSSKVFRHFLFGFPPYWEEYAAECLDGESVPDVNKEKMDEGLEDPSPVSTPSKSVEENEEGNCVIDVEINLCENMSQRTTIDAPKFSLLSENDYHPSSTINPVINKDDNAKNEVRKNDGSKDTSEDTPQSLGCEINDVRPLELEIQLPEEPLFCYVAQSSERMGDPSSSGKCQEWILXXXXXXXXXXXSVTTQNNASGESPISWSNKDVLPVVVTNTEGVRSRIHSVSVTQDKGQKSGNITQTGMQNDYSNRLARSPSGCDDVNHAYGLGTILEEGQSCTKEDSSFGERTKRRINFNVHVSFS

>Raphanus_raphanistrum_RrC35_p2

MATKSKLKSVPARRPSPRTRSGAVPEPISTPATRSRSVPKPNSDSIPRTTPFSSKFITPISGGGALLKPVSLSDWWLTKKANNNKGLGVSGFESKGGSKVRLFSSATISTRHDSTTLQTSDGLTVSISGFINRSRSLQNGFSSQVCNRFLLGFPYHWKDYTEEGFVEEDKNGYGVVSFDDIPVNRLQDVLFTASSCFQAKILDDAVDSLRDLLRSSTEKPDKECRTPRTDDGGEESLVLSVKGVETRGMLRRREEGEASIGERLHRSSKKKRDQ

>Raphanus_sativus_018437333.1

MATKSKLKSVSAHRSSPRTRSGAVSEPISTPATRSRSVPKPNSDSIPRTTPFSSKSITPISGGGALLKPVSLSDWWLTKKANNNKGLGVSGFESKGGSKVRLFSSATISTRHDSTTLETSDGLTVSISGFINRSRSLQNGFSPQLCNRFLLGFPYHWKDYTEEGFVEEDKNDYGVVSFDDIPVNRLQDVLFTASSCFQAKILDDSVDSLRDLLRSSTEKKECRTPRMDVGDGESLVVSVKGVETRGMLRRREEGEASIGERLHRSSKKKRDQ

>Schrenkiella_parvula_Tp2g05410

MAATKSTFQSLSACRSSPRTRSGALPQPNSSLRTRSGALPQPNSSLRTRSGAVPHPNSSTPRTCSKSVPELNVSPETEEIPPTPLSFKAITPVVGTLKSVSLSDWWLTKKTKENALGVTGFESKSGSEVRLFSSGAISTRHDSTTLETSDGITVCVSGFINRSRTLQNGFSSEVCNRFLLGFPYHWRDYNEEGFVEEEKKHFTVLFDDIPVNRLQDVLFTSSPCLKSKILDDVVDSLRDLVCPRTVKSDKKCEKSDKRCEKSRTVDESLVPSVVGVKTRGMLRRREENETTIGERVHLTSKKKRSRENNKGTKRCCREQRAMRIHGGGQATSVQSNPNGSNTIFRKVEYTLKIIDEDGDTFAKRAEMYYRKRPEIVNFVEEAFRSYRALAERYDHLSRELQSANRTIATAFPEHVQFPLEDDDDENEDHEGNPRKPPKHLHLIPKGSNIPEVPEIPKKEFRSQSMMLSRKGPAGLKRTVSSALAKREAAIVSSGLSKEEGLEEIDKLQKGILALQTEKEFVRSSYEQSYERYWDLENEVTEMQKRVCSLQDEFGLGAAIDDSKARTLMASTALSSCKDTLAKLEEKQKQSVEEAEIEKERITTAKERFDALRNKFENPESDGHDEVTKTEEKEKEADVVLESSYESEREDSNENLTVVKLAEKIDDLVQRVVSLETNASSHTALVKTLRSETDELHEHIHGLEEDKASLVSDSTVMKERISVLEDELSKVRKLFQKVEDQNKSLQNQFKEANWTAENLSGKLQDVKMDEDVEGAGIFQELPVVSGSEDSRDDLNSISKKTETSSVKERKNDAIVMKESEDTEGAQEEKSETKDSFALSETASTCFGTEGDELVTEDEDEETPNWRQLLPDGMEDREKVLLDEYTSVLRDYREVKRKLGEVEKKNREGFFELALQLRELKNAAAYKDVEIQSLRQKLDTPGKDSLHQVEGNNQLEHDQGQRESVSISPTSNFSVSTTPHHQVGEIKRTSGRTKSNEVRVKFADVDDSPRTKIPTVEDKVRADIDAVLEENLEFWLRFSTSVHQIQKYQTTVQDLKSELSKLRIESKQQQESPRSSSNHAAASEAKPIYRHLREIRTELQLWLENSAVLKDELQGRYASLANIQEEIARVTAQSGGSKVSDSEITGYQAAKFHGEILNMKQENKRVSSELQSGLDRVRALKTDVERILSKLEEDLGISSATEARTTPSKSSSSGRPRIPLRSFLFGVKLKKHKQQKQSSSSLFSCVSPSPALLKQSSYIRQPGKLPE

>Sisymbrium_irio_maker2387.1

MKMATKSKPHPLSARRSSPRTRSGAVPEPISSPRTCSRFVPELNSTDDIPSRTPFSSKAITPIVGTQKSVSLSDWWLTKKANEKGLRVTGFESKGGSEVRQFSSGPISIRHDSTTLETCDGITVCVSGLINRTRTLQNGFASEVCNRFLLGFPYQWKDCSEEGFVEEEEKKRFTVSFDDIPVNWLQDVLFTSSSCFKGDILHHVVDSLRGFVCPMSENSDKECDKSDKECEKSDKNSEKSDKECEKSRMDNDDEEEKSVVPSVVGVKTRGMRRREENETATIGKRVHTLSKKRSREKSKDMEEKVEYTLKIIDEDGDTFAKRAEMYYRKRPEIVNFVEEAFRSYRALAERYDHLSRELQSANRTIATAFPEHVQFPLEDDDDANEDHEGNPRKPHLHLIPKGSNIPEVPEIPKKEFRSQSMMLSRKGPAGLKRTVSSALAKREAAIVSSGLSKEEGLEEIDNLQKGILALQTEKEFVRSSYEQSYERYWDLENEVTEMQKRVCNLQDEFGLGAPIDDSDARTLMASTALSSCKDTLVKLEEKQKQSVEEAEIEKERITTAKQRFDALRNKFEKTESGDHDEFIKTKERQEENVTDVVQGSSYESERDDSNENLTVVKLAEKIDDLVQRVVSLETNASSHTALVKTLRSETDDLHEHIRGLEEDKASLVSDSTDMKQRITVLEDELSKVRKLFQKVEDQNKSLQKQFKEANWTAEDLSGKLKDVKMDEDVEGAGIFQELPVVSSEDSRDDLKSISKETGTRSSVEGRKKDANAVKEIEDIEGAQEEKPEIKDSFALSETASTCFGTEGEELVTEDEDEETPNWRQLLPDGMEDREKVLLDEYTSVLRDYREVKRKLGDVEKKNREGFFELALQLRELKNAVAYKDVEIQSLRQKLDTPGKDSPHQVEGNNQLEHDQGQRESVSISPTSNFSVSTTPHHQVGEMKRTPGRTKSNEVRVKFADVDDSPRTKIPTVEDKVRADIDAVLEENLEFWLRFSTSVHQIQKYQTTVQDLKSELSKLRIESKQQQESPRSSSNHAAASEAKPIYRHLREIRTELQLWLENSAVLKDELQGRYASLANIQEEIARVTAQSGGSKVSDSEISGYQAAKFHGEILNMKQENKRVSSELQSGLDRVRTLKTDVERILSKLEEDLGISSATEARTTPSKSSSSGRPRIPLRSFLFGVKLKKHRQQKQSASSLFSCVSPSPALQKQSSYVRPGKLPE*

>Tarenaya_hassleriana_010524136.1

MMGIPARWKTTMAKKRKSSSRPAPRSPCSRYQTRPDPISSPAHDTIARTPFSAGASTAVQSLNSVFLSDWWLRRVQGKGLSVAGFEAKGGSGVRLFSSGAISKRHDSTTLETVEGITVSISGLINRLRTLQNGFSFEVCSHFLMGFPWYWEHYATLSCGQEETGEKHDQTNEGRPASLKGESASCSTVSFDDIPVNRFYELLMSPPNHSKDCLGNKALDEVLGRLRHCISQGSPVKEQSDKRFETSADSINSEAGEEEEGADHDKKMATGMGVKTRGMMRLRRTIKEEGDLPFGCAPGKRFQNARYMASVEARNASRNSSVEKNGQSLSSKASSAYDDYSELTI

>Thlaspi_arvense_17942

MATKSKPLFLSARRSSPPVPELISSPRTRSGAVPELISSPRTFSKFVPEPNSSPRTDGIPRTPFSSGAVTPVVGALKSVSLSDWWLTKKANEKALGVTGFESKGGYEVRLFSSGTISIRHDSTTLETSDGIRVCISGFINRSRTLQNGFSSEVCNRFLLGFPYSWKDHDEDETVEEKKHFGISYEDLLFTSSSCLKSEILDDVVNSLRDLVCPRTEKSNEECEKSRMITEDDDDESVVPSVVGVKTRGMLRRSEENEASIGKRMLQRAASNAYSWWWASHIRTKQSKWLEHNLQDMEEKVEYTLKIIDEDGDTFAKRAEMYYRKRPEIVNFVEEAFRSYRALAERYDHLSRELQSANRTIATAFPEHVQFPLEDDDNENVDFEGNAPKHLHLIPKGSNIPEVPEIPKKEFRSQSMMLSRKGPAGLKRTVSSALAKREAAIVSSGLSKEEGLEEIDKLQKGILALQTEKEFVRSSYEQSYERYWDLENEVTEMQKRVCSLQDEFGLGAAIDDSEARTLMASTALSSCKDTLAKLEEKQKQSVEDAEIEKERIITAKERFDALRNKFEKLESDDHDEVFKTEEEEEADVVQESSYESEREDSNESLTVVKLAEKIDDLVQRVVSLETNASSHTALVKTLRSETDDLHEHIRGLEEDKASLVSDSTDMKQRITVLEDELSQVRKLFQKVEDQNKNLQKQFKEANTTADDLSGKLQDVKMDEDVEGAGIFQELPVISGSETKEAETRSSVEERKKDDTVVKESEEIEGGEEEKPEIKDSFALSETASTCFGTEGEEMVTEDEDDETPNWRQLLPDGMEDREKVLLDEYTSVLRDYREVKRKLGEVEKKNREGFFELAIQLRELKNAVAYKDVEIQTLRQKLDTHEKESPNQVEGSNQLDHDQGQRESVSISPSSNFSVSTTPHHQVGDMKRTPGRIKPNEVRVKFADVDDSPRTKIPTVEDKVRADIDSVLEENLEFWLRFSTSVHQIQKYQTTVQDLKSELSKLRIESKQRLDSPRSSSSNTAVASEAKPIYRHLREIRTELQLWLENSAVLKDELQGRYASLANIQEEIARVTSQSGGSKVSDSEISGYQAAKFHGEILNMKQENKRVSSELQSGLDRVRALKTEAERILSKLEEDLGISSATEARTTPSKSSSSGRPRIPLRSFLFGVKLKKHSKQKQSASSLFSCVSPSPALQKQSSYVKQPGRLPE
