## Supplementary Table.1 for "Structural Basis of βKNL2 Centromeric Targeting Mechanism and Its Role in Plant-Specific Kinetochore Assembly"

**Supplementary_table_1: List of all primers used in the study**

| Primers used for site directed mutagenesis | |
| --- | --- |
| βKNL2ΔN-For | ATGGTCACATTATCCGATTGGTGGCTAAC |
| βKNL2ΔN-Rev | GAAGCCTGCTTTTTTGTACAAAGTTGGC |
| βKNL2ΔSANTA-For | AATGAAGAAGAAGAAGAGAAGAAGAAGAAGAATGTTG |
| βKNL2ΔSANTA-Rev | GGATTTTAGGGTTTTGATTGGAGTGATGACAG |
| βKNL2ΔC-For | GACCCAGCTTTCTTGTACAAAGTTGGC |
| βKNL2ΔC-Rev | GTAATCTTCCCAATCATAAGGAAACCCTAAACG |
| βKNL2(C)-For | ATGAATGAAGAAGAAGAAGAGAAGAAGAAG |
| βKNL2ΔMotif I-For | TGTTTGAAAGATAAGATTTTGGACGATG |
| βKNL2ΔMotif I-Rev | CTTCTTCTTCTTCTCTTCTTCTTCTTC |
| βKNL2ΔMotif II-For | AAATCTGATAAGGCATGTGAGAAATCAAG |
| βKNL2ΔMotif II-Rev | ACAACCCTCAAGAGAATAAAGATCCTG |
| βKNL2ΔMotif III-For | TATGAAGCTTCTATTGGGAAAAGAGTTG |
| βKNL2ΔMotif III-Rev | ATCATCATCATCATCATCATCATCATCAAC |
| βKNL2ΔMotif III new-Rev | AACTCTTTTCCCAATAGAAGCTTCATAC |
| Primers used for EMSA | |
| KNL2pf3aF | GGTTGCGATCGCATGGATTACAAGGATGACGATGACAAGGCAGCCGGTATGACGACGACGAGGGCGAAGTCCAA |
| KNL2pf3aR | GTGTGTTTAAACTTACCAACCGAAACTTCTTC |
| pAL1_f | GGTTAGTGTTTTGGAGTCGAATATG |
| pAL1_r | TTGCTTCTCAAAGATTTCATGGT |
| Primers used for genotyping and sequencing of constructs, colonies and transformants | |
| attB1 | GGGGACAAGTTTGTACAAAAAAGCAGGCT |
| attB2 | GGGGACCACTTTGTACAAGAAAGCTGGGT |
| Primers used for BIFC vectors construction | |
| BamHI 35S-For | ATGGATCCGTAAAACGACGGCCAGTGCCTAGC |
| MCS-35S-Rev | CTCGCATATCTCATTAAAGCAGTCTAGAACTAGTGAATTCGCGAAAGCTCGAGAGAGATAG |
| MCS-tocs-For | CTATCTCTCTCGAGCTTTCGCGAATTCACTAGTTCTAGACTGCTTTAATGAGATATGCGAG |
| PstI-tocs-Rev | TACTGCAGCTGCTGAGCCTCGACATGTTGTCG |
| Eco-VenN-For | TAGAATTCATGGTGAGCAAGGGCGAGGAGC |
| Spe-VenN-cmyc-Rev | TAACTAGTAAGATCCTCCTCAGAAATCAACTTTTGCTCCTCGATGTTGTGGCGGATC |
| Eco-VenC-For | TAGAATTCATGGACAAGCAGAAGAACGGCA |
| Spe-VenC-HA-Rev | TAACTAGTAGCGTAATCTGGAACATCGTATGGGTACTTGTACAGCTCGTCCATGCCGAGA |
| SpeI-cmyc-VenN-For | TAACTAGTATGGAGCAAAAGTTGATTTCTGAGGAGGATCTTATGGTGAGCAAGGGCGAGG |
| VenN-XbaI-Rev | TATCTAGACTACTCGATGTTGTGGCGGATCTTG |
| SpeI-HA-VenC-For | TAACTAGTATGTACCCATACGATGTTCCAGATTACGCTGACAAGCAGAAGAACGGCATC |
| VenC-XbaI-Rev | TATCTAGATTACTTGTACAGCTCGTCCATGCCG |
| SpeI-attR1-For | ATACTAGTTCAACAAGTTTGTACAAAAAAGCTG |
| SpeI-attR2-Rev | ATACTAGTAACCACTTTGTACAAGAAAGCTGAA |
